# Discovery of molecular glues that bind FKBP12 and novel targets using DNA-barcoded libraries

**DOI:** 10.1101/2024.12.09.627499

**Authors:** Trevor A. Zandi, Michael J. Romanowski, Jessica S. Viscomi, Zher Yin Tan, Bingqi Tong, Simone Bonazzi, Frédéric J. Zécri, Stuart L. Schreiber, Gregory A. Michaud

## Abstract

Molecular glues are small molecules that engage their target and presenter proteins cooperatively. FKBP12 molecular glues (FK506 and rapamycin) were discovered several decades ago and have been used clinically, but our understanding of the breadth of FKBP12 molecular glues and targets has yet to be fully revealed. To identify novel targets of FKBP12 molecular glues, we constructed and screened a multi-million-member non-macrocyclic FKBP12-ligand DNA-encoded library using 25 structurally distinct proteins. Synthesis and validation of selected hits in biophysical and cell-based assays confirm FKBP12-dependent molecular-glue recruitment to bromodomain-containing protein 9 (BRD9) and quinoid dihydropteridine reductase (QDPR). One glue showed no measurable binding to QDPR alone but had appreciable binding in the presence of FKBP12 using either purified proteins or intact cells. The sites of recruitment were characterized with mutational analysis, competition-based methods and X-ray crystallography. The results of this study confirm that FKBP12-binding DELs can yield novel molecular glues generating highly selective FKBP12-target protein interactions.

## INTRODUCTION

Over 30 years ago, immunophilins (FKBP12 and CypA) were the first proteins demonstrated to function as “presenter proteins”. When bound to immunosuppressant molecular glues (FK506, rapamycin, and cyclosporin) they recruit and inhibit protein phosphatase calcineurin (FKBP12-FK506 and CypA-cyclosporine) and protein serine/threonine kinase mTOR (FKBP12-rapamycin)^1^. The presenters bind their respective small molecules and generate a modified surface that produces a gain-of-function enabling FKBP12 or CypA molecular glue binary complexes to bind their target proteins with high affinity and specificity. Despite these early insights, there remains a gap in our understanding of the potential of FKBP molecular glues, even as the chemically induced proximity (CIP) modality has been widely adopted to rewire biological systems by recruiting E3 ligases^2^, deubiquitinases^3^, kinases^4^, phosphatases^5^, transcription factors/effectors^6^, and other functional classes of proteins to various targets^7^. Chemical inducers of proximity generally fall into two classes of compounds: heterobifunctional molecules and molecular glues. Heterobifunctional molecules are engineered by connecting two monovalent binders with a linker to generate a bivalent chemical inducer of proximity that can still bind both proteins independently. Most emergent CIPs described above have used the heterobifunctional strategy due to the conceptual simplicity in the case where ligands against desired proteins are available. Molecular glues can bind one (FK506 binding FKBP12), or neither protein-of-interest (indisulam^8^), in binary fashion with high affinity and rely on cooperativity to form a ternary complex. This feature facilitates the existence of lower molecular-mass compounds, restricts targeting to tissues expressing the presenter protein, and avoids the hook effect, wherein target-only binding of the CIP inhibits formation of the ternary complex at higher concentrations. However, the current lack of systematic design principles has limited the use of molecular glues widely.

Molecular glues and their targets are often discovered during mechanism of action deconvolution; for example, the natural products FK506/rapamycin/cyclosporin, immunomodulatory drugs (IMiDs)^9–11^, and anticancer agent indusulam^8^. Despite this reliance on serendipity, there has been an explosion of basic and translational research in recent years to exploit the induced proximity modality with the aim of developing chemical probes and medicines. The relatively straightforward design of heterobifunctional compounds has hastened the generation of targeted CIPs, but it is often unclear how to develop a CIP against a target that is considered unligandable.

Molecular glues hold promise for targeting novel protein surfaces. An illuminating example came from Shigdel et al. who reported that a macrocyclic compound (WDB002) could recruit FKBP12 to a short coiled-coil domain within centrosomal protein 250 (CEP250), a structural feature long thought to be unligandable due to its flat, pocketless surface^12^. In addition, studies in IMiDs have revealed that CRBN can be recruited to multiple targets containing a β-hairpin structural motif, but in some rare instances to targets that are devoid of a β-hairpin structural degron (Solomon, J. Beyond the beta turn: A novel structural motif bound by a CRBN glue degrader. Presented at Chemical Biology in the HUB, Cambridge, MA, USA, September 19th, 2024.) ^13^. Overall, there is a shortage of strategies to select presenter protein-target pairs that do not fit into the small set of proteins known to be readily recruited by known classes of molecular glues, naturally leaving the option of screening large compound libraries with the hope of identifying novel CIPs.

FK506, rapamycin, and WDB002 bind FKBP12 with a shared motif and target distinct protein surfaces through their variable “effector” motifs, which suggests there is an opportunity for expansion of FKBP12 molecular glue discovery. To address this challenge, we explored the recently developed chemical inducer of proximity DNA-encoded library (CIP-DEL) screening strategy. Work from Mason et al. and Liu et al. identified VHL CIPs, including molecular glues, for the bromodomain-containing proteins BRD4 and BRD9^14,15^. We built an FKBP-biased CIP-DEL of 2.1 million unique compounds (Tan et al. Just accepted) and screened 25 diverse protein targets, including both bromodomains and unrelated proteins, to determine if novel FKBP12 molecular glues could be identified against disparate structural classes of proteins. Our results reveal that CIP-DEL screens can uncover novel FKBP molecular glues for both bromodomains and non-bromodomain-containing targets.

## RESULTS

### Design of FKBP-CIP-DEL

We designed an FKBP12-directed multi-million-membered DNA-encoded library by leveraging protein structure information of known FKBP12 binders. We identified two exit vectors for DNA-barcode and diversity elements. We chose a triazine-based diversity element due to its ability to incorporate a wide variety of readily available amine building blocks. In addition, we used a derivative of the minimal FKBP binder, SLF, with an amide in place of the typical ester due to increased chemical stability during steps in the DEL synthesis^16^. We synthesized a 3.2 million barcoded compound library, FKBP-CIP-DEL, containing an FKBP12-binding element connected through a variety of linkers (n=44) to a triazine library containing two points (290 BB1 and 249 BB2) of diversification (Figure 1a). The linkers consisted of rigid, cyclic moieties to mitigate the entropic penalty of ternary formation (Figure S1). Due to the symmetry inherent in triazine moieties, reversing the order of addition of the triazine building blocks resulted in identical compounds with distinct DNA-barcodes. This generated a subset of the library of approximately 1.1 million compounds that is double encoded, yielding a mechanism for gaining confidence in positives relating to these compounds.

**Figure 1.**
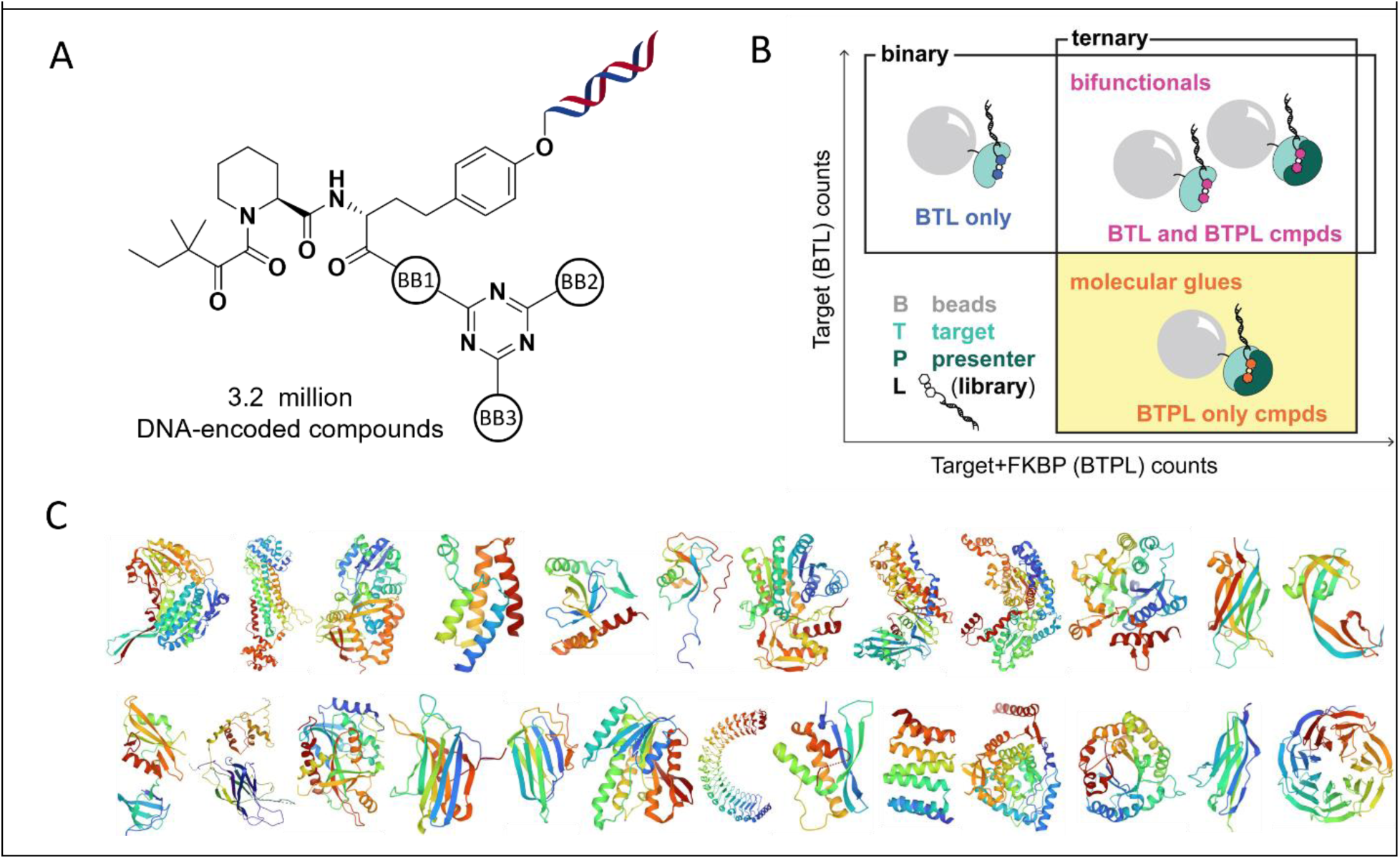
**FKBP-CIP-DEL library screening**. A. An FKBP scaffold-based DNA encoded library (3.2 million total compounds, 2.1 million unique compounds) was generated using three points of diversification, BB1, BB2, and BB3. B. Compounds in the upper left quadrant are interpreted as target binding compounds (with negative cooperativity when bound to FKBP12), compounds in the upper right are interpreted as bifunctional target binding compounds (with no negative or positive cooperativity when bound to FKBP12) and the bottom right quadrant are compounds predicted to be molecular glues, only interacting with target in the presence of FKBP12 (positive cooperativity). C. Twenty-five proteins were produced and screened, all of which correspond to proteins for which the protein coordinates yielded high resolution protein structures in the protein data bank (PDB). Protein structures images were retrieved from their respective PDB entry on the Research Collaboratory for Structural Bioinformatics (RCSB) website. In order from left to right and top down the proteins are: ALDH7A1, ASL, ASS1, BRD9, BRPF1, DCX, GALE, GCK, GPI1, GYG1, ITCH, KDM4C, LCK, PIK3R1, PNP, PRKCG, PTGES3, QDPR, SHOC2, SNX14, STIP1, TH, TPI, TTN, and WDR5.

### FKBP-CIP-DEL Screens

To identify FKBP12 molecular glues, we screened the FKBP-CIP-DEL against immobilized targets without and with soluble FKBP12 to find compounds that display FKBP12-dependent enrichment, i.e. molecular glues, but do not show enrichment in the absence of FKBP12 (Figure 1b). To our knowledge, there is currently no method to predict potential targets for any of the peptidyl-prolyl isomerases-based molecular glues. Therefore, we chose a diverse set of well-behaved targets by leveraging protein constructs that had been used to generate structures deposited in the Protein Data Bank (PDB) (Figure 1c/S2). 25 proteins were purified, and their quality was assessed by SDS-PAGE and analytical size-exclusion chromatography to identify monodisperse and pure proteins. Each of these proteins was screened in duplicate with FKBP-CIP-DEL in the presence of 0, 10 or 200 µM soluble FKBP12 and subjected to multiplexed next-generation sequencing to identify candidate molecular glues.

For most targets, a broad cloud of hits was observed with small numbers of outliers for either apparent target-only binders or molecular glues (Figure S3). To prioritize the molecular glue hits for follow-up, we looked for compounds that 1) appeared as outliers only in the presence of FKBP12, 2) repeated across the duplicates, 3) revealed hints of SAR and 4) were not observed as hits across the other targets, and 5) were double encoded within the library. The two target proteins that were prioritized for hit synthesis and follow-up were bromodomain-containing protein 9 (BRD9) and quinoid dihydropteridine reductase (QDPR).

### BRD9 screening results and validation

BRD9, a 597-amino-acid protein, contains a bromodomain that binds acetylated, butylated, and crotonylated-lysine residues present in histones, and is important for chromatin remodeling and the regulation of transcription as a component of the BAF complex^17,18^. Our screen of the BRD9 bromodomain (residues 135-242) contained promising positives that were only enriched in the presence of FKBP12 (Figure 2a). The observed barcodes with the greatest degree of FKBP12-dependence inferred both a 4-aminopiperidine linker (BB1) and an *N*-ethylpiperazine-1-carboxamide (BB2/BB3). Interestingly, a DEL singleton without structurally related hits, was also observed with FKBP12-dependent enrichment, in this case containing an ethylene-bridged, 4-aminopiperidine linker. We selected two compounds in the *N*-ethylpiperazine-1-carboxamide series, Compound 1 and Compound 2 and the singleton Compound 3 (Figure 2a), for off-DNA synthesis to assess two distinct chemotypes for apparent FKBP12-BRD9 molecular glue activity.

**Figure 2.**
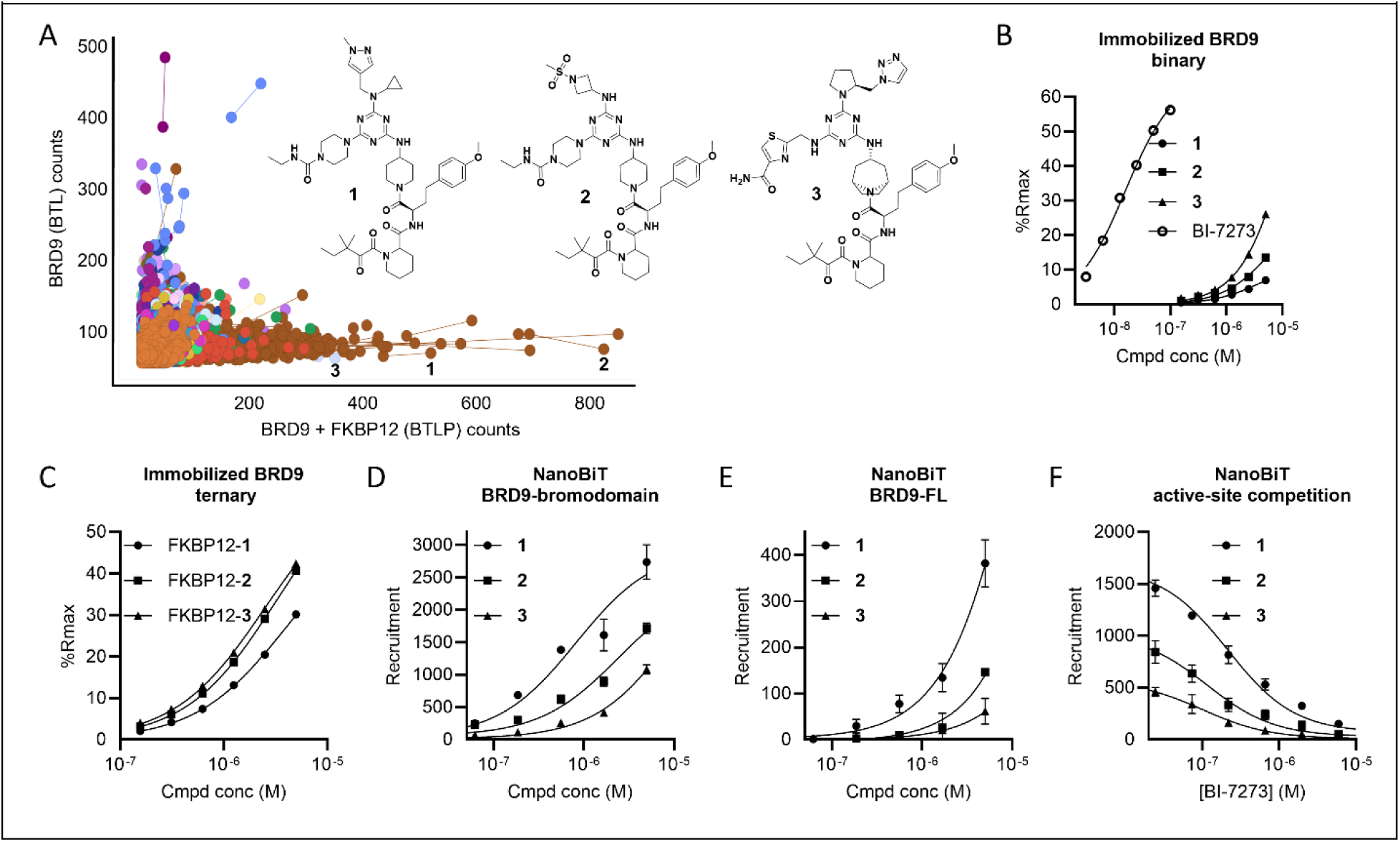
BRD9 screening results and validation. A. FKBP-CIP-DEL count data for BRD9 alone, BTL (y-axis) versus BRD9 in the presence of 200 μM FKBP12, BTLP (x-axis). DNA-barcoded compounds are colored by the linker. Compounds synthesized off-DNA are drawn below. B. Biotinylated-BRD9 SPR with titrations of BI-7273 or compounds 1-3. C. Ternary complex SPR for biotinylated-BRD9 in the presence of a fixed concentration of 10 µM FKBP12 and titrations of compounds 1-3. D. BRD9 (bromodomain) NanoBiT recruitment with titrations of compounds 1-3 E. BRD9 (full-length) NanoBiT recruitment with titrations of compounds 1-3 F. BRD9 (bromodomain) NanoBiT recruitment in the presence of a fixed concentration of 5 µM compounds 1-3 and a titration of BI-7273.

The three compounds were tested for binary affinity to BRD9 alone and compared to the well-characterized BRD9 bromodomain binder BI-7273. By surface plasmon resonance (SPR)^19^, BI-7273 has a measured KD of 16 nM with BRD9 (Figure S4b). While some BRD9 binding was observed for Compound 3, we were unable to determine a KD for any FKBP-CIP-DEL hit against BRD9 (Figure 2b/Figure S4b), in part due to poor solubility of the compounds. There are two distinct affinities in any ternary complex, that of the target-compound binary complex toward the presenter, and that of presenter-compound binary complex toward the target. We chose to measure only the affinity of the FKBP12-compound complex toward BRD9, as we were unable to generate a 1:1 binary complex of BRD9-compound due to low affinity and solubility limitations. Thus, we assessed FKBP12-compound-BRD9 ternary complex formation by SPR by adding 10µM soluble FKBP12 to our compound titration series and observed robust ternary formation to immobilized BRD9 in the low micromolar affinity range for compounds 1, 2 and 3 (Figure 2c/Figure S4b). To assess recruitment of FKBP12 to BRD9 by compounds 1-3 in cells, we used a NanoBiT split-luciferase reporter assay. Each of the three compounds induced recruitment of the BRD9 bromodomain, with Compound 1 showing the most potent activity, EC50/Amax of 0.82 μM/2736 (Figure 2d/Figure S4b). Compounds 1-3 were also tested for recruitment of full-length BRD9 with Compound 1 again displaying the greatest cellular activity, EC50/Amax of >5 μM/382 (Figure 2e/Figure S4b). To address the site of recruitment of FKBP12 to BRD9, recruitment of FKBP12 to BRD9 was assessed by NanoBiT in the presence of BI-7273. In the presence of fixed concentrations of compounds 1-3, a BI-7273 titration resulted in dose-responsive inhibition of recruitment with IC50s of 100-200 nM (Figure 2f). BI-7273 inhibition of FKBP12-compound recruitment to BRD9 was also confirmed by SPR (Figure S4c). Therefore, the competition data were consistent with FKBP12 recruitment overlapping with the acyl-lysine-binding pocket of BRD9.

### QDPR screening results and validation

The protein quinone dihydropteridine reductase (QDPR) is an enzyme that catalyzes the reduction of quinoid dihydrobiopterin using NADH.^20^ Screening as described above yielded two structurally related FKBP12-dependent hits in our DEL-screen (Figure 3a). We selected the higher enriched DNA-encoded positive for off-DNA synthesis (Compound 4). Compound 4 exists as a mixture of four diastereomers in the DEL and we thus chose to synthesize and test it as a mixture of diastereomers for off-DNA validation. We attempted to measure the binary affinity of Compound 4 with immobilized QDPR by SPR. The QDPR SPR assay was validated by measuring an affinity of 63 nM with the QDPR co-factor NADH (Figure 3b/Figure S5a), consistent with prior reports^21^. When we titrated Compound 4 up to 10 µM, we did not observe binding of Compound 4 alone to QDPR (Figure 3b/Figure S5a). We then assessed if Compound 4 would induce recruitment of FKBP12 to immobilized QDPR by SPR. Indeed, we observed ternary complex formation with an apparent affinity/KD of 3 µM (Figure 3c/Figure S5a). To address the region of recruitment on QDPR, we assessed ternary complex formation in the presence of NADH and observed no FKBP12-Compound 4 recruitment to QDPR (Figure 3c). These data are consistent with the FKBP12-Compound 4 complex recruiting to a region at or near the NADH-binding site of QDPR.

**Figure 3.**
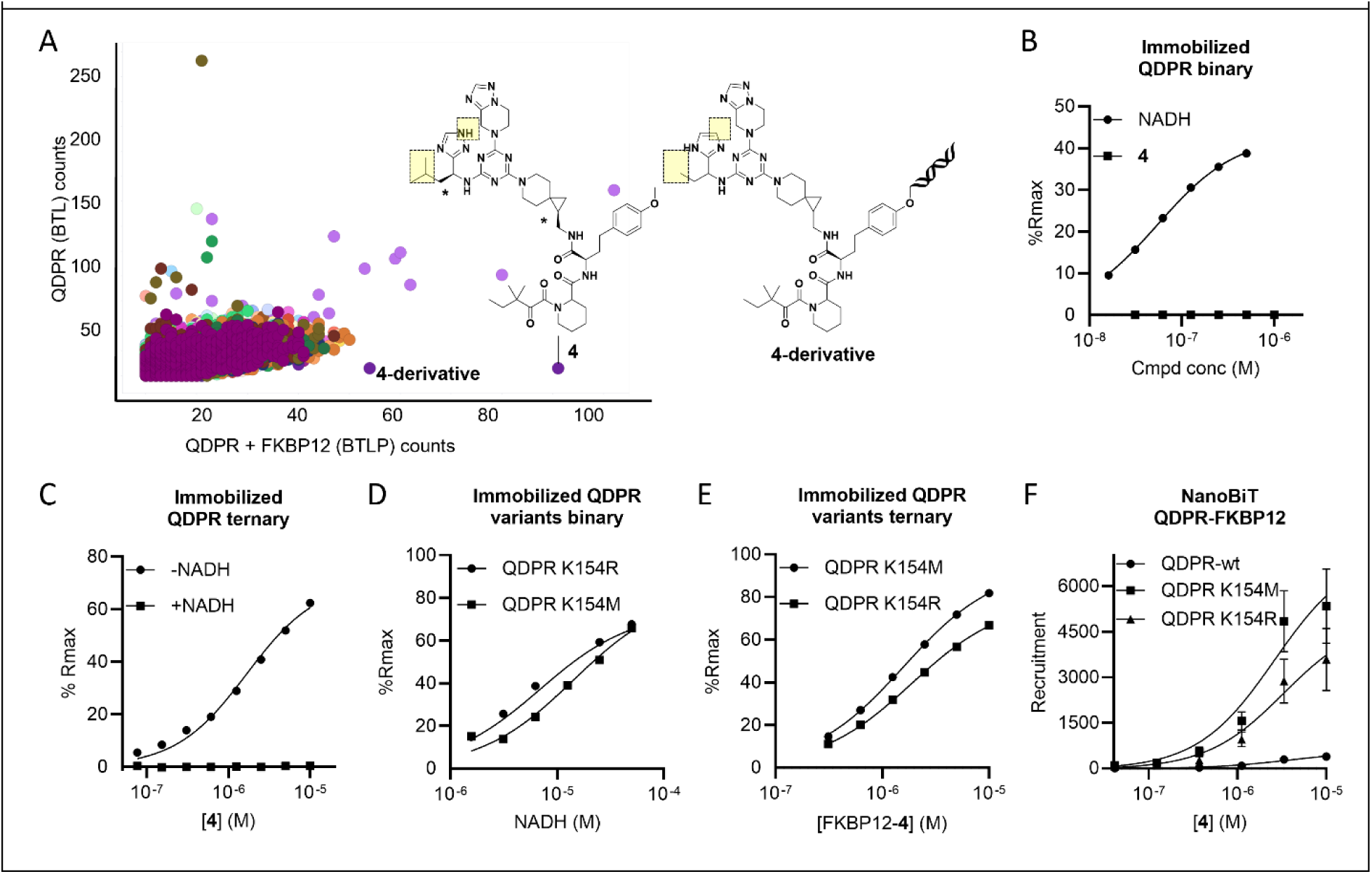
QDPR screening results and validation. A. FKBP-CIP-DEL count data for QDPR, BTL (y-axis) versus QDPR in the presence of 200 μM FKBP12, BTLP (x-axis). Compound 4 is depicted below the graph with stereochemistry assigned by X-ray crystallography. The stereocenters designated with asterisks remained racemic in all assays, both on-DNA and off-DNA. A related DNA-barcoded positive that was not made off-DNA is designated compound 4-derivative. Key differences in = 4 and 4-derivative are highlighted in yellow. B. Biotinylated-QDPR SPR with titrations of NADH or Compound 4. C. Ternary complex SPR for biotinylated-QDPR in the presence of fixed FKBP12 and titrations of Compound 4 in the absence or presence of 50 μM NADH. D. Binary complex SPR for biotinylated-QDPR,single mutants, K154M or K154R, with a titration of NADH. E. Ternary complex SPR for biotinylated-QDPR mutants, K154M and K154R, in the presence of a fixed concentration of 10 µM FKBP12 and titrations of Compound 4. F. QDPR NanoBiT recruitment with wild-type (wt) and two single mutants, K154M and K154R.

To address if Compound 4 induces recruitment of FKBP12 to QDPR in cells, as in the case of BRD9, we performed a NanoBiT split-luciferase reporter assay. Given that cellular NADH was measured in HEK293 cell as 9 µM^18^, our data predicted that the high intracellular NADH concentration would competitively inhibit recruitment of FKBP12 by Compound 4 by at least 90%. Therefore, we introduced two amino acid substitutions in the QDPR active site, Lys154->Met and Lys154->Arg, expressed and purified biotinylated forms of these single variants and measured their affinity for NADH. As expected, we demonstrated that the KD for QDPR(Lys154->Met) and QDPR(Lys154->Arg) toward NADH was approximately 200-fold higher than wild-type (Figure 3d/Figure S5a). However, the ternary KD of both protein variants with FKBP12 Compound 4 was ~2 µM, approximately unchanged from wild-type QDPR (Figure 3e/Figure S5a), demonstrating that Lys154 is critical for binding NADH but not the FKBP12-Compound 4 complex. In the NanoBiT assay, wildtype QDPR gave a dose-responsive increase in signal, but only about 4-fold over background, perhaps due to competition by endogenous cellular NADH (Figure 3f). Supporting this hypothesis was the observation that both QDPR Lys154 variants (Lys154->Met and Lys154->Arg) yielded signal increases of 14-fold and 9-fold over wild-type in cellular recruitment, respectively (Figure 3f). The data support the notion that the FKBP12-Compound 4 complex binds QDPR at a region that partially overlaps with NADH.

Lastly, the proximity inducers for BRD9 (Compounds 1-3) and QDPR (Compound 4) were tested in the QDPR and BRD9 NanoBiT assays, respectively. We did not observe robust recruitment for either the QDPR CIPs in the BRD9 assay (Figure S6a) or the BRD9 CIPs in the QDPR assay (Figure S6b), demonstrating the selectivity and robustness of the CIP-DEL approach to identify target-selective, FKBP12-dependent molecular glues.

### Protein structure analysis

To gain structural insights into recruitment of FKBP12 to the targets by CIPs, we solved the X-ray structure of the FKBP12-Compound 1-BRD9 ternary complex at 2.01-Å resolution (Figure 4a/Table 2). The structure revealed a 1:1:1 complex of the three components. In agreement with the competition experiments with the acyl-lysine site-binding compound, BI-7273, we observed that the FKBP12-ligand complex was bound in the same pocket of BRD9 (Figure 4a/S7a/b). The interactions of the small molecule are confined primarily to the pocket (Figure 4c), while FKBP12 makes contacts with amino acids around the pocket’s periphery (Figure 4a/b/S7c). The ZA-loop of the bromodomain is the major source of protein-protein interactions, highlighted by hydrophobic interactions between Met175 and Ile176 of BRD9 with Ile90 of the FKBP12 80s loop and additional protein–protein interactions between the N-terminal portion of the ZA loop and the 40’s loop of FKBP12^22,23^ (Figure 4b/S7c). The interactions of Compound 1 and BRD9 closely mimic those of a histone peptide’s butyryl-lysine with BRD9^18^ (Figure 4c/d/S7b). The carbonyl of the urea group in Compound 1 forms a hydrogen bond with Asn216 Nδ2, a common theme across bromodomain-ligand interactions, as that hydrogen bond is critical for binding of acylated histone lysines (Figure 4d). The *N*-ethyl group of Compound 1 penetrates further into the pocket, overlaying the distal two carbons in a histone-derived butyryl-Lys residue (Figure 4d). The residue Phe161, which forms the bottom of the BRD9 pocket, restricts the size of lysine modifications that can be accommodated in the pocket and shifts approximately 1 Å downward as depicted, expanding the pocket relative to acetyl-Lys-bound BRD9, in line with the structure of butyryl-Lys (Figure S7b).

**Table 2.**
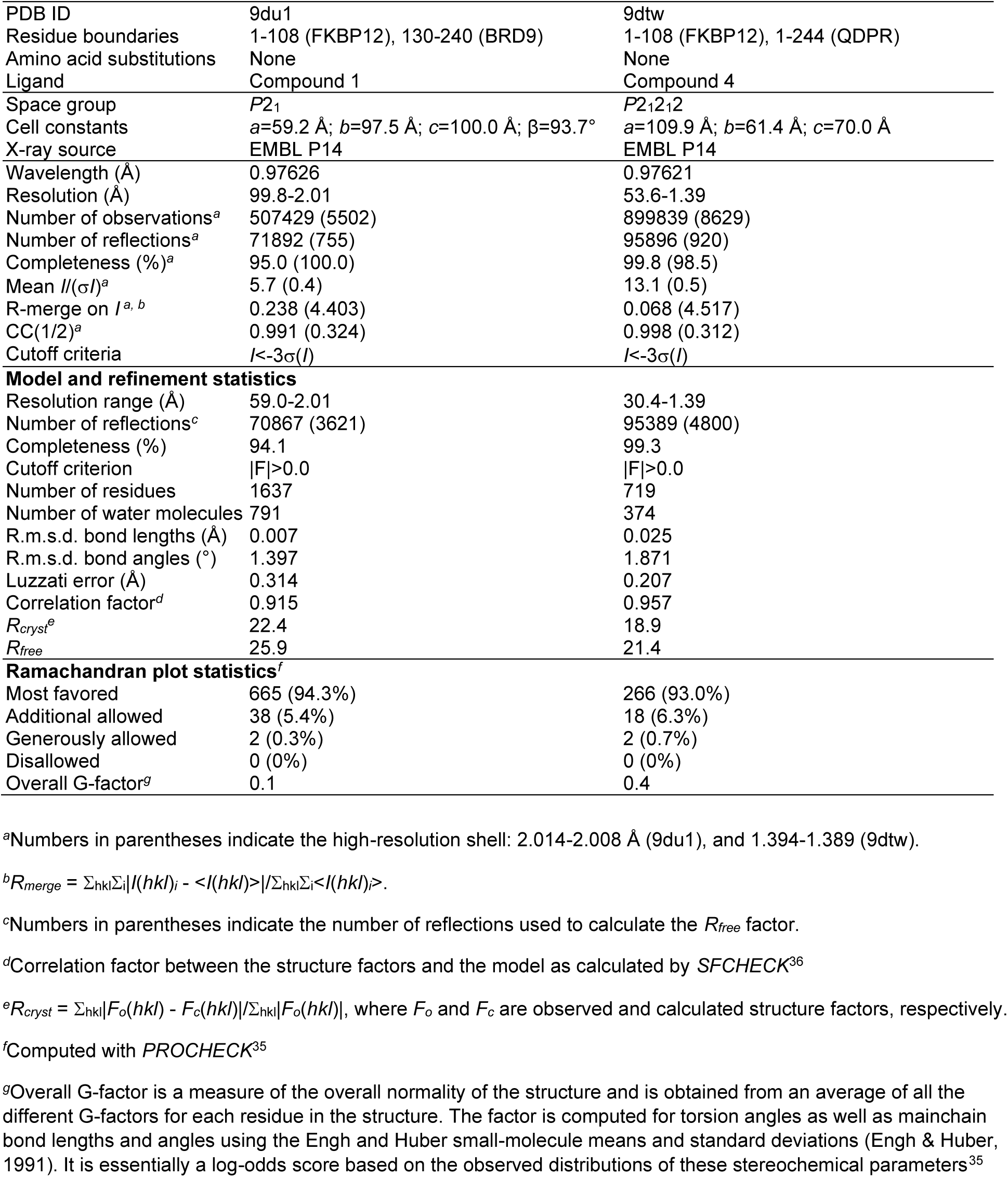
Crystallographic Data and Refinement Statistics.

**Figure 4.**
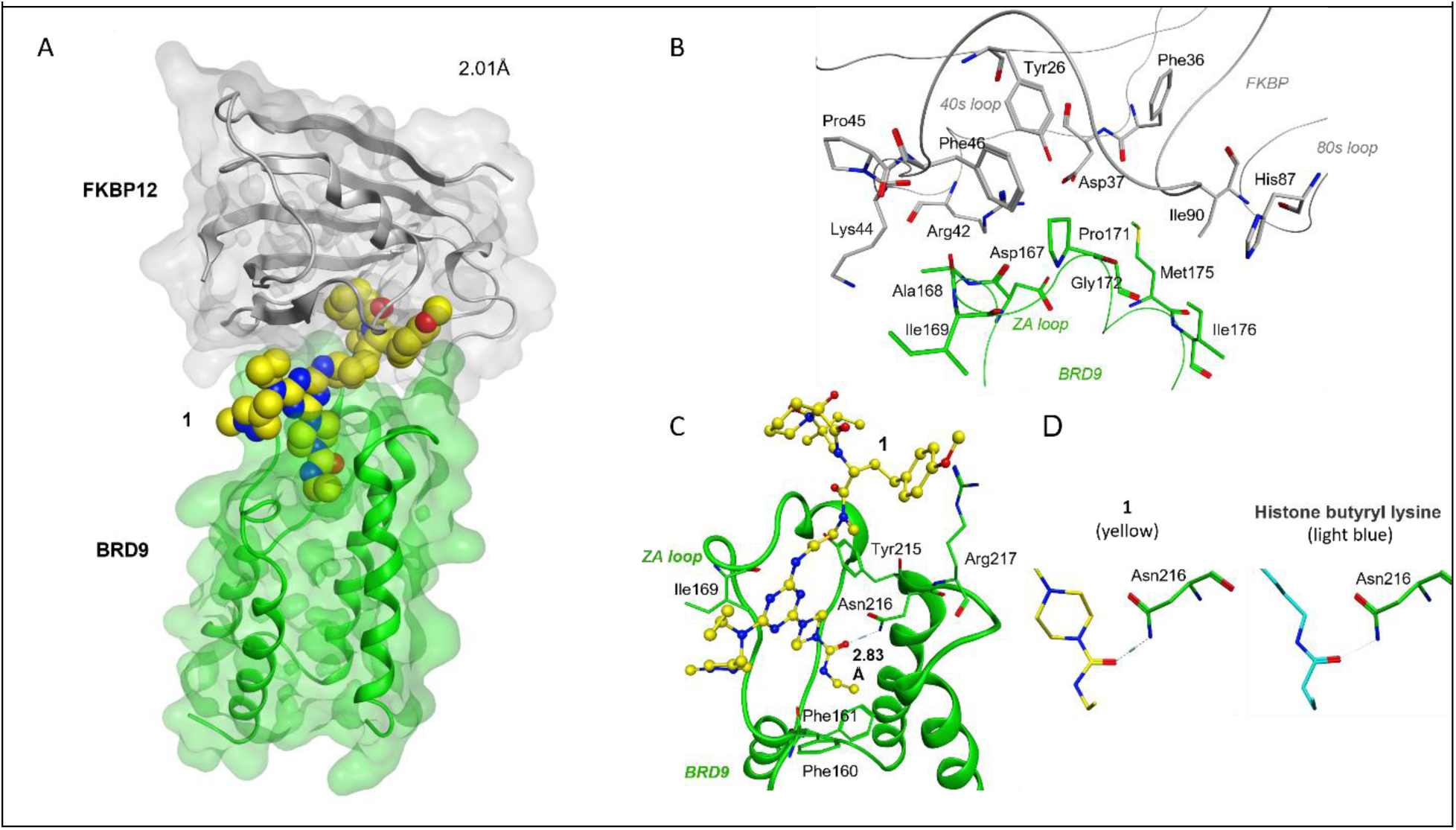
X-ray structure of FKBP12-cpd 1-BRD9 ternary complex. A. 2.01-Å resolution structure of FKBP12-Compound1-BRD9 ternary complex; FKBP12 in grey, Compound 1 in yellow and BRD9 in green;B. FKBP12-BRD9 protein-protein interactions within 4.5 Å, with FKBP12 (highlighting 40s and 80s loops) and BRD9 the ZA loop; C. Compound 1-BRD9 interactions within 3.5 Å; 1 shown as ball-and-stick with BRD9 ZA and BC loops labeled; note Asn216 H-bonding to Compound 1; D. X-ray structures of compound 1 (yellow), left, and butyryl-lysine (light blue, PDB ID code 4yy6), right, with Asn216 H-bonding to each molecule.

We also solved the X-ray structure of the FKBP12-Compound 4-QDPR ternary complex at 1.39-Å resolution (Figure 5a/Table 2). Like BRD9, the structure reveals a 1:1:1 complex of FKBP12-Compound 4-QDPR. Based on the high-resolution data collected, we were able to assign a single diastereomer of compound 4 bound in the complex (Figure S8a). An overlay of the human QDPR-NADH binary complex onto the ternary complex reveals that Compound 4 occupies a portion of the NADH-binding pocket (Figure 5b/S8b). Intriguingly, Compound 4 presents a nucleotide-like ring and triazole that reside in the same space respectively as the adenine and ribose moieties of NADH. Compound 4 makes several interactions with QDPR including a hydrogen bond between Asp41 and a nitrogen of the triazole ring (Figure 5c/S8c.) The 50s and 80s loops of FKBP12 interact with two regions bracketing the QDPR active site (Figure 5d/S8d). Most of the FKBP12-QDPR protein–protein interactions are non-polar except the H-bonding interactions between FKBP12 residues Ala84 and Thr85 with Glu44 of QDPR.

**Figure 5.**
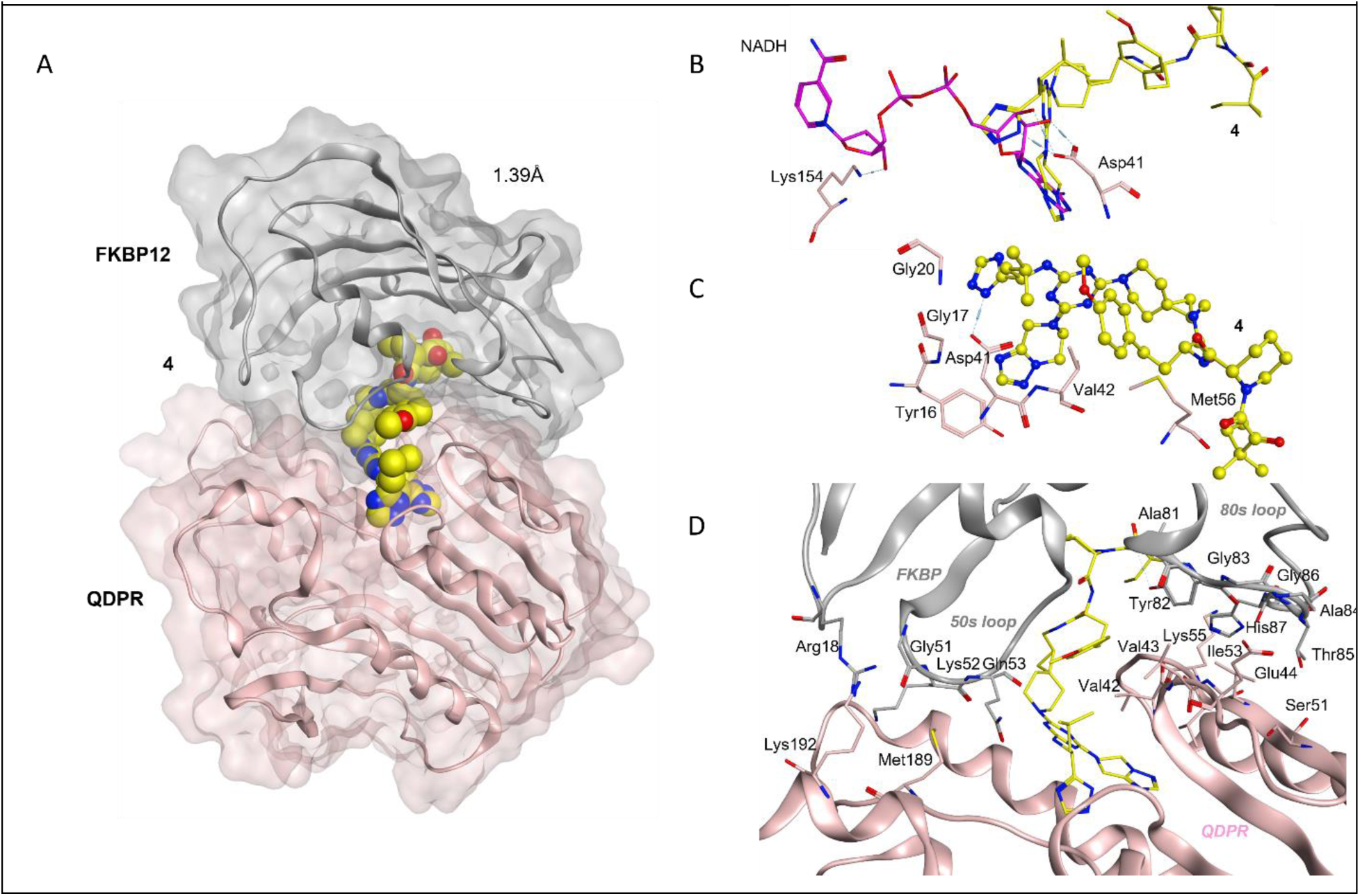
X-ray structure of FKBP12-cpd 4-QDPR ternary complex. A. 1.39-Å resolution structure of FKBP12-Compound 4-BRD9 ternary complex; FKBP12 in grey, 4 in yellow and QDPR in salmon; B. An overlay of Compound 4 (yellow) with the ribose-adenine portion of NADH (salmon, PDB ID code 4hdr); C. Compound 4-QDPR interactions within 3.5 Å, 4 shown as ball-and-stick. Note Asp41 H-bonding to Compound 4; D. FKBP12-QDPR protein-protein interactions, with FKBP12 (highlight of 50s and 80s loops) and QDPR. Note QDPR Glu44 H-bonding to FKBP12 Ala84 and Thr85.

## DISCUSSION

We synthesized an FKBP12 presenter-biased DEL to explore the potential of ternary complex DEL screening aimed at the discovery of FKBP molecular glues. Using protein domains characterized by X-ray crystallography (structures deposited in the Protein Data Bank), our CIP-DEL screens revealed that FKBP12 molecular glues can be identified. Interactions with targets were assessed using quantitative binding measurements of the CIP to the target protein in the absence or presence of FKBP12. We discovered FKBP12 molecular glues for the bromodomain-containing protein 9 (BRD9) and the non-bromodomain containing quinoid dihydropteridine reductase (QDPR). These CIPs reconstitute FKBP12 ternary complex formation in both biophysical and cellular assays. Both Compound 1 and 4 are chemical probes for investigating FKBP12 compound-dependent modulation of BRD9’s and QDPR’s functions in cells.

The high-resolution X-ray structures demonstrate that for BRD9, Compound 1 binds the acyl-lysine binding pocket of BRD9. For QDPR, Compound 4 interacts with a portion of the NADH-binding pocket. In both cases, only a single hydrogen bond is observed between the molecular glue and the target protein and the additional remaining compound interactions with the targets are nonpolar interactions, which is a characteristic of molecular glues and consistent with our inability to demonstrate any significant compound affinity to the targets alone. Lastly, FKBP12 loops (40s, 50s, 80s) known to contact other molecular glue targets^12^ form contacts with both BRD9 and QDPR, while FKBP12 His87 is the only shared protein–protein contact residue between the two ternary complexes, demonstrating the plasticity of FKBP12 and its ability to form unique contacts to drive cooperative interactions between disparate targets. Intriguingly, His87 of FKBP12 also forms protein–protein interactions with mTOR, calcineurin, and CEP250 providing increased evidence for the importance of the 80’s loop in diverse target recognition^12^.

Notably, in both FKBP12 ternary complexes, the molecular glues make contacts within hot spots that mimic interactions with both target’s known endogenous ligands. In the case of BRD9, the Asn216 Nδ2 hydrogen forms an H-bond with a histone’s acyl-lysine carbonyl oxygen and the same Asn216 Nδ2 hydrogen forms an H-bond with Compound 1’s carbonyl oxygen. In addition, the adjacent atoms of Compound 1 overlay with those of butyryl-lysine, a remarkable ability of a DEL screen to identify moieties that mimic endogenous ligand interactions without structure-guided design. The CIPs have co-opted BRD9’s propensity to recognize the PTM, butyryl-Lys, by displaying Compound 1 on the surface of FKBP12, much like IMiD drugs co-opt cereblon by resembling cereblon’s endogenous C-terminal cyclic imide degron^24^.

QDPR is an enzyme that catalyzes the reduction of quinoid dihydrobiopterin into tetrahydrobiopterin using the cofactor nicotinamide adenine dinucoletide (NADH), which consists of two nucleotides joined through phosphate groups with each nucleotide containing an adenine base or nicotinamide. This non-canonical reduction using NADH rather than NADPH distinguishes QDPR from other NAD(P)H-dependent enzymes^25^. QDPR Asp41 makes two hydrogen bonds with the cis-hydroxyls on the adenosine ribose ring of NADH, a highly conserved interaction known to confer selectivity of oxido-reductases for NADH over NADPH, as NADPH bears a 2’ phosphorylated adenosine ribose^26^. Compound 4 presents a structural motif that occupies the region of the QDPR NADH-binding site that overlays only with the adenosine portion of NADH. Like that observed for NADH, QDPR Asp41 makes one hydrogen bond with the triazine ring of Compound 4 that occupies the region bound to the adenosine ribose ring of NADH. Similarly to the adenine ring of NADH, Compound 4 presents a bicyclic ring into the active-site, with key differences, notably a reversal of the 5-6 membered rings and saturation of the six-membered ring. In addition, as we were able to assign a single diastereomer based on our X-ray data, we assume that the biochemical and biophysical binding we observed was due solely to that isomer, and thus the apparent ternary Kd/EC50 of 3.0/3.8 µM is most likely an underestimate of its true potency. The display of an endogenous ligand mimic (a co-factor in the case of NADH-QDPR) on the surface of FKBP12 results in a molecular glue binding through a combination of small molecule-protein and protein–protein interactions. The display from FKBP12 of a compound having an element that resembles the key motif of the target protein’s substrate, while only partially occupying the binding site, leverages protein plasticity to form ternary complexes. This feature is reminiscent of neo-protein-protein interactions resulting from oncogenic missense mutations^27^.

In this study, we identified and validated molecular glues for two protein targets. Key binding elements of our screening hits suggest where to focus library expansion in the future. For both BRD9 and QDPR, the molecular glues are compounds that showed high selectivity for the linker. From the screening hit profiles for both BRD9 and QDPR, in each case, only 1 of the 44 linkers in the FKBP-CIP DEL library yielded molecular glues with apparent high cooperativity. Given that for BRD9 the 4-aminopiperidine linker and for QDPR the spiro [5.2] piperidine based-linker were essential for formation of the ternary complex, libraries elaborated with an even greater array of linkers may be even more effective. Using multiple FKBP-ligands and/or exit vectors in the construction of future FKBP-CIP-DELs may yield more opportunities for FKBP12-target molecular-glue interactions. The FKBP CIP-DEL from a triazine-scaffold resulted in motifs that may be too branched and polar for optimal molecular-glue interactions.

Compound 4 was observed to interact only with the adenosine-binding portion of the QDPR active-site in our X-ray structure. It is possible that Compound 4, or derivatives, can bring about recruitment of FKBP12 to other NADH-binding proteins, or even simply adenosine-binding pockets such as ATP binding sites. If so, rational selection of targets related to previous successes will lead to further increases in hit rates.

We hope to catalyze an expansion in FKBP-molecular glue targets. We believe that continued exploration in CIP-DEL ternary complex screening should improve upon our understanding of what is possible for FKBP molecular glues as chemical probes and as new medicines.

## AUTHOR CONTRIBUTIONS

Trevor A. Zandi performed, interpreted, and summarized all the experiments in the paper, except for the X-ray crystallography experiments. Bingqi Tong synthesized Compound 4. Zher Yin Tan designed and produced the FKBP CIP-DEL library. Jessica S. Viscomi purified QDPR for co-crystallization experiments. Michael J. Romanowski and Jessica S. Viscomi performed and generated the X-ray data for both ternary complexes. Simone Bonazzi managed the outsourcing of compounds 1-3. Frédéric J. Zécri and Stuart L. Schreiber provided project guidance, operational support and assisted with data interpretation. Gregory A. Michaud generated the protein structure figures and both Trevor A. Zandi and Gregory A. Michaud drafted the manuscript.

## METHODS

### FKBP-CIP-DEL library synthesis

The synthesis of FKBP12 CIP-DEL can be found in Tan et. al. (just accepted). Briefly, the FKBP binding motif was adapted from Ligand 1 in Holt et al.^16^, where the ester was substituted to an amide to tolerate on-DNA chemistry, and the cyclohexyl ring was modified to accommodate different connectors that link the FKBP binding ligand to the triazine library. 44 connector-conjugated FKBP12 binding ligands was synthesized and attached to DNA via a CuAAC reaction, followed by sequential SNAr to attach the triazine core and two amine building blocks (290 for cycle 2 and 249 for cycle 3) via split-pool synthesis, with DNA barcodes ligated before each chemical reaction, to yield a 3.2-million-member library.

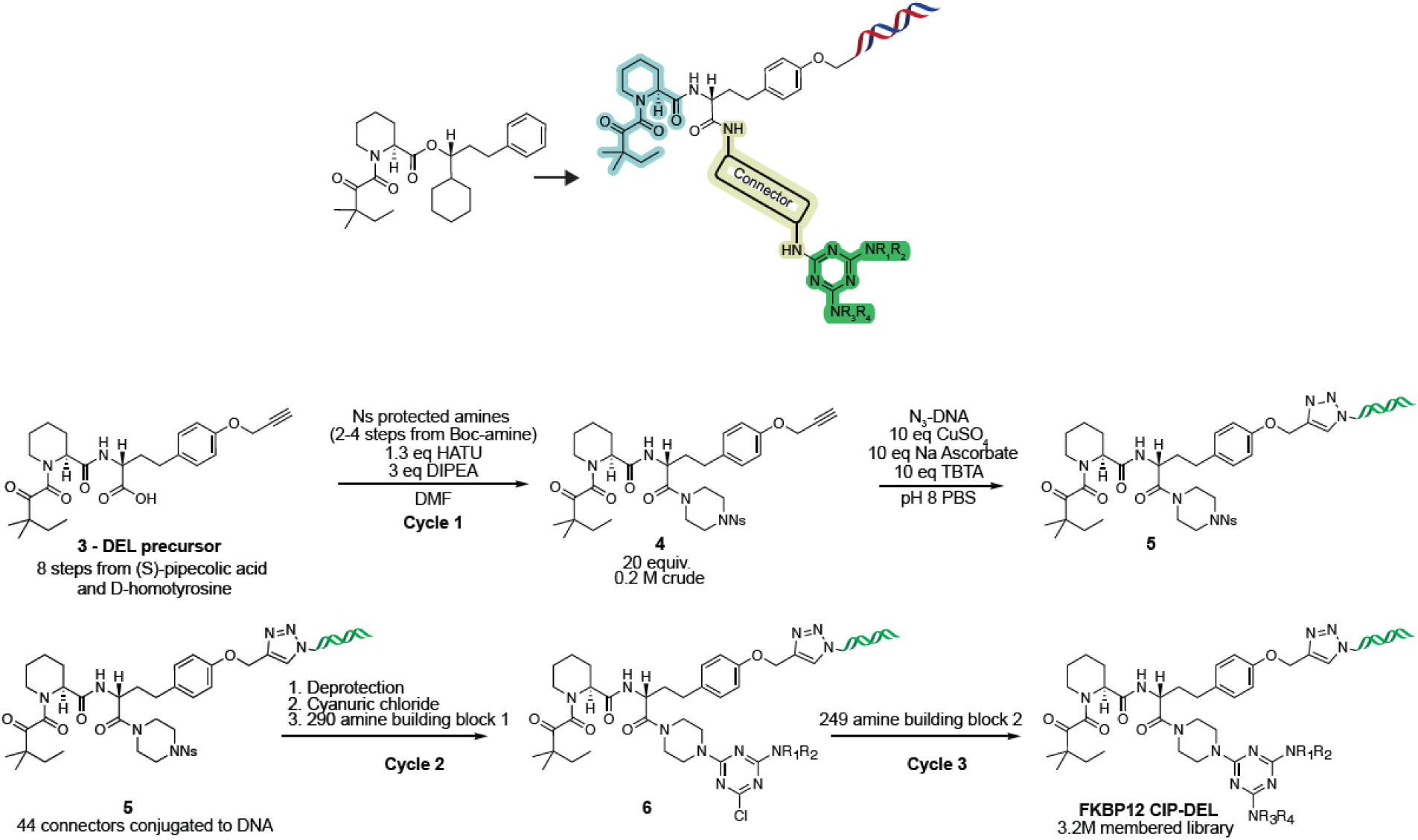

### High throughput protein expression and purification for CIP-DEL screening

A 0.5 µL portion of purified plasmid DNA was added to 10 µL *E. coli* BL21-CodonPlus (DE3)-RIPL cells (Agilent Technologies) in a 96-well PCR-plate (ThermoFisher) and incubated on ice for 10 minutes. The cell-DNA mixture was heat-shocked at 42 °C for 25 seconds. To each well was added 180 µL of SOC media (Corning®), the plate was covered with AirPore tape sheets (Qiagen), and the plate was shaken at 250 RPM at 37 °C for 1 hour. A Biomek FX (Beckman Coulter) was used to transfer 3 µL of each transformant to a Nunc omnitray plate with LB agar containing 50 µg/mL kanamycin. Plates were incubated overnight at 37 °C. Colonies were scraped with a Biomek FX and were inoculated individually into 1 mL of TB media (Corning) containing 50 µg/mL kanamycin sulfate in a 2 mL block plate. Cultures were grown with shaking at 250 rpm overnight. Overnight cultures were diluted by addition of 100 µL into 2 mL block plates each holding 0.9 mL of TB media containing 50 µg/mL kanamycin sulfate per well, and the process was repeated eight times to generate eight identical plates. Plates were grown until an OD600 of 0.4-0.6 before induction by addition of either IPTG or arabinose to a concentration of 1 mM or 0.08%, respectively. Induced cultures were shaken overnight at 18 °C. The next morning, cultures were transferred to 1 mL Masterblock plates (Greiner), sealed with a storage mat (Corning) and centrifuged at 4000 x g for 15 minutes to pellet cells. Media was removed from cell pellets by a Biomek FX and pellets were frozen at −80 °C for later purification.

Cells were thawed and lysed by resuspension in 200 µL of B-PER Complete Bacterial Protein Extraction Reagent (ThermoFisher) and inverted for 1 hour with a Rotator Genie (Scientific Industries). Using a Multidrop Combi+ (ThermoFisher), 400 uL of Wash Buffer I (50 mM Tris pH 7.5 (Teknova), 400 mM NaCl (Teknova), 15 mM imidazole (Teknova), 10% glycerol (Teknova)) was added to each well. Plates were then centrifuged at 4000 x g for 30 minutes to sediment the cellular debris. Supernatant was transferred to fresh 1 mL Masterblock plates equipped with a 100 µL 33% slurry of HisPure Ni-NTA resin (ThermoFisher) which had been equilibrated in Wash Buffer I using a Biomek FX. The supernatant and resin were incubated with rotation at 4 °C on a Rotator Genie for 1 hour. The resin was then sedimented by centrifugation at 2000 x g for 1 min and 400 µL supernatant was discarded with a Biomek FX. Resin was resuspended in remaining supernatant and added to 500 µL 0.45 µm AcroPrep filter plates (Cytiva). Filter plates were placed above 250 µL 96-well collection plates (Greiner) and centrifuged at 2000 x g for 1 minute at 4 °C. Supernatant from the collection plate was discarded and an additional 200 µL of Wash Buffer was added to the filter plate followed by centrifugation at 2000 x g for 1 minute at 4 °C to collect and discard the wash buffer. This wash step was repeated twice, followed by three additional washes with Wash Buffer II (50 mM Tris pH 7.5, 400 mM NaCl, 30 mM imidazole, 10% glycerol). After the final wash was discarded, the filter plate was centrifuged for 1 minute at 4 °C to remove residual wash buffer. A fresh 250 µL collection was placed under the filter plate and 35 µL Elution Buffer (50 mM Tris pH 7.5, 150 mM NaCl, 250 mM imidazole, 10% glycerol) was added, followed by incubation for 30 minutes at 4 °C, and elution by centrifugation at 2000 x g for 1 minute at 4 °C. This process was repeated and the eight identical elution plates were combined in a 1 mL Masterblock plate. Samples were concentrated to ~0.1 mL in 0.5 mL Amicon Ultra Centrigual Filters with a 3 kDa MW cut off (EMD Millipore) and imidazole was removed by two passes through ZEBA Spin Desalting Plates which had been equilibrated in protein storage buffer (50 mM HEPES pH 7.5 (Teknova), 150 mM NaCl, 10% glycerol), following the manufacturer’s instructions. The protein samples were analyzed for yield and purity by SDS-PAGE Criterion gels in a Dodeca cell (Bio-Rad). Proteins with visible expression by SDS-PAGE were analyzed for homogeneity by analytical size exclusion chromatography on a BioInert HPLC (Agilent) with a SuperDex S75 10/300 GL column (Cytiva). Targets with at least ~90% homogeneity were flash frozen in liquid nitrogen, stored at −80 °C, and prioritized for FKBP-CIP-DEL screens.

### Large-scale protein expression and purification (for biophysics)

A 1.0 uL portion of plasma DNA was added to 20 uL *E. coli* BL21-CodonPlus (DE3)-RIPL cells (Agilent Technologies) in sterile 1.5 mL microcentrifuge tubes (VWR) and incubated on ice for 10 minutes. The cell-DNA mixture was then heated at 42 °C for 25 seconds. The mixture was recovered in 1 mL of SOC media at 37 °C for 1 hour with shaking at 250 rpm. A 0.1 mL sample was then removed from the recovered transformation and plated on 100 mm LB agar plates with 50 µg/mL kanamycin (Teknova). The plate was incubated at 37 °C overnight and a single colony was used to inoculate 75 mL of TB media with 50 µg/mL kanamycin sulfate in a 250 mL polycarbonate, baffled Erlenmeyer flask with a vented cap. Inoculated media was grown overnight at 37 °C with shaking at 250 rpm. The next morning, three 4.0 L flasks were filled with 1.0 L each of TB media with 50 µg/mL kanamycin sulfate and inoculated with 20.0 mL each of the overnight starter culture. Cultures were grown at 37 °C with shaking at 250 rpm until they reached OD600 of 0.4-0.6 at which cultures were induced with either a final concentration of 1 mM IPTG or 0.08% arabinose. Protein was then expressed from induced cultures overnight at 18 °C with shaking at 250 rpm. Cultures were pelleted by centrifugation at 4000g for 10 min in 1.0 L centrifuge flasks (Nalgene), media was discarded, and pellets were frozen in liquid nitrogen and stored at −80 °C. Cell pellets were thawed and lysed by addition of 4.0 mL B-PER Complete Bacterial Protein Extraction Reagent per gram of wet cell mass in a 50.0 mL centrifuge tube and inversion by a Rotator Genie for 30 minutes at 4 °C. To the lysate was added 8.0 mL of Wash Buffer I (50 mM Tris pH 7.5, 150 mM NaCl, 15 mM imidazole, 10% glycerol) and lysate was then clarified by centrifugation. The supernatant was transferred to a clean 50.0 mL centrifuge tube, charged with 2.0 mL of Wash Buffer I equilibrated HisPure Ni-NTA resin per gram of wet cell mass. The supernatant-resin mixture was rotated with inversion by a Rotator Genie for 1.5 hours at 4 °C, at which point the supernatant was strained through the glass frit of an empty Econo-Pac chromatography column (Bio-Rad). The resin was washed with three portions of 12.0 mL of Wash Buffer I, followed by three portions of Wash Buffer II (50 mM Tris pH 7.5, 150 mM NaCl, 30 mM imidazole, 10% glycerol). The protein of interest was eluted with two portions of 4.0 mL of Elution Buffer (50 mM Tris pH 7.5, 150 mM NaCl, 250 mM imidazole, 10% glycerol). The sample was concentrated to 1 mL in a 15 mL Amicon Ultra Centrifugal Filter (EMD Millipore) and imidazole was removed over a PD MidiTrap G-25 column (Cytiva) which had been equilibrated in protein storage buffer (50 mM HEPES pH 7.5, 150 mM NaCl, 10% glycerol). Purity and homogeneity was verified by SDS-PAGE and analytical size exclusion chromatography, respectively. Samples were frozen in liquid nitrogen and stored at −80 °C.

### DNA-encoded library selection

To a 1.5 mL microfuge tube, an amount of 10 µL per sample of magnet His-Tag Dynabeads (Invitrogen) were added, and magnetic beads were sedimented with a MagJET separation rack (ThermoFisher). Storage solution was removed and the resin was washed with 3×1.0 mL of target-binding buffer (25 mM HEPES pH 7.4, 150 mM NaCl, 0.05% tween-20 (VWR)). Beads were then diluted to 100 uL per sample in target-binding buffer. Two 96-well KingFisher plates (ThermoFisher) were prepared for automated DEL selection. In plate I, 100 µL of prepared magnetic beads was added to row A, 100 uL of 10 µM His-tagged target protein or target-binding buffer alone as a control was added to row B, 200 µL DEL target-binding buffer was added to rows C and H, DEL-selection buffer (25 mM HEPES pH 7.4, 150 mM NaCl, 0.05% tween-20, 0.3 mg/mL salmon sperm DNA (Invitrogen), 10 mM imidazole, 0.1 mM TCEP) was added to rows D, E, and G, and finally 100 µL of FKBP-CIP-DEL library containing either 10^6^, 5×10^6^, or 10^7^ average copies per member with either 0 or 100 µM FKBP presenter protein was added to row F. Plate II was prepared with 200 µL target-binding buffer in row A as a final wash and in row B was placed 100 µL elution buffer (25 mM HEPES pH 7.4, 150 mM NaCl, 0.05% tween-20). An automated DEL-selection protocol was run at room temperature on a KingFisher Duo system (ThermoFisher) in which target was mixed with His-Tag Dynabeads for 30 minutes, followed by 3×3 minute wash steps, a 1 hour DEL binding step, and 3×3 minute wash steps, before magnetic beads were released for elution. The post-selection target-bound magnetic beads were transferred to PCR-tubes and heated at 95 °C for 10 minutes to denature targets and release bound DNA-encoded molecules. Beads were sedimented with a DynaMag-PCR magnet (ThermoFisher) and solutions of eluted DNA-encoded compounds were transferred to a 96-well PCR plate (VWR).

### Preparation of DNA-encoded library selection for sequencing

The small molecule and headpiece were removed by restriction digest. A 20.0 µL sample of post-selection DNA-encoded compounds was added to 56.5 µL of CutSmart buffer containing 1.0 unit of StuI (New England Biolabs) in a 96-well PCR plate and the samples were incubated at 37 °C. DNA was then purified with a ChargeSwitch PCR clean-Up Kit (ThermoFisher) according to the manufacturers protocol on a KingFisher Duo system. Each sample was quantified by a SybrGreen qPCR protocol on a QuantStudio 7 Flex real-time PCR system (Applied Biosystems) using a standard curve derived from a titration series of the same, pre-selection DNA-encoded library (10^3^ to 10^−2^ copies per member per µL). Samples were amplified, sequencing adapters were added, DNA was indexed by PCR for next-generation sequencing with a variable number of cycles guided by qPCR quantification to target ~5-20 ng per µL of DNA. To 19 uL of DNA was added 25 µL of 2x Platinum HotStart Taq MasterMix (ThermoFisher) and 3 µL each of 10 µM solutions of custom Nextera indexing primers with forward primers of sequence 5’-AATGATACGGCGACCACCGAGATCTACAC[i5 sequence]ACCATGCTCACTGCGGCTTCTACCTCCATCAG-3’, and reverse primers of sequence 5’CAAGCAGAAGACGGCATACGAGAT[i7 sequence]ACCATGCTCACTGCGGCTTCTACCTCCATCAG-3’ to 0.6 µM final concentration. Samples were amplified by the following PCR protocol: denaturation for 2 minutes at 95 °C, N cycles of [15 seconds at 95 °C, 15 seconds at 55 °C, 30 seconds at 72 °C], with a final extension for 7 minutes at 72 °C. DNA post-PCR was quantified by QuBit dsDNA broad range assay kit (ThermoFisher), and DNA was subjected to additional PCR cycles if yield was below 5 ng per µL. DNA was pooled for sequencing by combining 30.0 ng per µL of each sample. Pooled DNA was purified for sequencing using a 2% E-Gel EX agarose gel (Invitrogen) followed by a QIAquick gel extraction kit (Qiagen). DNA was sequenced on a NovaSeq6000 system using the following custom primers (read-1 primer: 5’–CTTAGCTCCCAGCGACCTGCTTCAATGTCGGATAGTG–3’, index-1 primer: 5’-CTGATGGAGGTAGAAGCCGCAGTGAGCATGGT-3’, index-2 primer: 5’–CACTATCCGACATTGAAGCAGGTCGCTGGGAGCTAAG–3’) with 50 cycles of single-end reads and 8 cycles for index 1 and index 2.

### DNA-encoded library data analysis

Next-generation sequencing data was converted into counts of individual bar-coded molecules by adapting and parallelizing the Python scripts described previously (10.1038/s41589-023-01458-4) for compatibility with a high-performance compute cluster. Briefly, barcodes with near-identical unique-molecular identifiers were consolidated, duplicate count data was summed, barcodes with greater than 50 counts against the beads only control were removed, and count data was analyzed directly. Counts were plotted and analyzed in Spotfire Analyst (TIBCO).

### Cellular NanoBiT recruitment assay

Transfection solution was made by diluting Fugene HD 1:18 into minimal essential medium, (OPTIMEM, ThermoFisher). DNA encoding both Herpes simplex virus thymidine kinase (HSVTK) promoter-driven LgBiT tagged FKBP12 and either cytomegalovirus (CMV) promoter-driven SmBiT tagged BRD9, or HSVTK promoter-driven SmBiT QDPR were added 1:1 to the transfection solution to a total DNA concentration of 18.5 µg/mL and the solution was allowed to incubate for 15 minutes at room temperature. Transfection was performed by diluting the DNA-transfection solution mixture 1:10 into HEK293T cells and plating at a density of 0.625 x 10^6^ cells per mL of complete media (DMEM with 10% FBS, ThermoFisher). Cells were grown in an appropriately sized flask (Corning) at 37 °C and 5% CO2 for 24 hours, upon which time cells were trypsinized, diluted in complete media (DMEM with 10% FBS) to 0.5 x 10^6^ cells per mL, and 5 µL per-well was plated in 1536-well plates (Greiner) with a single-tip dispenser (GNF Systems One Tip Dispenser) to give approximately 2500 cells to each well. The plates were covered and incubated at 37 °C with 5% CO2 for 16 hours (overnight). The following morning, the plates were removed from the incubator and brought to room temperature for 10 minutes. An Echo 655 acoustic liquid handler (Labcyte) was used to add 10 nL of compound per well from Echo-compatible 384-well microplates (Labcyte) containing a 1:3 titration series of each compound in 100% DMSO vehicle, as well as 100% DMSO alone as a control. Plates containing transfected cells and compound were returned to the incubator for 2 hours at 37 °C and 5% CO2. Detection reagent was prepared by 1:25 dilution of the furimazine NanoGlo Live Cell Substrate (LCS) (Promega) into a 1:1 mixture of phosphate-buffered saline (ThermoFisher) and NanoGlo LCS buffer (Promega). The plates were brought from the incubator and allowed to cool at room temperature for 10 minutes before 5 µL of detection reagent was added with the single-tip dispenser and incubation for an additional 10 minutes at room temperature. Luminesence measurements of the plate were then read on a PHERAstar FS microplate reader (BMG Labtech). Luminesence was normalized with the following equation: 100 x (sample luminescence-neutral control luminescence)/(neutral control luminesence). The neutral control luminescence is derived from an average of DMSO-only cells from a given transfection.

### Surface Plasmon Resonance

#### Target-compound (binary complexes)

Biotinylated proteins were immobilized to ~1000 RUs for binary kinetics/affinity. Compounds 1-4 were diluted to 50x in DMSO and serially diluted 1:2 six times and 3.1 µL of each solution was added to 150 µL of SPR buffer in a 384-well plate (Greiner). NADH was dissolved in SPR buffer with 2% DMSO at 50X and diluted 1:2 six times in 150 uL of SPR buffer with 2% DMSO. Single-cycle kinetics. Compounds were injected at 45 µL/min with 120 seconds of contact time and 1,200 seconds of disassociation time. Data were fit with a 1:1 kinetic binding model and/or a steady-state binding model, where appropriate, to calculate the kinetic parameters. SPR buffer consisted of 50 mM Hepes pH 7.5, 150 mM NaCl, 0.01% tween-20, 1 mM TCEP and 2% DMSO.

#### Target-cpd-FKBP12 (ternary complexes)

Biotinylated-QDPR or biotinylated-BRD9 was immobilized to approximately ~150 RUs for ternary kinetics/affinity on a streptavidin chip using a Biacore 8K (Cytiva). Compounds 1-4 were diluted to 50x in DMSO, serially diluted 1:2 six times, and 3.1 µL of each solution was added to 150 µL of SPR buffer containing 10 µM FKBP12, for QDPR, 0 or 50 µM NADH, and for BRD9, 10 µM BI-7273, in a 384-well plate. Single cycle kinetics was run using FKBP12 compound mixtures following the method described in binary SPR.

### BRD9 expression and purification for x-ray crystallography

BRD9 bromo domain-encoding DNA (residues 130-240; UniProt accession number Q9H8M2) was codon-optimized for expression in *E. coli* and cloned into the 5’*Bam*HI and 3’*Xho*I sites of pGEX6P-1 (GenScript).

The plasmid was transformed into BL21(DE3) Star cells (Invitrogen) containing the pRare plasmid from Rosetta cells (Novagen). Cells were grown at 37°C to an OD600 of 0.8-1.0 in Terrific Broth Complete (Teknova), the temperature in the incubator-shaker was then lowered to 18°C, and cultures were cooled for 1 hr. Protein expression was induced with 0.3 mM IPTG (Teknova) and expression continued overnight at 18°C.Cells were harvested by centrifugation and the pellets resuspended in a lysis buffer containing 1X PBS pH 7.4 buffer, 400 mM KCl, 1 mM MgCl2, 5 mM DTT (Sigma), 10% glycerol plus 100 µg/ml DNAse I (Sigma), EDTA-free protease inhibitor tablets (Pierce) and 1 mM PMSF (Calbiochem). All buffers, salts and glycerol were supplied by Teknova. Cells were homogenized for 1 min and passed 2X through a microfluidizer. The lysates were centrifuged for 30 minutes at 199,000 x *g* in a Ti-45 rotor at 4°C to pellet cell debris. The supernatant was applied to Glutathione Sepharose 4b FF beads (Cytiva) and incubated with gentle stirring for 1-2 hrs. The slurry was decanted into a glass Econo-column (Bio-Rad) to remove the lysate. The beads were washed with 200 CV with a wash buffer containing 1X PBS pH 7.4, 400 mM KCl, 5 mM DTT and 10% glycerol. The protein was eluted with 200 mM Tris pH 8.0, 400 mM KCl, 5 mM DTT and 15 mM reduced glutathione (MP Biochemicals). Following dialysis against 2 4-L changes of the buffer consisting of 1X PBS pH 7.4, 400 mM KCl and 5 mM DTT, and overnight cleavage with non-cleavable GST-tagged HPV 3C protease at 4°C at a 1:2000 molar ratio of recombinant fusion protein to protease, the GST tag was removed on fresh glutathione beads. The GST-free protein was concentrated and applied to a Superdex-200 size-exclusion column pre-equilibrated with 25 mM HEPES pH 7.5, 100 mM KCl and 5 mM DTT. The monodisperse fractions were pooled, concentrated to 5.29 mg/ml and flash-frozen in aliquots suitable for subsequent crystallization experiments. The final yields were 1.06 mg of purified protein per 1 g of *E. coli* biomass.

### QDPR expression and purification for x-ray crystallography

Full-length QDPR (UniProt accession number P09417) was codon-optimized for expression in *E. coli* as an N-terminal TEV protease-cleavable 6xHis tag fusion and cloned into pSpeedET plasmid. Protein expression was performed as described for BRD9 except that 1 mM IPTG was used to induce expression. Cells were harvested by centrifugation and the pellets resuspended in a lysis buffer containing 50 mM Tris-HCl pH 8, 500 mM NaCl, 20 mM imidazole, 5% glycerol, 2 mM DTT, 1 mM MgCl2, 100 µg/ml DNAse I (Sigma) and EDTA-free protease inhibitor tablets (Pierce). All buffers, salts and glycerol were supplied by Teknova. Cells were homogenized for 1 min and passed 2X through a microfluidizer. The lysates were centrifuged for 30 minutes at 199,000 x *g* in a Ti-45 rotor at 4°C to pellet cell debris. The supernatant was applied to Ni-NTA beads and stirred gently for 1.5 hr at 4°C to maximize protein binding. The slurry was decanted into a glass Econo-column (Bio-Rad) to remove the lysate. The beads were washed with 200 CV with a wash buffer containing 50 mM Tris-HCl pH 8, 500 mM NaCl, 20 mM imidazole, 2 mM DTT and 5% glycerol. The protein was eluted with 50 mM Tris-HCl pH 8, 500 mM NaCl, 400 mM imidazole, 2 mM DTT and 5% glycerol. Following dialysis against 2 4-L changes of the buffer consisting of wash buffer without glycerol, and overnight cleavage with TEV protease at 4°C at a 1:1000 molar ratio of recombinant fusion protein to protease, the His tag was removed on fresh Ni-NTA beads pre-equilibrated with the cleavage buffer. The His tag-free protein was concentrated and applied to a Superdex-75 16/60 size-exclusion column pre-equilibrated with 50 mM HEPES pH 7.4, 150 mM NaCl and 2 mM DTT. The monodisperse fractions were pooled, concentrated to 10 mg/ml and flash-frozen in aliquots suitable for subsequent crystallization experiments. The final yields were 1.4 mg of purified protein per 1 g of *E. coli* biomass.

### Protein crystallization

The Compound 1-mediated FKBP12-BRD9 bromo domain complex, in a final protein buffer composed of 20 mM HEPES pH 7.5, 150 mM NaCl and 1 mM TCEP, was concentrated to 10 mg/ml. Co-crystals were obtained by sitting-drop vapor diffusion in 96-well plates at 4°C against a reservoir of 0.1 M HEPES pH 7.5 and 20% (w/v) PEG 8000. The Compound 4-mediated FKBP12-QDPR complex, in a final protein buffer composed of 50 mM HEPES pH 7.4, 150 mM NaCl and 2 mM DTT, was concentrated to 10 mg/mL. Co-crystals were obtained by sitting-drop vapor diffusion in 96-well plates at 4°C against a reservoir of 0.1 M Tris-HCl pH 7,0.2 M (CH3COO)2Ca·H2O and 20% (w/v) PEG 3000. 0.15 nL of the protein sample and 0.15 nL of the crystallization buffer were used per drop in both co-crystallization experiments. The crystals appeared within a week of incubation. All crystals for data collection were cryoprotected in mother liquor supplemented with 20% (v/v) glycerol for 30-90 sec and immersion in liquid nitrogen.

### Data collection, processing, and structure determination

The synchrotron data was collected under standard cryogenic conditions on Beamline P14 operated by EMBL Hamburg at the PETRA III storage ring (DESY, Hamburg, Germany) using a Dectris EIGER2 CdTe 16M photon-counting detector, integrated with *XDS* and scaled with *XSCALE*^28^. The structures were determined from single-wavelength native diffraction experiments by molecular replacement with *PHASER*^29^ using search models from previously determined structures (PDB ID 8ahc, 1hdr and 7u8d for BRD9, QDPR and FKBP12 respectively)^21,30,31^ The refinement of the initial solutions with *BUSTER*^32^ yielded experimental electron density maps suitable for model building with *Coot* ^33^Final refinement of the atomic coordinates was performed in *PHENIX*^34^

The BRD9 residues 130-137 (chain E), as well as residues 130-137 and 239-240 (chains F, G and H) in 9du1, and QDPR residues 1-9 (chain A) in 9dtw, were not visible in the electron density maps and were omitted from refinement of the final atomic models. *PROCHECK*^35^ revealed no disallowed (φ, ψ) combinations and excellent stereochemistry (see Table 1 for a summary of X-ray data and refinement statistics). All structure figures were prepared with MOE (Molecular Operating Environment). The atomic coordinates and structure factors have been deposited in the Protein Data Bank (PDB ID codes 9du1 and 9dtw).

### Compounds (1-4): Synthesis and Characterization

Starting materials, reagents, and solvents were obtained from commercial sources with purity >95% unless otherwise noted and used as received. THF, DCM, and DMF were anhydrous grade. Analytical HPLC was conducted using an Agilent 1260 series with UV detection at 210 nm. Analytical LC/MS was conducted using either an Agilent 1200 series with UV detection at 210 nm and MS with electrospray mode (ESI) coupled with mass detector or Agilent 1100 series with UV detection at 214 and 254 nm and MS with electrospray mode (ESI) coupled with a Waters ZQ single quad mass detector or a Waters Classic AcQuity UPLC with UV detection at 214 and 254 nm and an electrospray mode (ESI) coupled with a Waters SQ single quad mass detector. Purification of intermediates and final products was carried out on normal phase using an ISCO CombiFlash system and prepacked SiO2 cartridges eluted with optimized gradients of either heptane-ethyl acetate mixture or dichloromethane-methanol as described. Preparative high pressure liquid chromatography (HPLC) was performed on ISCO. HPLC separation was performed at 75 mL/min flow rate with the indicated gradient using formic acid modifier as described. Columns were RediSep® Prep C18Aq Column 5 μm (30 mm x 100 mm). All target compounds had purity of >95% as established by analytical HPLC, unless otherwise noted. NMR spectra were recorded on a Bruker Ultrashield AV400 (Avance 400 MHz) plus or a AV300 (Avance 300 MHz) instrument. Chemical shifts (d) are reported in parts per million (ppm) relative to deuterated solvent as the internal standard (CDCl3 7.26 ppm, DMSO-*d*6 2.50 ppm, CD3OD 3.31 ppm), and coupling constants (*J*) are in hertz (Hz).

### Synthetic scheme for compound 8

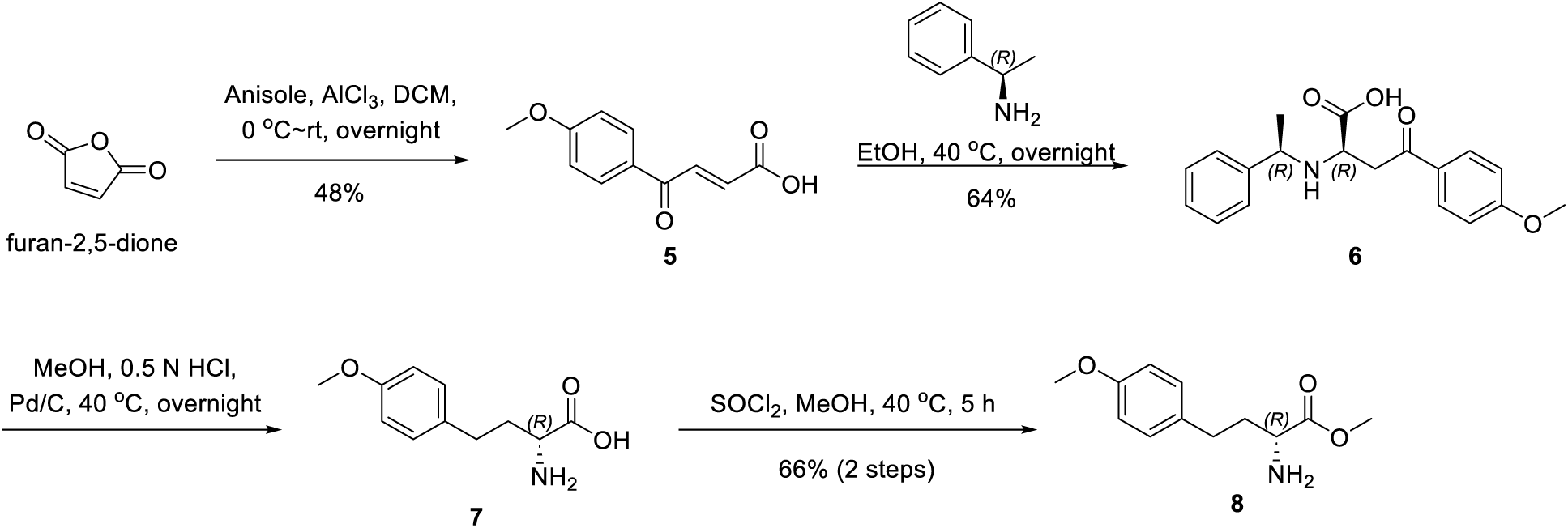

### (E)-4-(4-methoxyphenyl)-4-oxobut-2-enoic acid (5)

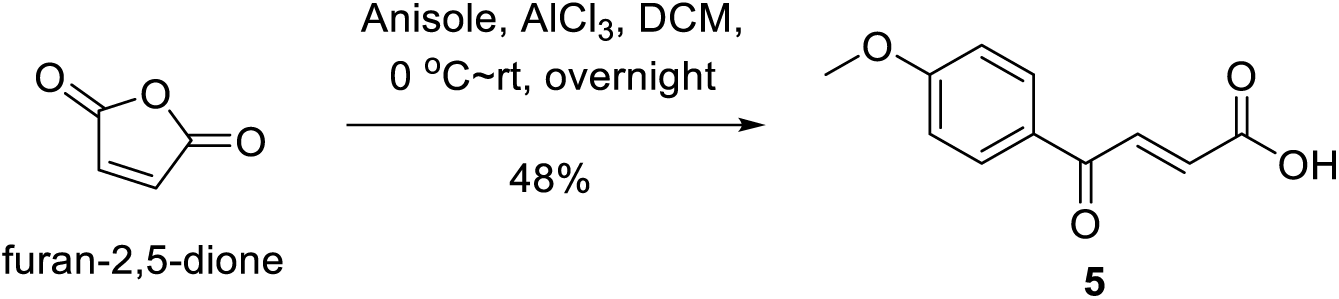

To a solution of furan-2,5-dione (200.0 g, 2.0 mol, 1.0 eq) in DCM (4 L) was added AlCl3 (400.0 g, 3.0 mol, 1.5 eq) in portions at 0 °C. Anisole (220.0 g, 2034.4 mmol, 1.0 eq) in DCM (1 L) was added to the mixture at 0 °C and then the reaction mixture was stirred at room temperature overnight allowing to return to rt. The mixture was poured into iced water (1000 mL), extracted with EtOAc (2000 mL x 3). The combined organic phase was washed with brine (2000 mL), dried over Na2SO4 and concentrated to give a residue. The residue was triturated with DCM (800 mL) to afford (E)-4-(4-methoxyphenyl)-4-oxobut-2-enoic acid (**5**) (200.0 g, 48% yield) as a yellow solid.

**^1^H NMR (300 MHz, CDCl3):** *δ* ppm 11.33 (brs, 1H), 8.03-7.98 (m, 3H), 6.99 (d, *J* = 8.8 Hz, 2H), 6.88 (d, *J*

= 15.5 Hz, 1H), 3.89 (s, 3H).

**LCMS:** 206.9 ([M+H]^+^).

### (R)-4-(4-methoxyphenyl)-4-oxo-2-(((R)-1-phenylethyl)amino)butanoic acid (6)

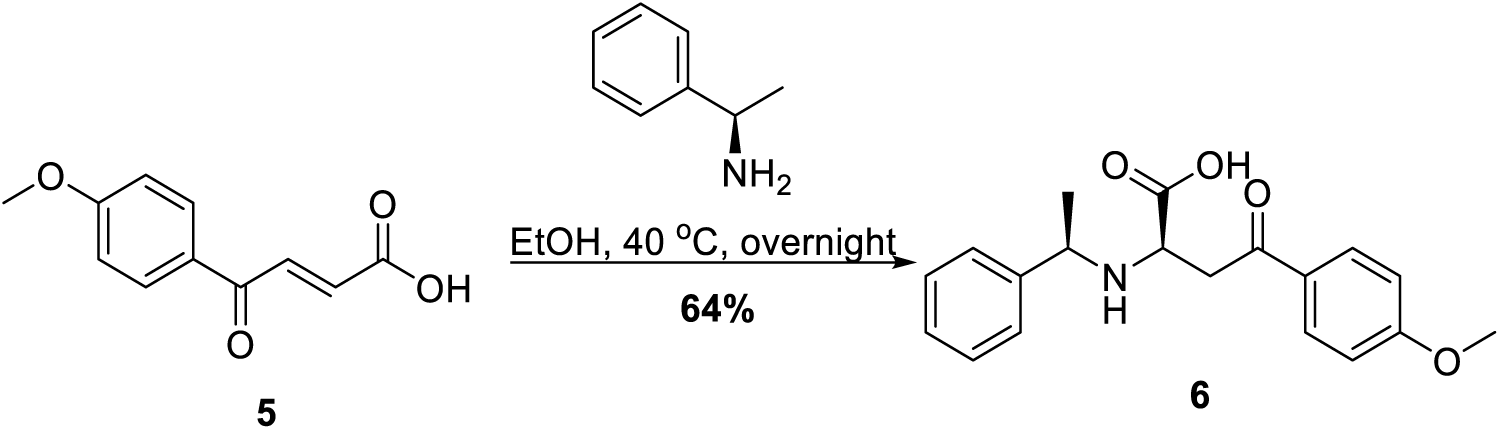

To a solution of (E)-4-(4-methoxyphenyl)-4-oxobut-2-enoic acid (**5**) (200.0 g, 969.9 mmol, 1.0 eq) in EtOH (4 L) was added (*R*)-1-phenylethan-1-amine (176.0 g, 1454.9 mmol, 1.5 eq) at 40 °C. The reaction was stirred at 40 °C overnight. The mixture was filtered, the filter cake was collected and triturated in MeOH (500 mL) to afford (R)-4-(4-methoxyphenyl)-4-oxo-2-(((R)-1-phenylethyl)amino)butanoic acid (**6**) (204.0 g, 64% yield) as a white solid.

**^1^H NMR (400 MHz, DMSO-*d6*):** *δ* ppm 7.88 (d, *J* = 8.8 Hz, 2H), 7.29-7.22 (m, 5H), 7.02 (d, *J* = 8.8 Hz, 2H),

3.99 (q, *J* = 4.5 Hz, 1H), 3.84 (s, 3H), 3.32 (t, *J* = 6.2 Hz, 1H), 3.21-3.22 (m, 2H), 1.26 (d, *J* = 2.6 Hz, 3H).

**LCMS:** 328.2 ([M+H]^+^).

### (R)-2-amino-4-(4-methoxyphenyl)butanoic acid (7)

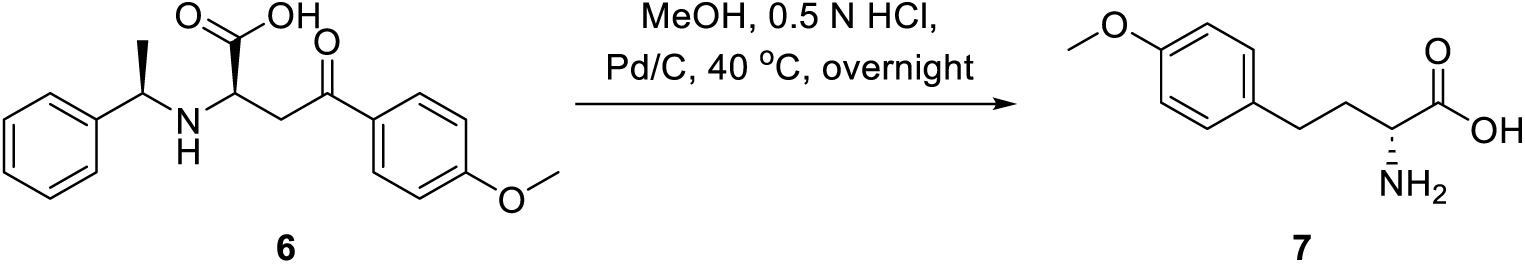

To a solution of (R)-4-(4-methoxyphenyl)-4-oxo-2-(((R)-1-phenylethyl)amino)butanoic acid (**6**) (100.0 g, 305.4 mmol, 1.0 eq) in MeOH (3 L) and 0.5 N HCl (3 L) was added Pd/C (20.0 g, 20% wt). The reaction mixture was stirred at 40 °C overnight under H2 atmosphere. The mixture was filtered, the filtrate was concentrated to afford (R)-2-amino-4-(4-methoxyphenyl)butanoic acid (**7**) (200.0 g, crude) as a white solid.

**^1^H NMR (400 MHz, D2O):** *δ* ppm 7.29 (d, *J* = 8.4 Hz, 2H), 7.00 (d, *J* = 8.3 Hz, 2H), 3.84 (s, 3H), 3.78 (t, *J* =

5.9 Hz, 1H), 2.73-2.67 (m, 2H), 2.19-2.10 (m, 2H).

**LCMS:** 210.2 ([M+H]^+^).

### methyl (R)-2-amino-4-(4-methoxyphenyl)butanoate (8)

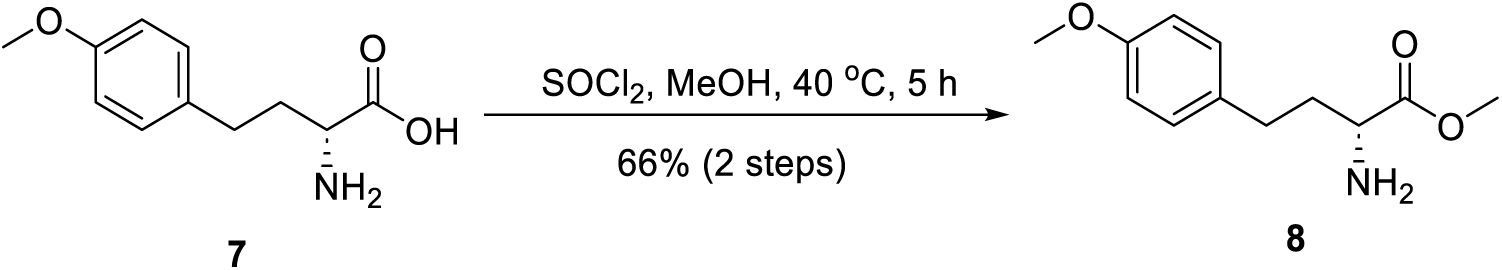

To a solution of (R)-2-amino-4-(4-methoxyphenyl)butanoic acid (**7**) (100.0 g, 478.0 mmol, 1.0 eq) in MeOH (1 L) was added SOCl2 (170.6 g, 1434.0 mmol, 3.0 eq) and the reaction mixture was stirred at 40 °C for 5

1. h. The solvent was removed in vacuo to afford methyl (R)-2-amino-4-(4-methoxyphenyl)butanoate (**8**) (45.0 g, 66% over 2 steps) as a white solid, which was directly used to the next step without further purification.

**^1^H NMR (400 MHz, CD3OD):** *δ* ppm 7.14 (d, *J* = 8.6 Hz, 2H), 6.87(d, *J* = 8.6 Hz, 2H), 4.02 (t, *J* = 6.4 Hz, 1H), 3.82 (s, 3H), 3.75 (s, 3H), 2.73-2.64 (m, 2H), 2.24-2.05 (m, 2H).

**LCMS:** 224.2 ([M+H]^+^).

### Synthetic scheme for compound 15

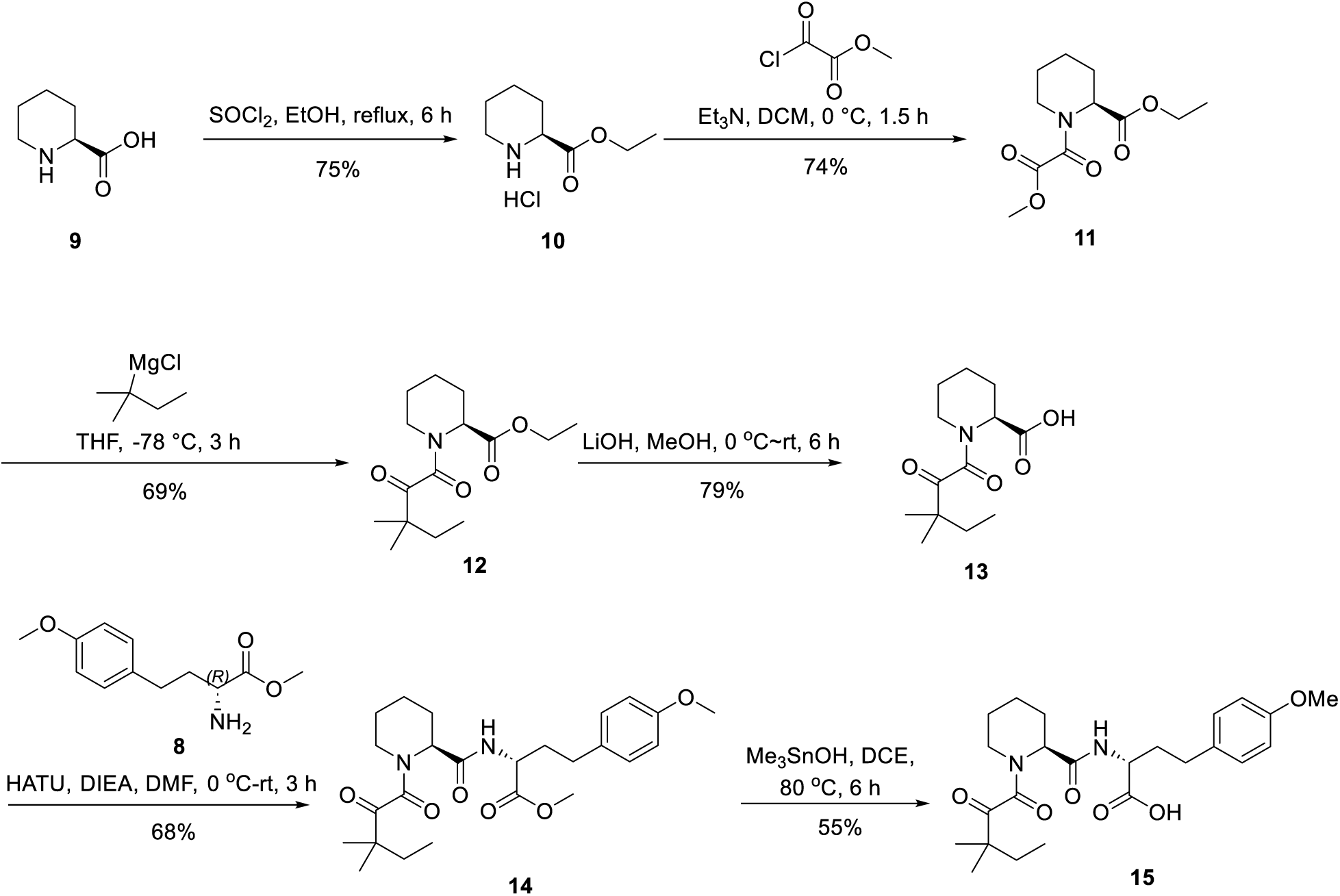

### Ethyl (S)-piperidine-2-carboxylate hydrochloride (10)

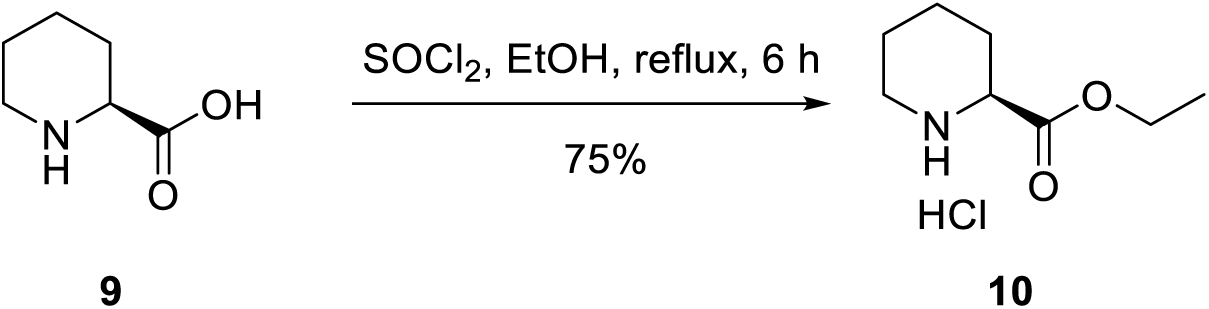

To a solution of (S)-piperidine-2-carboxylic acid (**9**) (385.0 g, 3.0 mol, 1.0 eq) in EtOH (5.0 L) was added SOCl2 (1.1 Kg, 9.0 mol, 3.0 eq) dropwise at room temperature. The mixture was stirred at reflux for 6 hours. The mixture was concentrated in vacuo to afford ethyl (S)-piperidine-2-carboxylate hydrochloride (**10**) (350.0 g, 75% yield) as a white solid, which was directly used to the next step without further purification.

**^1^H NMR (300 MHz, CDCl3):** *δ* ppm 9.89 (s, 1H), 9.67 (s, 1H), 4.26 (q, *J* = 7.1 Hz, 2H), 3.93 (s, 1H), 3.60-3.56 (m, 1H), 3.11-3.08 (m, 1H), 2.25-2.21 (m, 1H), 2.12-1.98 (m, 2H), 1.86-1.75 (m, 2H), 1.62-1.55 (m, 1H), 1.27 (t, *J* = 7.1 Hz, 3H).

### Ethyl (S)-1-(2-methoxy-2-oxoacetyl)piperidine-2-carboxylate (11)

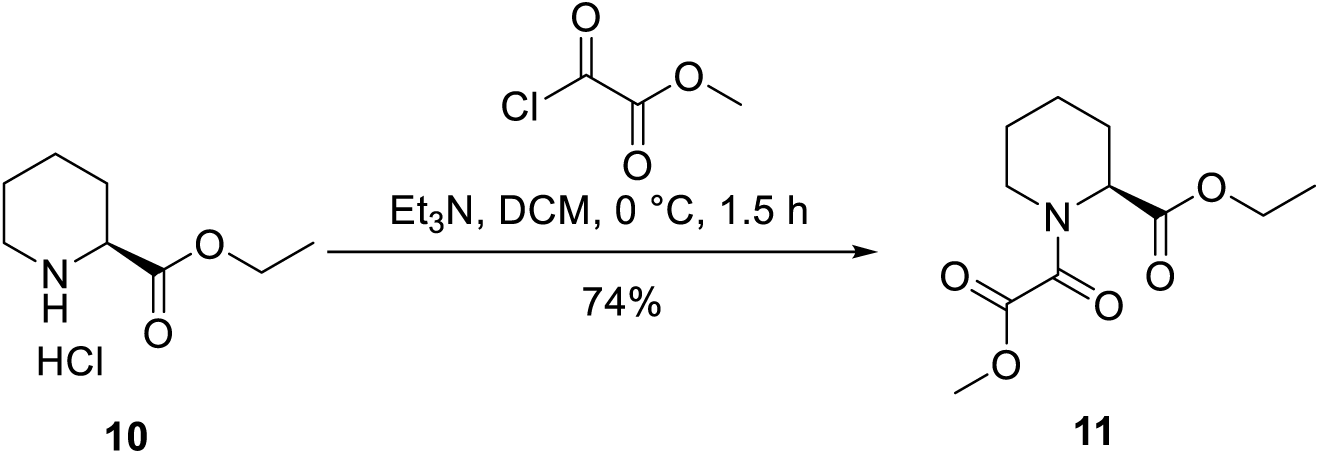

To a solution of ethyl (S)-piperidine-2-carboxylate hydrochloride (**10**) (350.0 g, 2.2 mol, 1.0 eq) in DCM (4.6 L) was added Et3N (888.8 g, 8.8 mol, 4.0 eq) at 0 °C. The mixture was stirred at this temperature for 15 min, then cooled to 0 °C and methyl 2-chloro-2-oxoacetate (296.5 g, 2.4 mol, 1.1 eq) in DCM (4 L) was added dropwise. The reaction mixture was stirred at 0 °C for 1.5 h. The mixture was diluted with water (8 L), extracted with DCM (8 L x 3), dried over Na2SO4, filtered and concentrated. The residue was purified with silica gel chromatography (Petroleum ether: EtOAc = 5: 1) to afford ethyl (S)-1-(2-methoxy-2-oxoacetyl)piperidine-2-carboxylate (**11**) (400.0 g, 74% yield) as a yellow solid.

**^1^H NMR (300 MHz, DMSO-*d6*):** *δ* ppm 5.02-5.00 (m, 0.6H), 4.55-4.53 (m, 0.3H), 4.20-4.13 (m, 2H), 3.83-3.77 (m, 3H), 3.52-3.47 (m, 1H), 3.25-3.13 (m, 1H), 2.17-2.12 (m, 1H), 1.70-1.63 (m, 3H), 1.41-1.25 (m, 2H), 1.20 (t, *J* = 7.1 Hz, 3H).

### Ethyl (S)-1-(3,3-dimethyl-2-oxopentanoyl)piperidine-2-carboxylate (12)

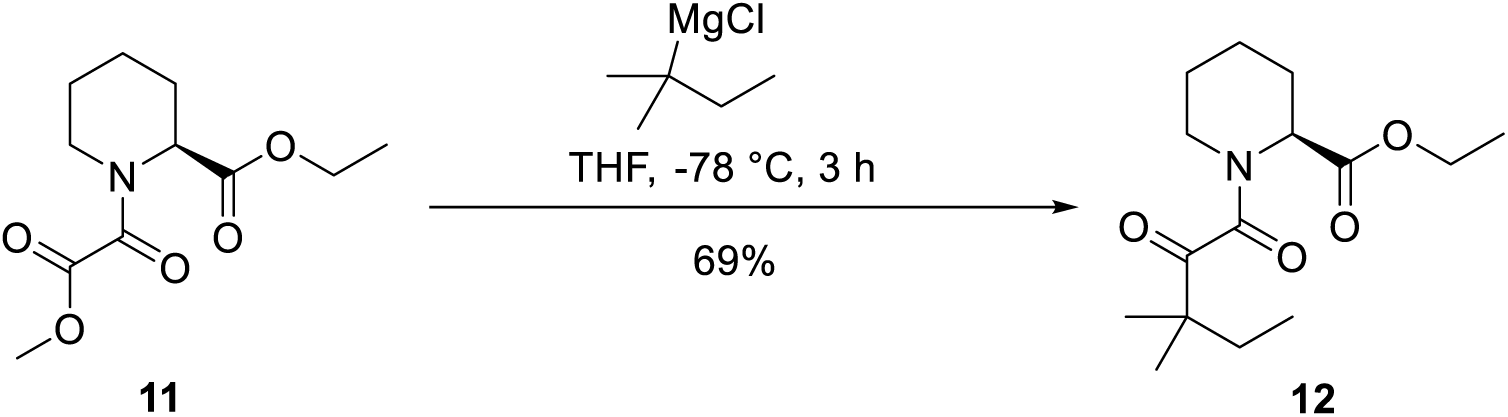

**Tert-pentylmagnesium chloride:** To a mixture of Mg (23.0 g, 946.5 mmol, 1.0 eq) and I2 (catalytic amount) was added THF (280 mL) under an argon atmosphere. Several drops of 2-chloro-2-methylbutane were added to the mixture. The mixture was heated to reflux for 1 h before 2-chloro-2-methylbutane (100.0 g, 938.1 mmol, 1.0 eq) in THF (240 mL) was added dropwise to the mixture. Then the mixture was stirred at reflux. After 6 h, the mixture was cooled to room temperature to afford tert-pentylmagnesium chloride (520 mL) as a dark brown solution, which was directly used to the next step.

To a solution of ethyl (S)-1-(2-methoxy-2-oxoacetyl)piperidine-2-carboxylate (**11**) (150.0 g, 616.6 mmol, 1.0 eq) in THF (1700 mL) was added tert-pentylmagnesium chloride (924.9 mmol, 1.5 eq, as a THF solution) at −78 °C under N2 atmosphere. The reaction mixture was stirred at −78 °C for 3 h. The reaction mixture was quenched with NH4Cl (1.7 L) and extracted with EtOAc (1.7 L x 3). The combined organic layer was washed with brine (1.7 L x 3), dried over Na2SO4, filtered and concentrated. The residue was purified with silica gel chromatography (Petroleum ether: EtOAc = 5: 1) to afford ethyl (S)-1-(3,3-dimethyl-2-oxopentanoyl)piperidine-2-carboxylate (**12**) (120.0 g, 69% yield) as a yellow solid.

**^1^H NMR (300 MHz, CDCl3):** *δ* ppm 5.23-5.21 (m, 1H), 4.25-4.14 (m, 2H), 3.39-3.34 (m, 1H), 3.24-3.15 (m, 1H), 2.32-2.24 (m, 1H), 1.78-1.59 (m, 7H), 1.29-1.24 (m, 3H), 1.22 (s, 3H), 1.18-1.17 (m, 3H), 0.89-0.84 (m, 3H).

### (S)-1-(3,3-dimethyl-2-oxopentanoyl)piperidine-2-carboxylic acid (13)

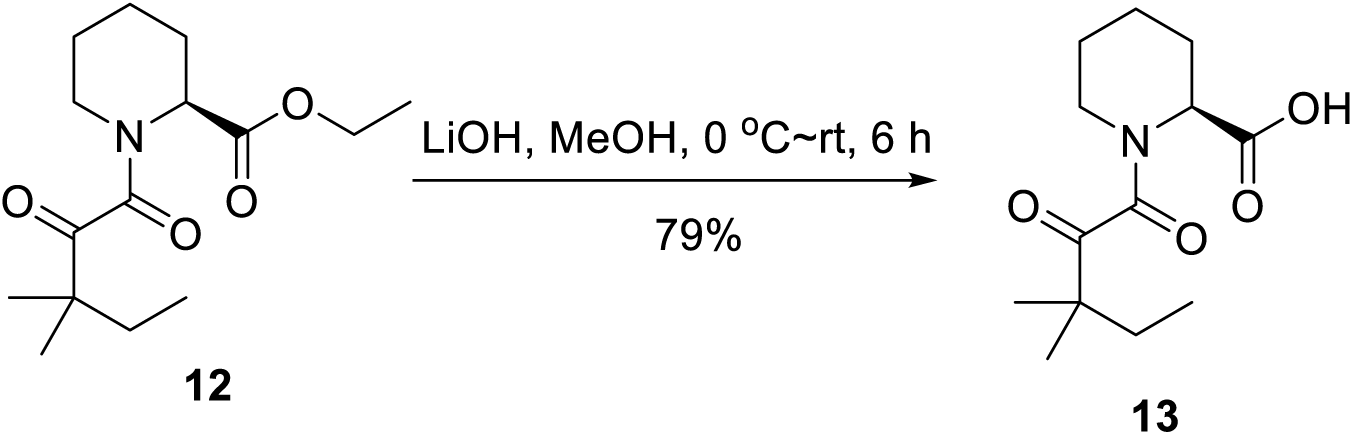

To a solution of ethyl (S)-1-(3,3-dimethyl-2-oxopentanoyl)piperidine-2-carboxylate (**12**) (120.0 g, 423.5 mmol, 1.0 eq) in MeOH (1.5 L) and H2O (47.6 mL, 2.6 mol, 5.0 eq) was added LiOH·H2O (26.7 g, 635.1 mmol, 1.5 eq) at 0 °C. The mixture was stirred at room temperature for 6 h and then concentrated. The residue was acidified with 10% aqueous HCl (100 mL) and the white solid was filtered and dried to afford (S)-1-(3,3-dimethyl-2-oxopentanoyl)piperidine-2-carboxylic acid (**13**) (85.0 g, 79%) as a white solid, which was directly used to the next step without further purification.

**^1^H NMR (400 MHz, CDCl3):** *δ* ppm 5.33 (d, *J* = 5.5 Hz, 1H), 3.42 (d, *J* = 13.3 Hz, 1H), 3.28-3.21 (m, 1H), 2.34 (d, *J* = 13.7 Hz, 1H), 1.83-1.61 (m, 5H), 1.58-1.43 (m, 2H), 1.23 (s, 3H), 1.20-0.87 (m, 3H), 0.89 (t, *J* = 7.4 Hz, 3H).

**LCMS:** 256.2 ([M+H]^+^).

### Methyl (R)-2-((S)-1-(3,3-dimethyl-2-oxopentanoyl)piperidine-2-carboxamido)-4-(4-methoxyphenyl)butanoate (14)

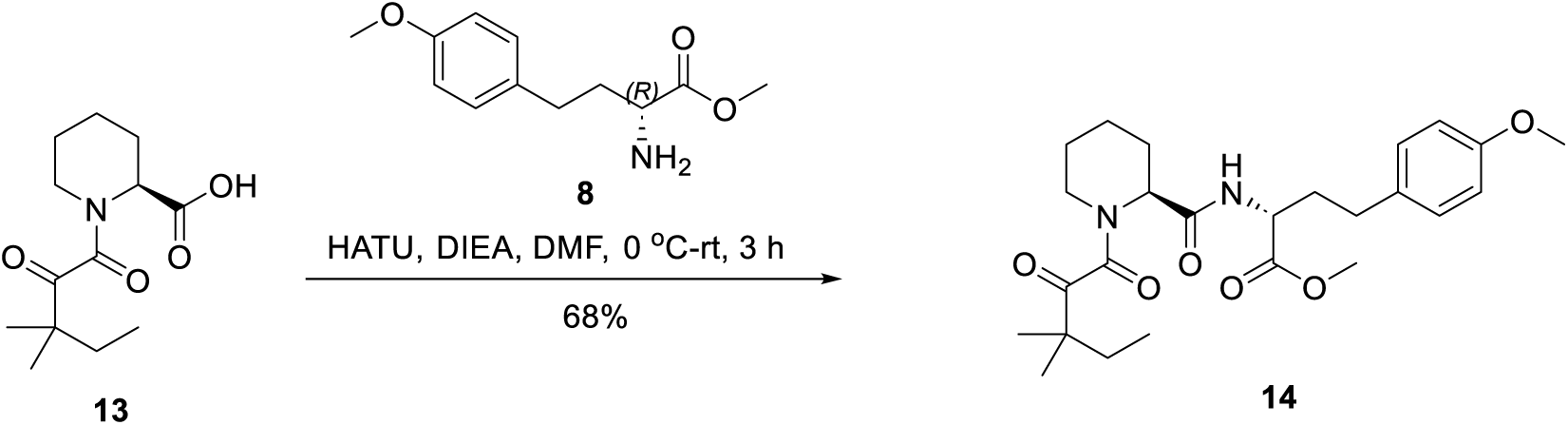

To a solution of (S)-1-(3,3-dimethyl-2-oxopentanoyl)piperidine-2-carboxylic acid (**13**) (61.0 g, 238.9 mmol, 1.0 eq) in DMF (610 mL) was added DIPEA (92.5 g, 716.8 mmol, 3.0 eq) and methyl (R)-2-amino-4-(4-methoxyphenyl)butanoate (**8**) (53.3 g, 238.9 mmol, 1.0 eq). HATU (136.2 g, 358.4 mmol, 1.8 eq) was added in portions to the mixture at 0 °C and then the reaction was stirred at room temperature for 3 h. The reaction was quenched with H2O (2 L), extracted with EtOAc (1 L x 3), washed with H2O (1 L x 3) and brine (1 L x 3), dried over Na2SO4, filtered and concentrated. The residue was purified by silica gel chromatography (Petroleum ether: EtOAc = 5: 1) to afford methyl (R)-2-((S)-1-(3,3-dimethyl-2-oxopentanoyl)piperidine-2-carboxamido)-4-(4-methoxyphenyl)butanoate (**14**) (75.0 g, 68% yield) as a yellow oil.

**^1^H NMR (300 MHz, CDCl3):** *δ* ppm 7.13 (d, *J* = 8.5 Hz, 2H), 7.07 (d, *J* = 8.5 Hz, 0.4H), 6.83-6.80 (m, 2H), 6.45 (d, *J* = 7.8 Hz, 0.6H), 5.17-5.16 (m, 0.6H), 4.62-4.55 (m, 1H), 4.53-4.51 (m, 0.4H), 3.99 (d, *J* = 5.2 Hz, 0.4H), 3.77 (s, 3H), 3.70 (s, 3H), 3.39-3.20 (m, 1H), 2.84-2.60 (m, 1H), 2.60-2.55 (m, 1H), 2.46-2.35 (m, 1H), 2.22-2.09 (m, 1H), 2.04-1.87 (m, 1H), 1.87-1.83 (m, 0.6H), 1.77-1.63 (m, 5H), 1.56-1.40 (m, 2H), 1.27-1.21 (m, 6H), 0.91 (t, *J* = 7.5 Hz, 3H).

**LCMS:** 461.2 ([M+H]^+^).

### (R)-2-((S)-1-(3,3-dimethyl-2-oxopentanoyl)piperidine-2-carboxamido)-4-(4-methoxyphenyl)butanoic acid (15)

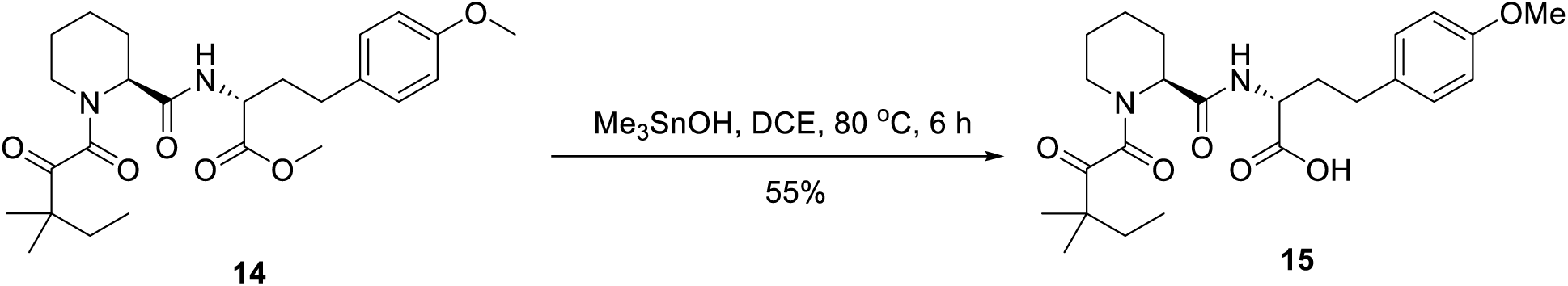

To a solution of methyl (R)-2-((S)-1-(3,3-dimethyl-2-oxopentanoyl)piperidine-2-carboxamido)-4-(4-methoxyphenyl)butanoate (**14**) (75.0 g, 162.8 mmol, 1.0 eq) in DCE (750 mL) was added Me3SnOH (88.3 g, 488.5 mmol, 3.0 eq). The reaction was stirred at 80 °C for 6 h. The mixture was cooled to room temperature and washed with sat. NH4Cl (700 mL x 3). The aqueous layer was extracted with DCM (500 mL x 3), and the combined organic phase was washed with brine (500 mL x 3), dried over Na2SO4, filtered and concentrated. The residue was purified by silica gel chromatography (Petroleum ether: EtOAc = 5: 1) to afford (R)-2-((S)-1-(3,3-dimethyl-2-oxopentanoyl)piperidine-2-carboxamido)-4-(4-methoxyphenyl)butanoic acid (**15**) (40.0 g, 55% yield) as a yellow solid.

**^1^H NMR (300 MHz, DMSO-*d6*):** *δ* ppm 12.6 (s, 1H), 8.30-8.24 (m, 1H), 7.08 (d, *J* = 8.5 Hz, 2H), 6.84 (d, *J* = 8.5 Hz, 2H), 5.02 (d, *J* = 4.4 Hz, 0.6H), 4.25 (d, *J* = 13.7 Hz, 0.3H), 4.18-4.11 (m, 1H), 3.72 (s, 3H), 3.56-3.48 (m, 1H), 3.20 (d, *J* = 12.6 Hz, 1H), 2.61-2.51 (m, 1H), 2.22-2.18 (m, 1H), 2.07-1.85 (m, 2H), 1.69-1.53 (m, 5H), 1.45-1.23 (m, 3H), 1.14-1.11 (m, 6H), 0.80 (t, *J* = 7.4 Hz, 3H).

**LCMS:** 447.2 ([M+H]^+^).

### Synthetic scheme for compound 3

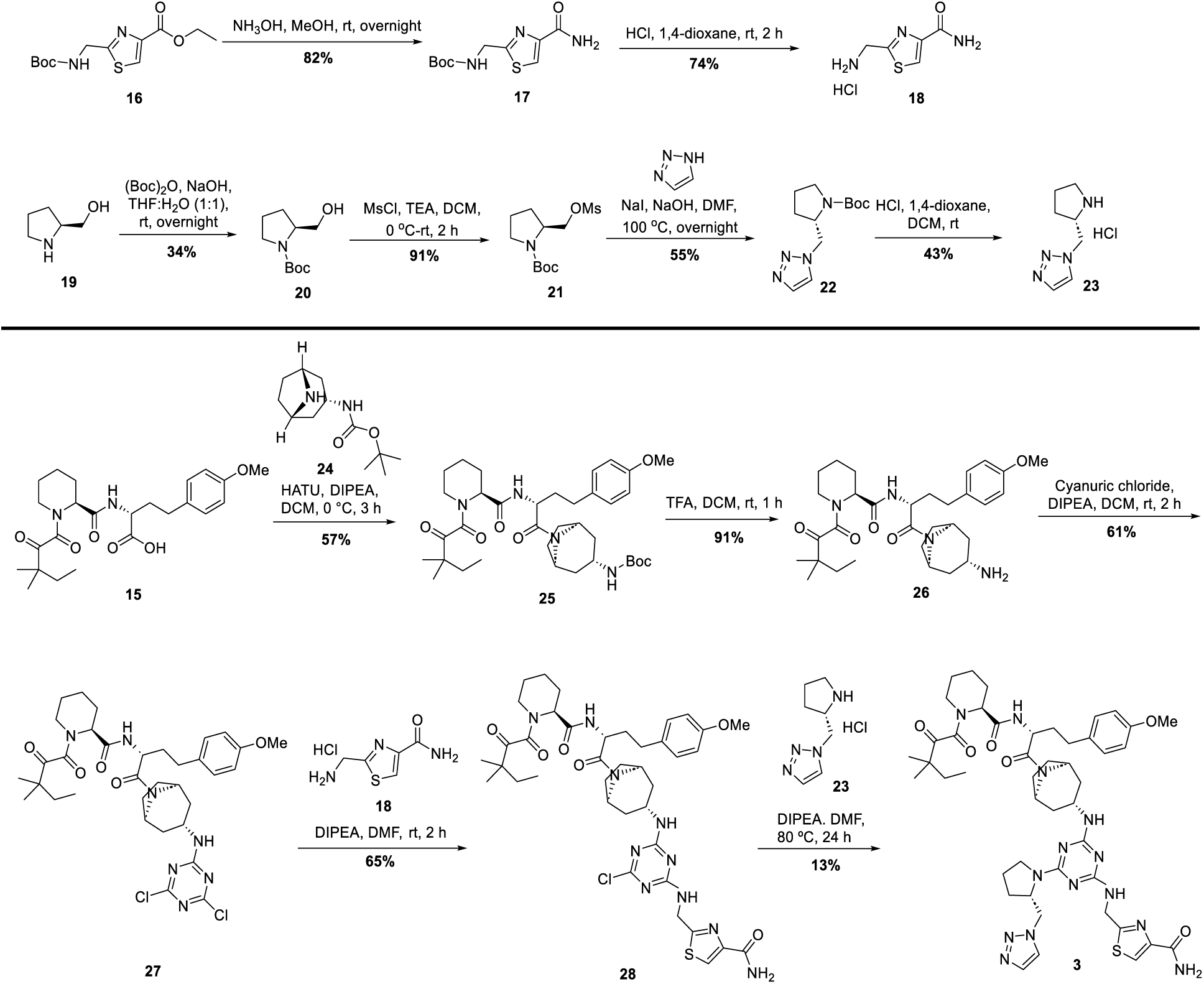

### Tert-butyl ((4-carbamoylthiazol-2-yl)methyl)carbamate (17)

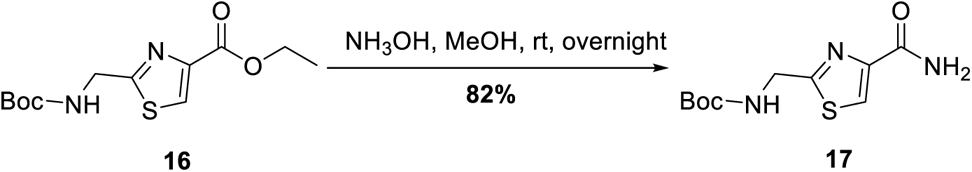

To a solution of ethyl 2-(((tert-butoxycarbonyl)amino)methyl)thiazole-4-carboxylate (**16**) (1.0 g, 3.5 mmol, 1.0 eq) in MeOH (35 mL) was added NH3OH (18 mL) at room temperature. The mixture was stirred at room temperature overnight, then concentrated to give tert-butyl ((4-carbamoylthiazol-2-yl)methyl)carbamate

1. (**17**) (740.0 mg, 82% yield) as a yellow solid, which was used for the next step without further purification.

**LCMS:** 258.0 ([M+H]^+^).

### 2-(aminomethyl)thiazole-4-carboxamide hydrochloride (18)

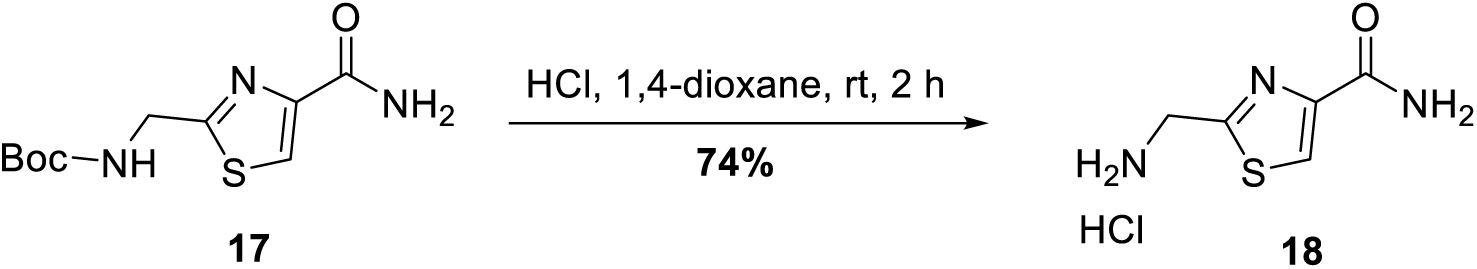

To a solution of tert-butyl ((4-carbamoylthiazol-2-yl)methyl)carbamate (**17**) (740.0 mg, 2.9 mmol, 1.0 eq) in DCM (8 mL) was added 4M HCl-dioxane (8 mL) at 0 °C. The mixture was stirred at room temperature for 2 h, then concentrated to give 2-(aminomethyl)thiazole-4-carboxamide hydrochloride (**18**) (410.0 mg, 74% yield) as a white solid.

**^1^H NMR (400 MHz, DMSO-*d6*):** *δ* ppm 8.88 (s, 3H), 8.31 (s, 1H), 7.79 (s, 1H), 7.68 (s, 1H), 4.43-4.42 (m, 2H).

**LCMS:** 157.9 ([M-HCl+H]^+^).

### tert-butyl (S)-2-(hydroxymethyl)pyrrolidine-1-carboxylate (20)

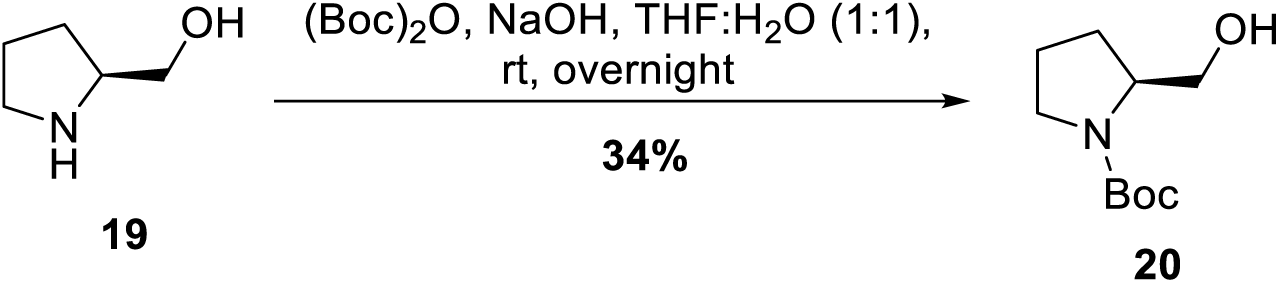

To a solution of (S)-pyrrolidin-2-ylmethanol (**19**) (3.0 g, 29.6 mmol, 1.0 eq) in THF (30 mL) and H2O (20 mL) was added NaOH (1.5 g, 38.5 mmol, 1.3 eq) and (Boc)2O (7.7 g, 35.5 mmol, 1.2 eq) at 0 °C. The mixture was stirred at room temperature overnight. The mixture was concentrated and H2O (30 mL) was added. The mixture was extracted with DCM (20 mL x 3), washed with brine (20 mL x 3), dried over Na2SO4, filtered and concentrated. The crude was purified by silica gel chromatography (Petroleum ether: EtOAc = 1: 1) to afford tert-butyl (S)-2-(hydroxymethyl)pyrrolidine-1-carboxylate (**20**) (2.0 g, 34% yield) as a clear oil.

**LCMS:** 146.1 ([M-tBu+H]^+^).

### tert-butyl (S)-2-(((methylsulfonyl)oxy)methyl)pyrrolidine-1-carboxylate (21)

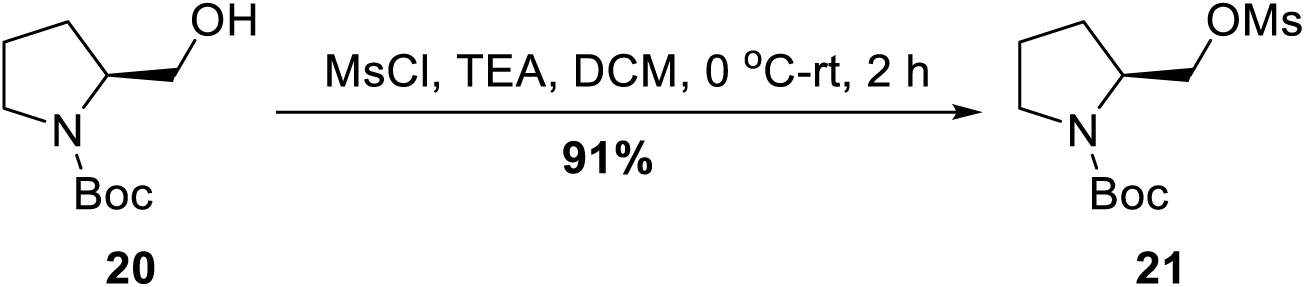

To a solution of tert-butyl (S)-2-(hydroxymethyl)pyrrolidine-1-carboxylate (**20**) (1.5 g, 7.5 mmol, 1.0 eq) in DCM (10 mL) was added TEA (2.3 g, 22.5 mmol, 3.0 eq) at 0 °C. MsCl (1.3 g, 11.3 mmol, 1.5 eq) in DCM (10 mL) was added dropwise to the mixture at 0 °C. The mixture was stirred at room temperature for 2 h. After completed, the mixture was quenched with H2O (10 mL) and extracted with DCM (10 mL x 3), washed with brine (10 mL x 3), dried over Na2SO4, filtered and concentrated. The crude was purified by silica gel chromatography (Petroleum ether: EtOAc = 1: 1) to give tert-butyl (S)-2-(((methylsulfonyl)oxy)methyl)pyrrolidine-1-carboxylate (**21**) (1.9 g, 91% yield) as a clear oil.

**LCMS:** 224.1 ([M-56+H]^+^).

### tert-butyl (S)-2-((1H-1,2,3-triazol-1-yl)methyl)pyrrolidine-1-carboxylate (22)

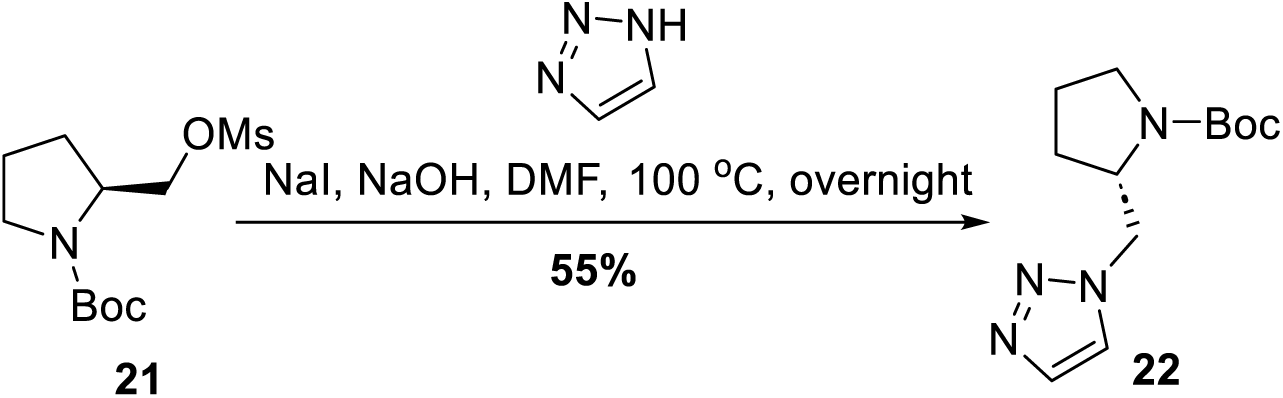

To a solution of tert-butyl (S)-2-(((methylsulfonyl)oxy)methyl)pyrrolidine-1-carboxylate (**21**) (1.0 g, 3.6 mmol, 1.0 eq) in DMF (10 mL) was added NaOH (216 mg, 5.4 mmol, 1.5 eq), NaI (540 mg, 3.6 mmol, 1.0 eq) and 1,2,3-1*H*-Triazole (347 mg, 5.0 mmol, 1.4 eq). The mixture was stirred at 100 °C overnight. H2O (50 mL) was added and the mixture extracted with EtOAc (30 mL x 3), washed with H2O (50 mL x 3) and brine (50 mL x 3), dried over Na2SO4, concentrated and purified with silica gel chromatography (Petroleum ether: EtOAc = 1: 1) to give tert-butyl (S)-2-((1H-1,2,3-triazol-1-yl)methyl)pyrrolidine-1-carboxylate (**22**) (500.0 mg, 55% yield) as a white solid.

**^1^H NMR (300 MHz, CDCl3):** *δ* ppm 7.70 (s, 1H), 7.52-7.47 (m, 2H), 4.70-4.49 (m, 2H), 4.11-4.09 (m, 1H), 3.37-3.09 (m, 2H), 1.92 (s, 2H), 1.71 (s, 1H), 1.49 (s, 9H), 1.42-1.34 (m, 1H).

**LCMS:** 253.2 ([M+H]^+^).

### (S)-1-(pyrrolidin-2-ylmethyl)-1H-1,2,3-triazole hydrochloride (23)

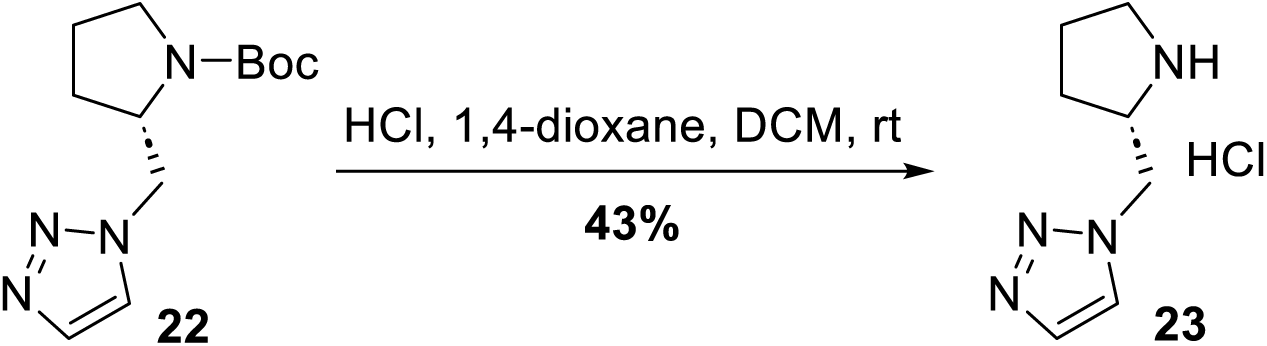

To a solution of tert-butyl (S)-2-((1H-1,2,3-triazol-1-yl)methyl)pyrrolidine-1-carboxylate (**22**) (250.0 mg, 1.0 mmol, 1.0 eq) in DCM (2 mL) was added 4M HCl-dioxane (2 mL) at 0 °C. The mixture was stirred at room temperature for 1.5 h, then concentrated and dried by lyophilization to give (S)-1-(pyrrolidin-2-ylmethyl)-1H-1,2,3-triazole hydrochloride (**23**) (80.0 mg, 43% yield) as a white solid.

**^1^H NMR (400 MHz, DMSO-*d6*):** *δ* ppm 9.25 (s, 1H), 9.00 (s, 1H), 8.19 (s, 1H), 7.81 (s, 1H), 4.80-4.67 (m, 2H), 4.00-3.96 (m, 1H), 3.26-3.14 (m, 2H), 2.14-2.06 (m, 1H), 1.98-1.85 (m, 2H), 1.73-1.64 (m, 1H).

**LCMS:** 153.2 ([M-HCl+H]^+^).

### tert-butyl ((1R,3S,5S)-8-((R)-2-((S)-1-(3,3-dimethyl-2-oxopentanoyl)piperidine-2-carboxamido)-4-(4-methoxyphenyl)butanoyl)-8-azabicyclo[3.2.1]octan-3-yl)carbamate (25)

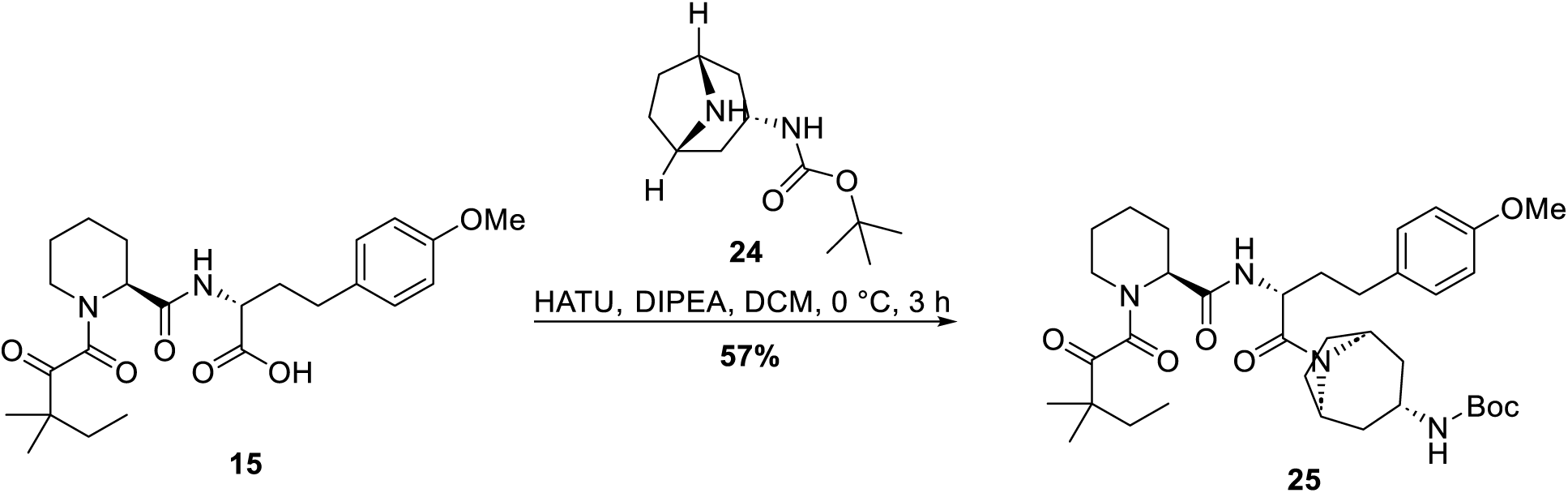

To a solution of (R)-2-((S)-1-(3,3-dimethyl-2-oxopentanoyl)piperidine-2-carboxamido)-4-(4-methoxyphenyl)butanoic acid (**15**) (520.0 mg, 1.2 mmol, 1.0 eq) in DCM (130 mL) was added DIPEA (464.4 mg, 3.6 mmol, 3.0 eq), tert-butyl ((1R,3r,5S)-8-azabicyclo[3.2.1]octan-3-yl)carbamate (**24**) (316.4 mg, 1.4 mmol, 1.2 eq) and HATU (532.0 mg, 1.4 mmol, 1.2 eq) at 0 °C. The mixture was stirred at this temperature for 3 h and then concentrated to give a residue. The residue was purified by silica gel chromatography (Petroleum ether: EtOAc = 5: 1) to afford tert-butyl ((1R,3S,5S)-8-((R)-2-((S)-1-(3,3-dimethyl-2-oxopentanoyl)piperidine-2-carboxamido)-4-(4-methoxyphenyl)butanoyl)-8-azabicyclo[3.2.1]octan-3-yl)carbamate (**25**) (430.0 mg, 57% yield) as a white solid.

**LCMS:** 655.4 ([M+H]^+^).

### (S)-N-((R)-1-((1R,3S,5S)-3-amino-8-azabicyclo[3.2.1]octan-8-yl)-4-(4-methoxyphenyl)-1-oxobutan-2-yl)-1-(3,3-dimethyl-2-oxopentanoyl)piperidine-2-carboxamide (26)

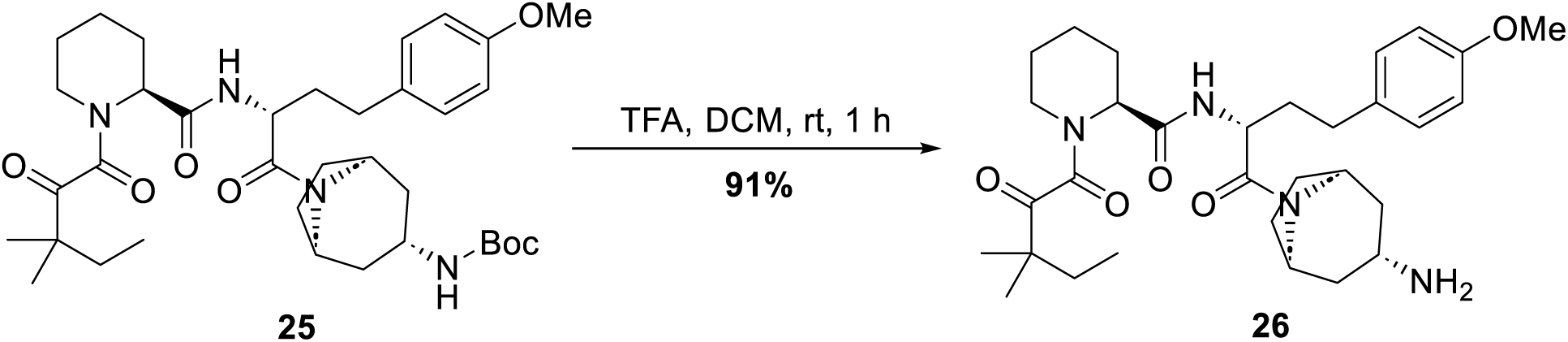

To a solution of tert-butyl ((1R,3S,5S)-8-((R)-2-((S)-1-(3,3-dimethyl-2-oxopentanoyl)piperidine-2-carboxamido)-4-(4-methoxyphenyl)butanoyl)-8-azabicyclo[3.2.1]octan-3-yl)carbamate (**25**) (430.0 mg, 0.7 mmol, 1.0 eq) in DCM (4.5 mL) was added TFA (1.5 mL). The mixture was stirred at room temperature for 1 h, then concentrated and dried by lyophilization to afford (S)-N-((R)-1-((1R,3S,5S)-3-amino-8-azabicyclo[3.2.1]octan-8-yl)-4-(4-methoxyphenyl)-1-oxobutan-2-yl)-1-(3,3-dimethyl-2-oxopentanoyl)piperidine-2-carboxamide (**26**) (353.0 mg, 91% yield) as a white solid.

**^1^H NMR (400 MHz, CDCl3):** *δ* ppm 8.09-7.97 (m, 2H), 7.17-7.06 (m, 2H), 6.82 (d, *J* = 8.0 Hz, 2H), 5.23-5.13 (m, 1H), 4.69-4.46 (m, 2H), 4.07 (s, 1H), 3.77-3.75 (m, 3H), 3.70 (s, 2H), 3.57-3.42 (m, 1H), 3.39-3.28 (m, 1H), 3.11-2.80 (m, 1H), 2.78-2.20 (m, 4H), 2.08 (s, 2H), 2.00-1.85 (m, 4H), 1.81-1.66 (m, 6H), 1.54-1.38 (m, 3H), 1.27-1.21 (m, 6H), 0.91-0.86 (m, 3H).

**LCMS:** 555.2 ([M+H]^+^).

### (S)-N-((R)-1-((1R,3S,5S)-3-((4,6-dichloro-1,3,5-triazin-2-yl)amino)-8-azabicyclo[3.2.1]octan-8-yl)-4-(4-methoxyphenyl)-1-oxobutan-2-yl)-1-(3,3-dimethyl-2-oxopentanoyl)piperidine-2-carboxamide (27)

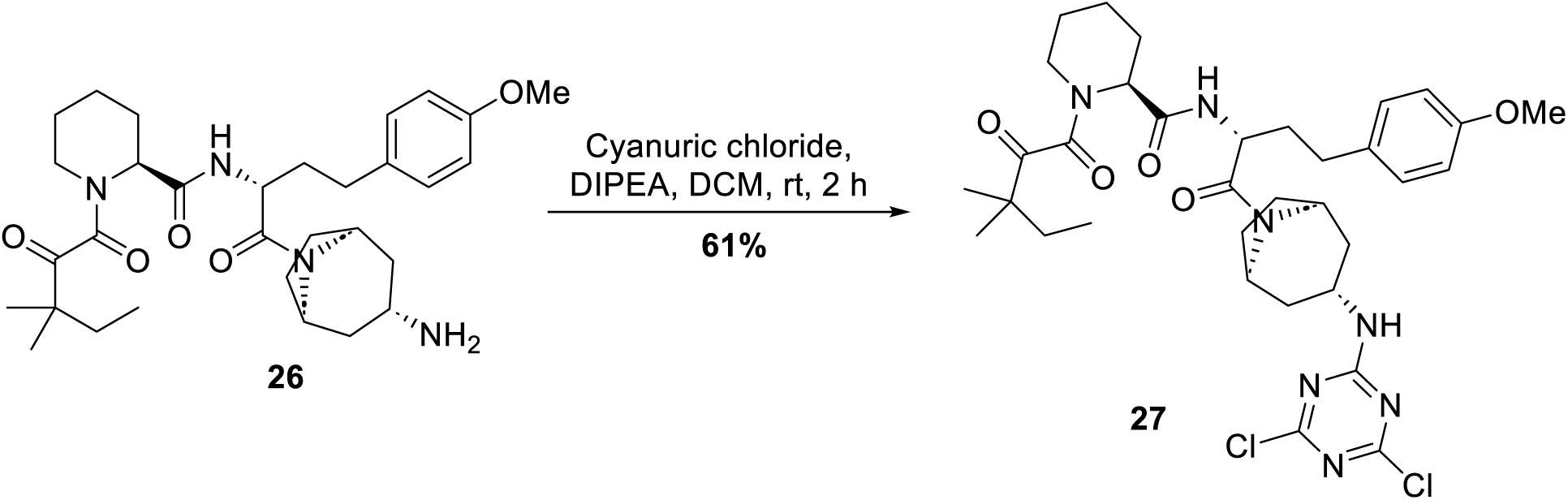

To a solution of (S)-N-((R)-1-((1R,3S,5S)-3-amino-8-azabicyclo[3.2.1]octan-8-yl)-4-(4-methoxyphenyl)-1-oxobutan-2-yl)-1-(3,3-dimethyl-2-oxopentanoyl)piperidine-2-carboxamide (**26**) (270.0 mg, 0.5 mmol, 1.0 eq) in DCM (3 mL) was added DIPEA (322.2 mg, 2.5 mmol, 5.0 eq). Cyanuric chloride (110.4 mg, 0.6 mmol, 1.2 eq) was added to the mixture at 0 °C. The mixture was stirred at this temperature for 2 h and then concentrated to give a residue. The residue was purified by silica gel chromatography (Petroleum ether: EtOAc = 3: 1) to give (S)-N-((R)-1-((1R,3S,5S)-3-((4,6-dichloro-1,3,5-triazin-2-yl)amino)-8-azabicyclo[3.2.1]octan-8-yl)-4-(4-methoxyphenyl)-1-oxobutan-2-yl)-1-(3,3-dimethyl-2-oxopentanoyl)piperidine-2-carboxamide (**27**) (210.0 mg, 61% yield) as a yellow oil.

**LCMS:** 702.2 ([M+H]^+^).

### 2-(((4-chloro-6-(((1R,3S,5S)-8-((R)-2-((S)-1-(3,3-dimethyl-2-oxopentanoyl)piperidine-2-carboxamido)-4-(4-methoxyphenyl)butanoyl)-8-azabicyclo[3.2.1]octan-3-yl)amino)-1,3,5-triazin-2-yl)amino)methyl)thiazole-4-carboxamide (28)

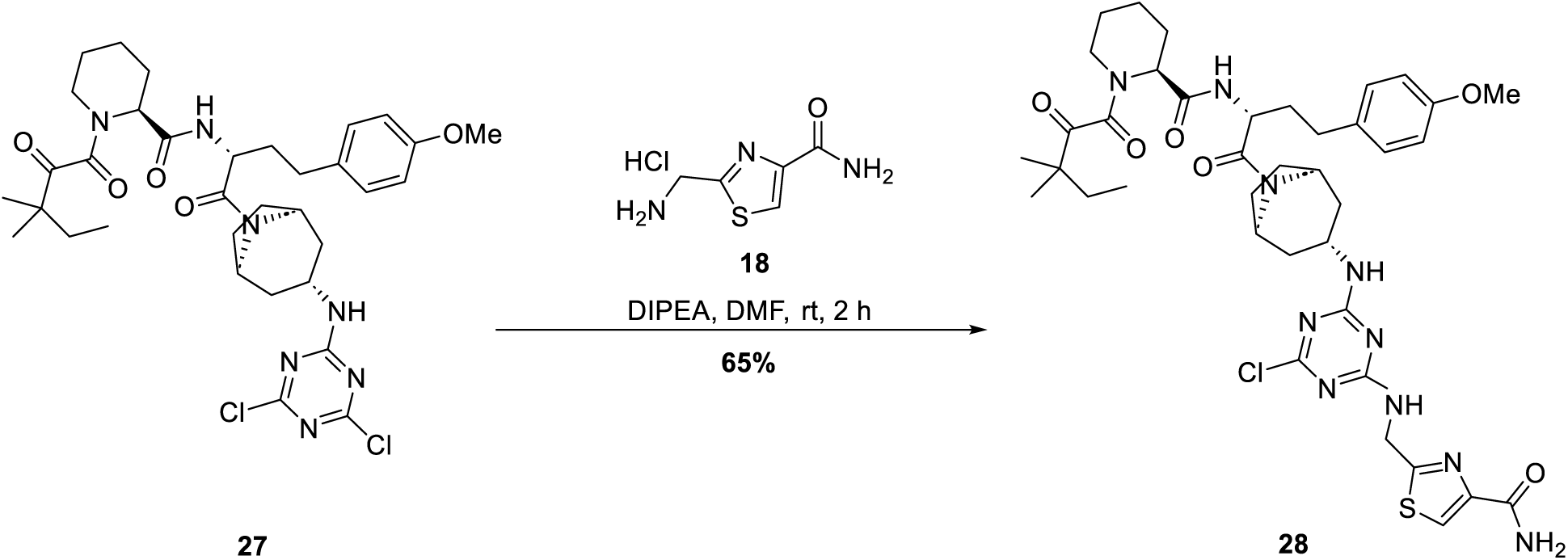

To a solution of (S)-N-((R)-1-((1R,3S,5S)-3-((4,6-dichloro-1,3,5-triazin-2-yl)amino)-8-azabicyclo[3.2.1]octan-8-yl)-4-(4-methoxyphenyl)-1-oxobutan-2-yl)-1-(3,3-dimethyl-2-oxopentanoyl)piperidine-2-carboxamide (**27**) (210.0 mg, 0.30 mmol, 1.0 eq) in DMF (2.5 mL) was added DIPEA (193.5 mg, 1.5 mmol, 5.0 eq) and 2-(aminomethyl)thiazole-4-carboxamide hydrochloride (**18**) (81.3 mg, 0.42 mmol, 1.4 eq). The mixture was stirred at room temperature for 2 h, then water (10 mL) was added and the mixture extracted with EtOAc (10 mL x 3). The combined organic phase was washed with brine (10 mL x 3), dried over Na2SO4, filtered and concentrated to give 2-(((4-chloro-6-(((1R,3S,5S)-8-((R)-2-((S)-1-(3,3-dimethyl-2-oxopentanoyl)piperidine-2-carboxamido)-4-(4-methoxyphenyl)butanoyl)-8-azabicyclo[3.2.1]octan-3-yl)amino)-1,3,5-triazin-2-yl)amino)methyl)thiazole-4-carboxamide (**28**) (160.0 mg, 65% yield) as a yellow oil.

**LCMS:** 823.2 ([M+H]^+^).

### 2-(((4-((S)-2-((1H-1,2,3-triazol-1-yl)methyl)pyrrolidin-1-yl)-6-(((1R,3S,5S)-8-((R)-2-((S)-1-(3,3-dimethyl-2-oxopentanoyl)piperidine-2-carboxamido)-4-(4-methoxyphenyl)butanoyl)-8-azabicyclo[3.2.1]octan-3-yl)amino)-1,3,5-triazin-2-yl)amino)methyl)thiazole-4-carboxamide (3)

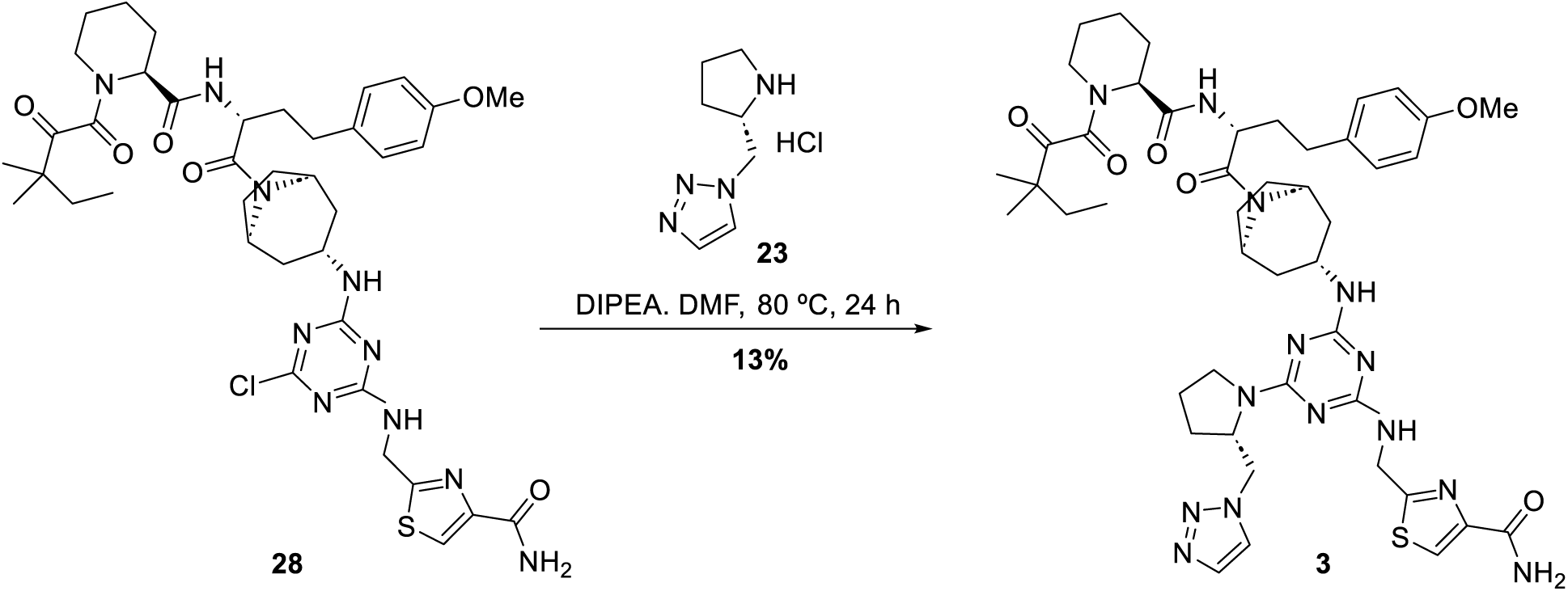

To a solution of 2-(((4-chloro-6-(((1R,3S,5S)-8-((R)-2-((S)-1-(3,3-dimethyl-2-oxopentanoyl)piperidine-2-carboxamido)-4-(4-methoxyphenyl)butanoyl)-8-azabicyclo[3.2.1]octan-3-yl)amino)-1,3,5-triazin-2-yl)amino)methyl)thiazole-4-carboxamide (**28**) (160.0 mg, 0.19 mmol, 1.0 eq) in DMF (2 mL) was added DIPEA (125.1 mg, 0.97 mmol, 5.0 eq) and (S)-1-(pyrrolidin-2-ylmethyl)-1H-1,2,3-triazole hydrochloride (**23**) (54.7 mg, 0.29 mmol, 1.5 eq). The mixture was stirred at 80 °C for 24 h. Water (10 mL) was added and the mixture extracted with EtOAc (10 mL x 3). The combined organic phase was washed with brine (10 mL x 3), dried over Na2SO4 and concentrated to give a residue. The residue was purified by Prep-TLC (DCM: MeOH = 10: 1) to give 2-(((4-((S)-2-((1H-1,2,3-triazol-1-yl)methyl)pyrrolidin-1-yl)-6-(((1R,3S,5S)-8-((R)-2-((S)-1-(3,3-dimethyl-2-oxopentanoyl)piperidine-2-carboxamido)-4-(4-methoxyphenyl)butanoyl)-8-azabicyclo[3.2.1]octan-3-yl)amino)-1,3,5-triazin-2-yl)amino)methyl)thiazole-4-carboxamide (**3**) (25 mg, 13% yield) as a white solid.

**^1^H NMR (400 MHz, DMSO-*d6*):** *δ* ppm 8.39 - 8.29 (m, 1H), 8.21 - 7.95 (m, 2H), 7.74 - 7.51 (m, 3H), 7.11- 7.08 (m, 2H), 6.85 (d, *J* = 7.6 Hz, 2H), 5.04 - 4.94 (m, 1H), 4.73 - 4.24 (m, 9H), 4.15 - 3.93 (m, 1H), 3.71- 3.69 (m, 3H), 3.64 - 3.54 (m, 1H), 3.29 (s, 3H), 3.26 - 3.12 (m, 1H), 2.26 - 2.08 (m, 1H), 2.10 - 1.19 (m, 23H), 1.18 - 1.02 (m, 6H), 0.84 - 0.71 (m, 3H).

**LCMS:** 939.1 ([M+H]^+^).

### Synthetic scheme for compound **4**

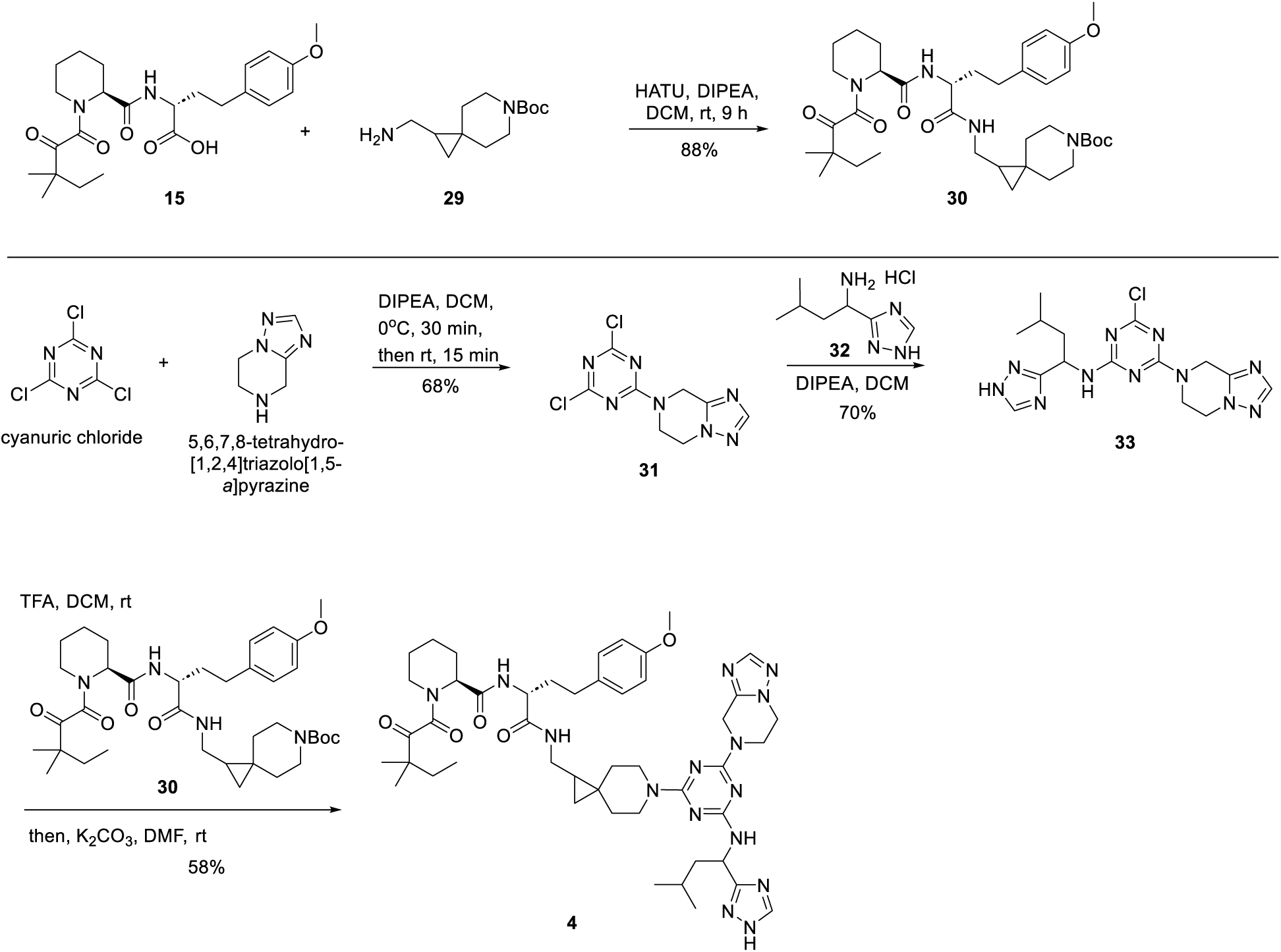

### tert-butyl 1-(((R)-2-((S)-1-(3,3-dimethyl-2-oxopentanoyl)piperidine-2-carboxamido)-4-(4-methoxyphenyl)butanamido)methyl)-6-azaspiro[2.5]octane-6-carboxylate (30)

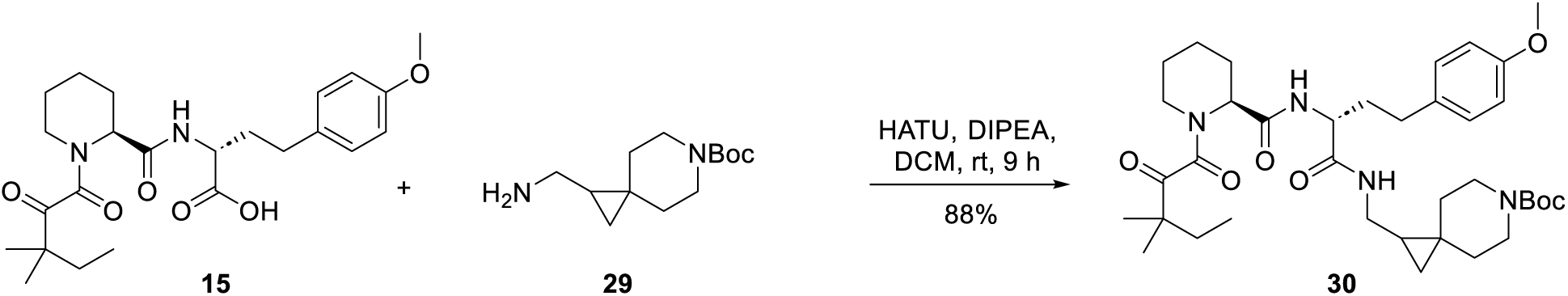

To a solution of (R)-2-((S)-1-(3,3-dimethyl-2-oxopentanoyl)piperidine-2-carboxamido)-4-(4-methoxyphenyl)butanoic acid (**15**) (75 mg, 168 μmol, 1.0 eq), tert-butyl 1-(aminomethyl)-6-azaspiro[2.5]octane-6-carboxylate (**29**) (101 mg, 420 μmol, 2.5 eq) in DCM (0.30 mL) was added DIPEA (117 μL, 672 μmol, 4 eq). HATU (96 mg, 252 μmol, 1.5 eq) was added in two portions (75% in first portion, then remaining added after 10 min). The reaction was stirred at room temperature for 7 h. A second portion of HATU (32 mg, 84 μmol, 0.5 eq) was added and the reaction stirred at room temperature for another 2 h. The reaction was quenched with saturated NH4Cl (aq), extracted with DCM (3x), washed with brine (1x), dried over Na2SO4, filtered and concentrated. The residue was purified by silica gel chromatography to afford tert-butyl 1-(((R)-2-((S)-1-(3,3-dimethyl-2-oxopentanoyl)piperidine-2-carboxamido)-4-(4-methoxyphenyl)butanamido)methyl)-6-azaspiro[2.5]octane-6-carboxylate (**30**) (99 mg, 88% yield).

**LCMS:** 569.4 ([M-Boc+H]^+^).

### 7-(4,6-dichloro-1,3,5-triazin-2-yl)-5,6,7,8-tetrahydro-[1,2,4]triazolo[1,5-a]pyrazine (31)

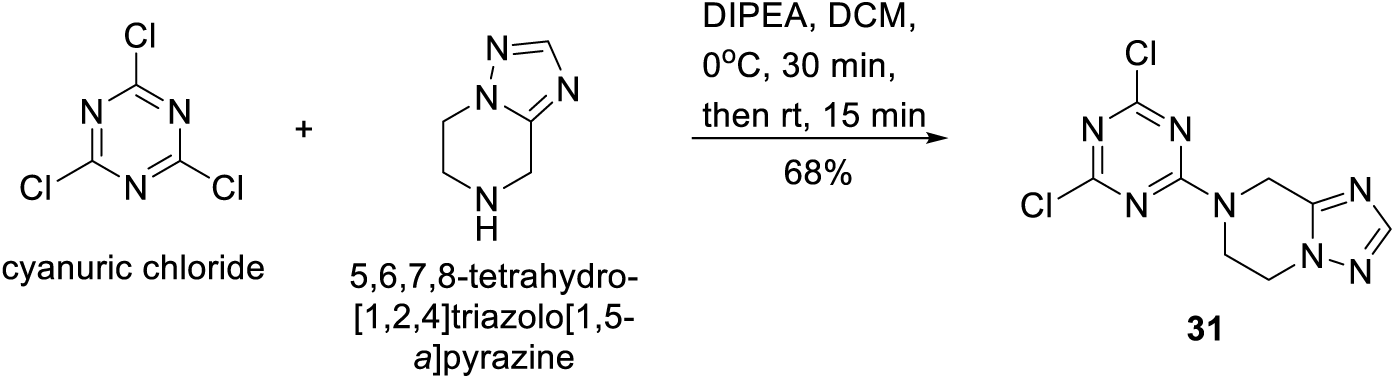

To a solution of cyanuric chloride (80 mg, 434 μmol, 1.0 eq) in DCM (1.2 mL) at 0 °C was added dropwise a solution of 5,6,7,8-tetrahydro-[1,2,4]triazolo[1,5-a]pyrazine (49 mg, 390 μmol, 0.9 eq) and DIPEA (227 μL, 1.30 mmol, 3.0 eq) in DCM (1.3 mL). The reaction mixture was stirred at 0 °C for 30 min and then at RT for 15 min. The reaction mixture was loaded and purified by silica gel chromatography (ethyl acetate/heptane 70-80%) to afford 7-(4,6-dichloro-1,3,5-triazin-2-yl)-5,6,7,8-tetrahydro-[1,2,4]triazolo[1,5-a]pyrazine (**31**) (80 mg, 68% yield).

**LCMS:** 271.9 ([M+H]^+^).

### 4-chloro-6-(5,6-dihydro-[1,2,4]triazolo[1,5-a]pyrazin-7(8H)-yl)-N-(3-methyl-1-(1H-1,2,4-triazol-3-yl)butyl)-1,3,5-triazin-2-amine (33)

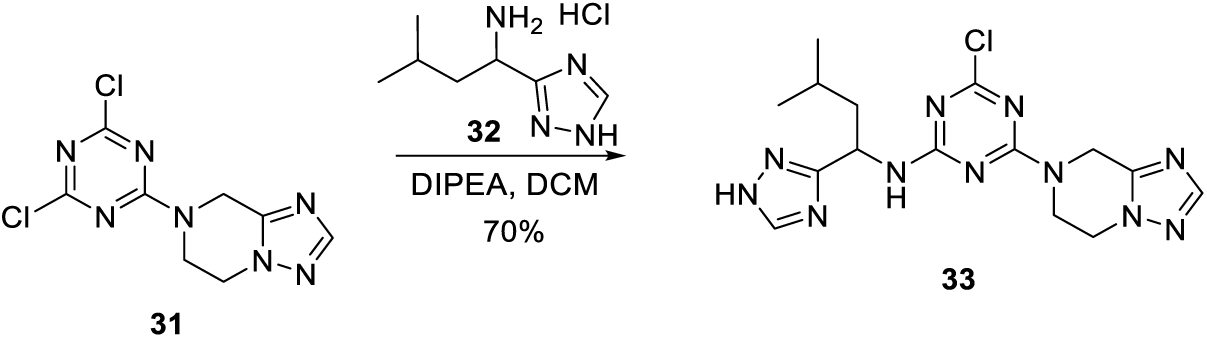

To a solution of 7-(4,6-dichloro-1,3,5-triazin-2-yl)-5,6,7,8-tetrahydro-[1,2,4]triazolo[1,5-a]pyrazine (**31**) (25 mg, 92 μmol, 1.0 eq) in DCM (0.6 mL) were added 3-methyl-1-(1H-1,2,4-triazol-3-yl)butan-1-amine hydrochloride (**32**) (21 mg, 110 μmol, 1.2 eq) and DIPEA (64 μL, 368 μmol, 4.0 eq) and the mixture stirred at room temperature for 30 minutes. The reaction mixture was loaded and purified by silica gel chromatography (MeOH/DCM 3-10%) to afford 4-chloro-6-(5,6-dihydro-[1,2,4]triazolo[1,5-a]pyrazin-7(8H)-yl)-N-(3-methyl-1-(1H-1,2,4-triazol-3-yl)butyl)-1,3,5-triazin-2-amine (**33**) (25 mg, 70% yield).

**LCMS:** 390.3 ([M+H]^+^).

### (2S)-N-((2R)-1-(((6-(4-(5,6-dihydro-[1,2,4]triazolo[1,5-a]pyrazin-7(8H)-yl)-6-((3-methyl-1-(1H-1,2,4-triazol-3-yl)butyl)amino)-1,3,5-triazin-2-yl)-6-azaspiro[2.5]octan-1-yl)methyl)amino)-4-(4-methoxyphenyl)-1-oxobutan-2-yl)-1-(3,3-dimethyl-2-oxopentanoyl)piperidine-2-carboxamide (4)

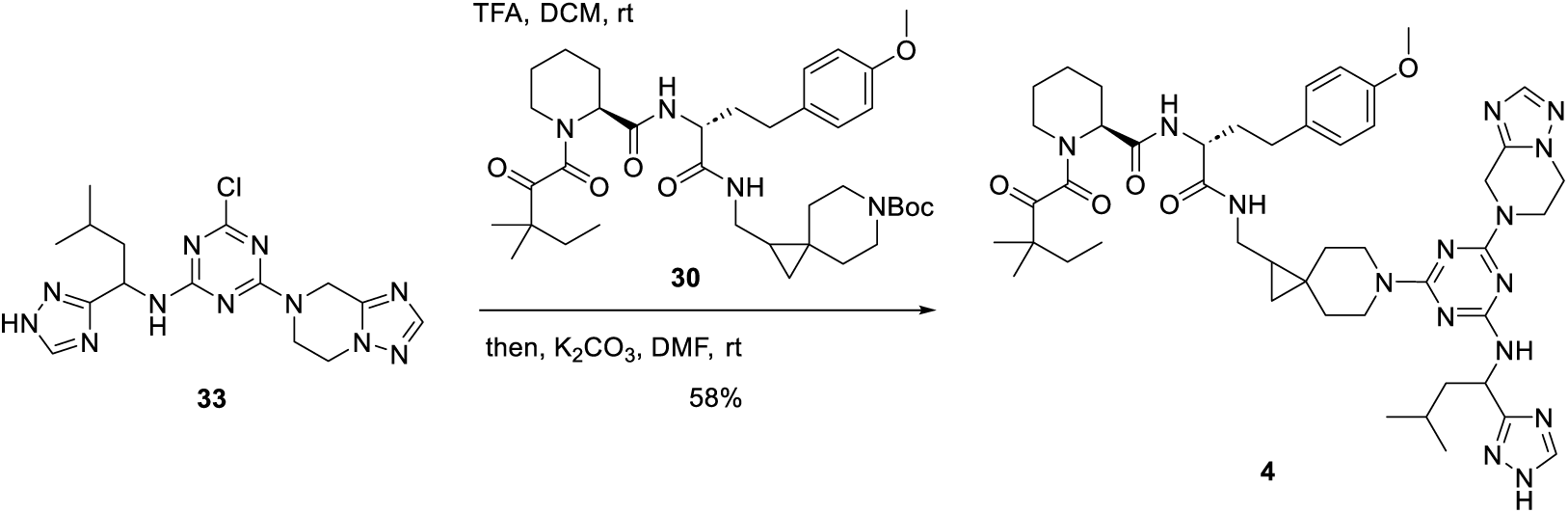

Tert-butyl1-(((R)-2-((S)-1-(3,3-dimethyl-2-oxopentanoyl)piperidine-2-carboxamido)-4-(4-methoxyphenyl)butanamido)methyl)-6-azaspiro[2.5]octane-6-carboxylate (**30**) (15.0 mg, 22.4 μmol, 1.0 eq) was dissolved in DCM (0.3 mL) and TFA (60 μL, 780 μmol, 35 eq) was added. The reaction was stirred at room temperature for 2.5 h and then concentrated. The residue was dissolved in DMF (0.3 mL) and 4-chloro-6-(5,6-dihydro-[1,2,4]triazolo[1,5-a]pyrazin-7(8H)-yl)-N-(3-methyl-1-(1H-1,2,4-triazol-3-yl)butyl)-1,3,5-triazin-2-amine (**33**) (12.5 mg, 32.1 μmol, 1.4 eq) and K2CO3 (9.0 mg, 67.3 μmol, 3.0 eq) was added. The mixture was heated to 80 °C and stirred for 24 h. The mixture was diluted with saturated NH4Cl (aq) and extracted with DCM(×3). The combined organic phase was dried over Na2SO4, concentrated, and purified by reverse phase HPLC (35-60% ACN in water with 0.1% formic acid as modifier) to afford 2S)-N-((2R)-1-(((6-(4-(5,6-dihydro-[1,2,4]triazolo[1,5-a]pyrazin-7(8H)-yl)-6-((3-methyl-1-(1H-1,2,4-triazol-3-yl)butyl)amino)-1,3,5-triazin-2-yl)-6-azaspiro[2.5]octan-1-yl)methyl)amino)-4-(4-methoxyphenyl)-1-oxobutan-2-yl)-1-(3,3-dimethyl-2-oxopentanoyl)piperidine-2-carboxamide (**4**) (12.0 mg, 58% yield).

**LCMS:** 922.6 ([M+H]^+^).

### Synthetic scheme for compound 2

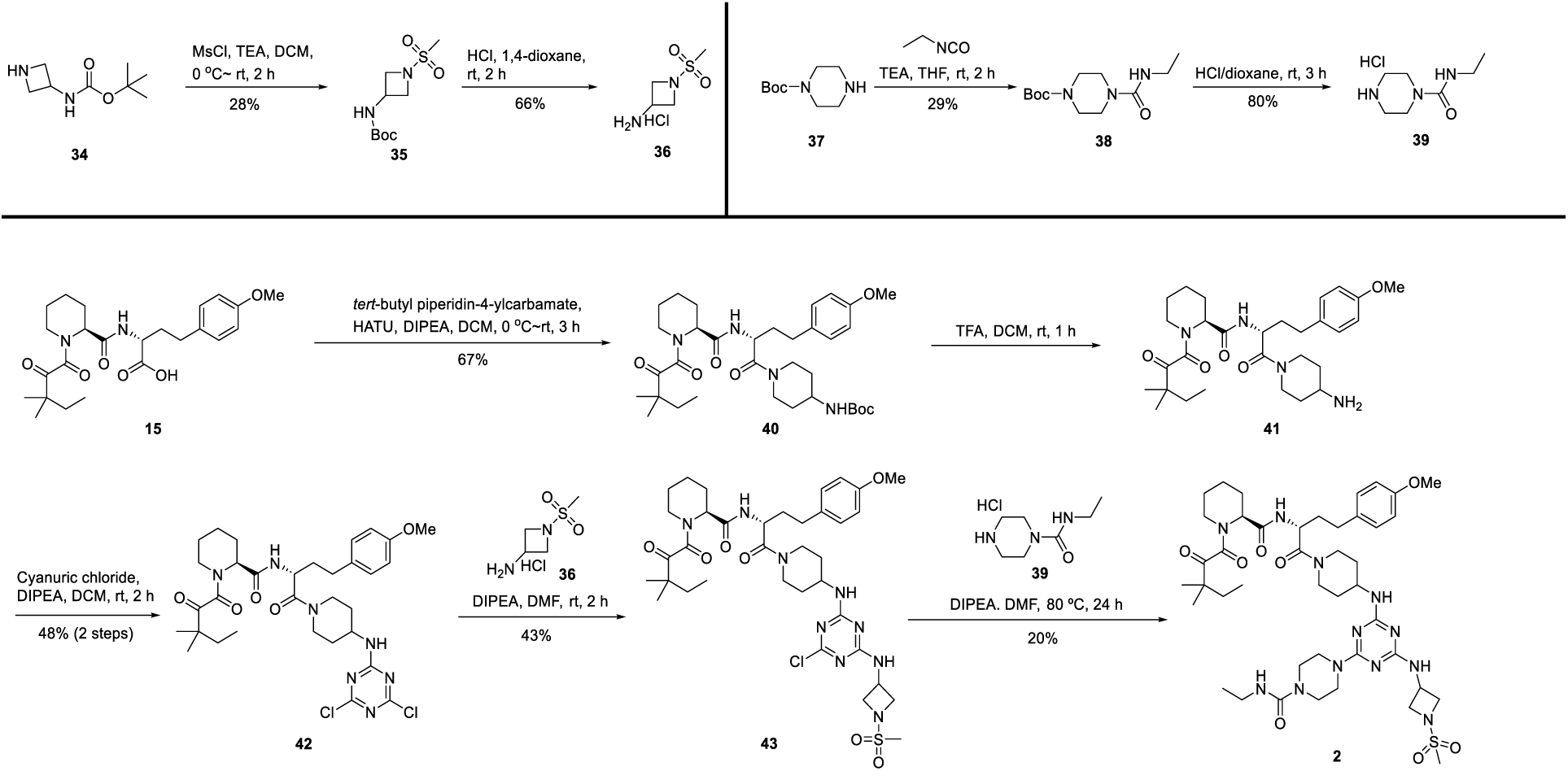

### tert-butyl (1-(methylsulfonyl)azetidin-3-yl)carbamate (35)

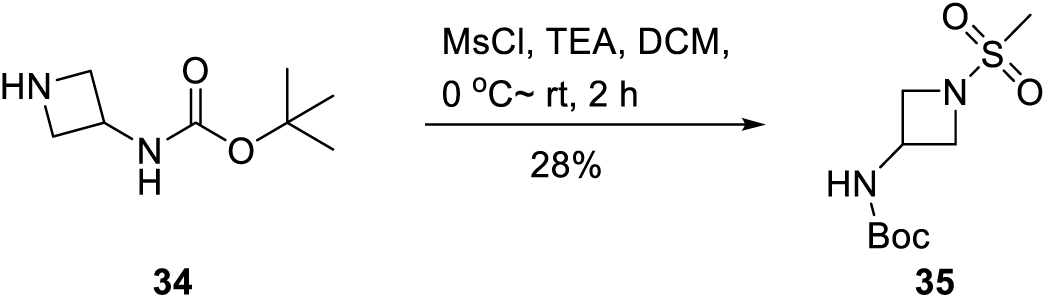

To a solution of tert-butyl azetidin-3-ylcarbamate (**34**) (500 mg, 2.9 mmol, 1.0 eq) and TEA (354 mg, 3.5 mmol, 1.2 eq) in DCM (5 mL) at 0 °C under N2, was added MsCl (401 mg, 3.5 mmol, 1.2 eq) dropwise. The mixture was stirred at room temperature for 2 h, then the reaction quenched with H2O (5 mL). The mixture was extracted with DCM (5 mL x 3), washed with H2O (5 mL x 3) and brine (5 mL x 3), dried over Na2SO4, filtered and concentrated to afford tert-butyl (1-(methylsulfonyl)azetidin-3-yl)carbamate (**35**) (200 mg, 28% yield) as a white solid.

**^1^H NMR (300 MHz, DMSO-*d6*):** *δ* ppm 7.65 (d, *J* = 7.5 Hz, 1H), 4.36-4.31 (m, 1H), 4.03 (t, *J* = 8.2 Hz, 2H), 3.82-3.76 (m, 2H), 2.99 (s, 3H), 1.38 (s, 9H).

### 1-(methylsulfonyl)azetidin-3-amine hydrochloride (36)

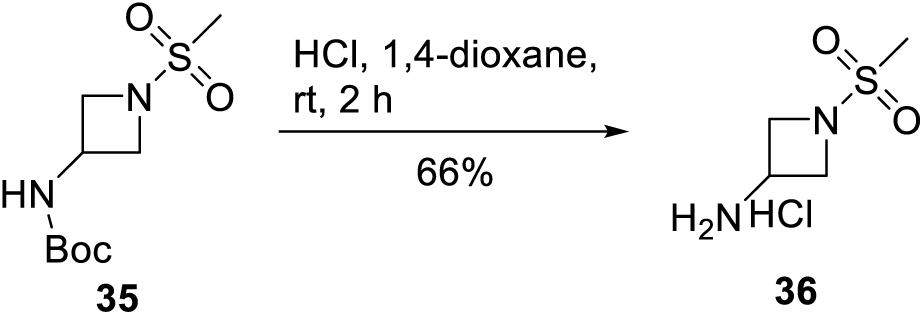

To a solution of tert-butyl (1-(methylsulfonyl)azetidin-3-yl)carbamate (**35**) (200 mg, 0.8 mmol, 1.0 eq) in DCM (1 mL) was added 4M HCl-dioxane (2 mL) at 0 °C. The mixture was stirred at room temperature for 2 h and then concentrated to afford 1-(methylsulfonyl)azetidin-3-amine hydrochloride (**36**) (100 mg, 66% yield) as a white solid.

**^1^H NMR (400 MHz, D2O):** *δ* ppm 4.79-4.62 (m, 1H), 4.56-4.43 (m, 2H), 4.38-4.31 (m, 1H), 4.27-4.19 (m,1H), 4.13-4.02 (m, 1H), 3.13 (d, *J* = 6.7 Hz, 3H).

### tert-butyl 4-(ethylcarbamoyl)piperazine-1-carboxylate (38)

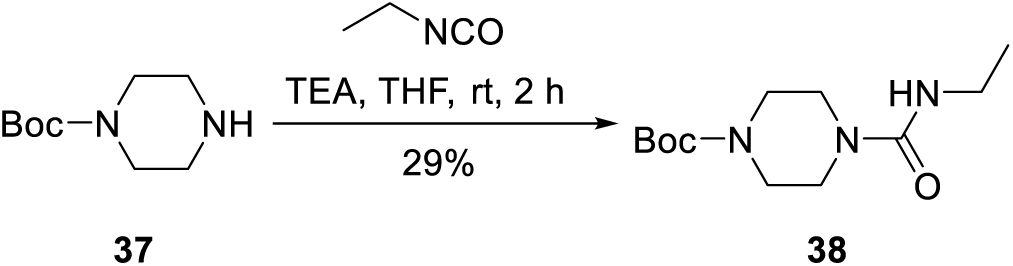

To a solution of tert-butyl piperazine-1-carboxylate (**37**) (5.0 g, 26.8 mmol, 1.0 eq) and TEA (5.4 g, 53.6 mmol, 2.0 eq) in THF (50 mL) at 0 °C under N2 was added ethyl isocyanate (2.3 g, 32.2 mmol, 1.2 eq). The mixture was stirred at room temperature for 2 h and then quenched with H2O (30 mL). The mixture was extracted with DCM (20 mL x 3), washed with brine (20 mL x 3), dried over Na2SO4, filtered and concentrated. The residue was purified with silica gel column (Petroleum ether: EtOAc = 1: 1) to afford tert-butyl 4-(ethylcarbamoyl)piperazine-1-carboxylate (**38**) (2.0 g, 29% yield) as a white solid.

**^1^H NMR (400 MHz, CDCl3):** *δ* ppm 4.39 (s, 1H), 3.48-3.39 (m, 4H), 3.34 (d, *J* = 5.3 Hz, 4H), 3.28 (dd, *J* =7.1, 5.5 Hz, 2H), 1.46 (s, 9H), 1.14 (t, *J* = 7.2 Hz, 3H).

### N-ethylpiperazine-1-carboxamide hydrochloride (39)

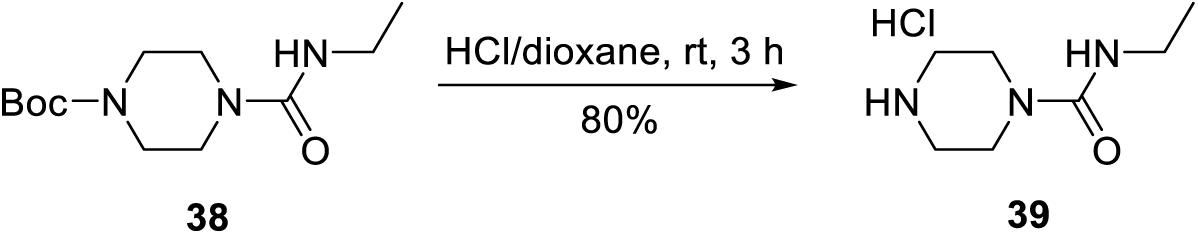

To a solution of tert-butyl 4-(ethylcarbamoyl)piperazine-1-carboxylate (**38**) (2.0 g, 7.8 mmol, 1.0 eq) in DCM (20 mL) at 0 °C was added 4M HCl-dioxane (5 mL). The mixture was stirred at room temperature for 3 h and then concentrated to give N-ethylpiperazine-1-carboxamide hydrochloride (**39**) (1.2 g, 80% yield) as a white solid, which used for next step without any further purification.

**^1^H NMR (400 MHz, DMSO-*d6*):** *δ* ppm 9.43 (s, 2H), 3.64-3.51 (m, 4H), 3.08-3.02 (m, 2H), 3.02-3.00 (m,4H), 1.06 (t, *J* = 7.2 Hz, 3H).

### tert-butyl (1-((R)-2-((S)-1-(3,3-dimethyl-2-oxopentanoyl)piperidine-2-carboxamido)-4-(4-methoxyphenyl)butanoyl)piperidin-4-yl)carbamate (40)

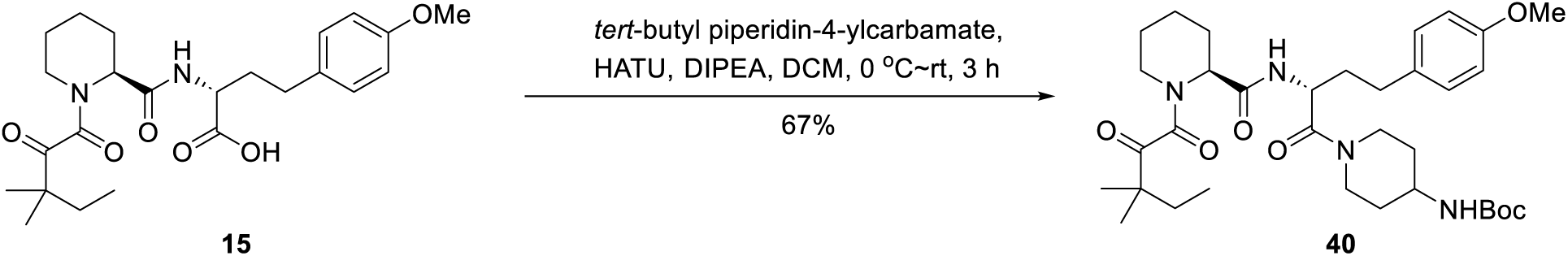

To a solution of (R)-2-((S)-1-(3,3-dimethyl-2-oxopentanoyl)piperidine-2-carboxamido)-4-(4-methoxyphenyl)butanoic acid (**15**) (1.3 g, 2.9 mmol, 1.0 eq) in DCM (20 mL) was added DIPEA (1.1 g, 8.7 mmol, 3.0 eq) and *tert*-butyl piperidin-4-ylcarbamate (697.0 mg, 3.5 mmol, 1.2 eq). The mixture was cooled to 0 °C and HATU (1.3 g, 3.5 mmol, 1.2 eq) was added. The reaction was stirred at room temperature for 3 h, then quenched with H2O (10 mL), extracted with DCM (10 mL x 3), washed with brine (10 mL x 1), dried over Na2SO4, filtered and concentrated. The residue was purified by silica gel chromatography (eluted with Petroleum ether: EtOAc = 3: 1) to afford tert-butyl (1-((R)-2-((S)-1-(3,3-dimethyl-2-oxopentanoyl)piperidine-2-carboxamido)-4-(4-methoxyphenyl)butanoyl)piperidin-4-yl)carbamate (**40**) (1.2 g, 67% yield) as a yellow solid.

**^1^H NMR (300 MHz, CDCl3):** *δ* ppm 7.14-7.08 (m, 2H), 6.84-6.80 (m, 2H), 6.25-6.18 (m, 0.6H), 4.89-4.86 (m, 1H), 4.58-4.33 (m, 2H), 4.06-4.02 (m, 0.4H), 3.79 (s, 3H), 3.68-3.52 (m, 2H), 3.41-3.27 (m, 2H), 3.02-2.88 (m, 1H), 2.85-2.54 (m, 3H), 2.45-2.33 (m, 1H), 2.00-1.82 (m, 4H), 1.79-1.62 (m, 6H), 1.52-1.50 (m, 2H), 1044 (s, 9H), 1.27-1.21 (m, 7H), 0.93-0.88 (m, 3H).

**LCMS:** 629.4 ([M+H]^+^).

### (S)-N-((R)-1-(4-aminopiperidin-1-yl)-4-(4-methoxyphenyl)-1-oxobutan-2-yl)-1-(3,3-dimethyl-2-oxopentanoyl)piperidine-2-carboxamide (41)

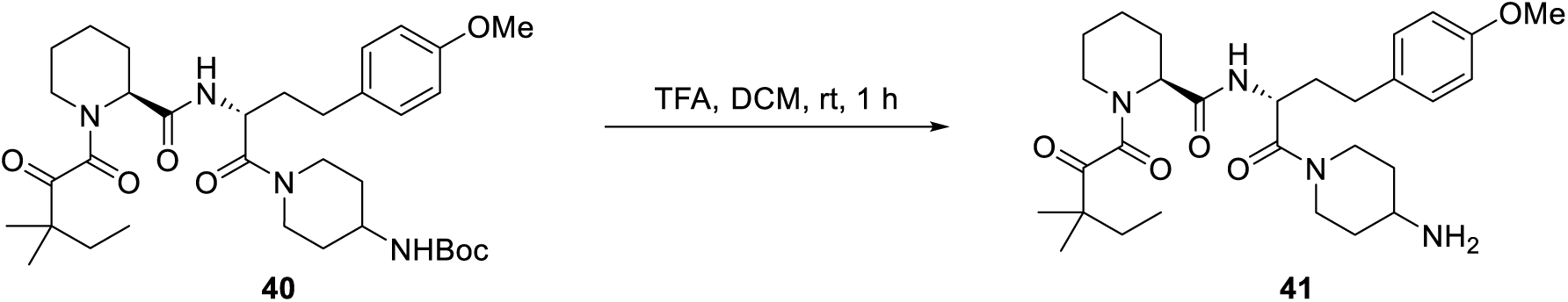

To a solution of tert-butyl (1-((R)-2-((S)-1-(3,3-dimethyl-2-oxopentanoyl)piperidine-2-carboxamido)-4-(4-methoxyphenyl)butanoyl)piperidin-4-yl)carbamate (**40**) (1.1 g, 1.7 mmol, 1.0 eq) in DCM (11 mL) was added TFA (2.5 mL). The mixture was stirred at room temperature for 1 h and then concentrated to afford (S)-N-((R)-1-(4-aminopiperidin-1-yl)-4-(4-methoxyphenyl)-1-oxobutan-2-yl)-1-(3,3-dimethyl-2-oxopentanoyl)piperidine-2-carboxamide (**41**) (1.0 g, crude) as a yellow oil. The material was used in the next step without further purification.

**^1^H NMR (400 MHz, CDCl3):** *δ* ppm 8.43 (s, 1H), 7.53-7.47 (m, 2H), 7.18-7.06 (m, 2H), 6.81 (d, *J* = 7.9 Hz, 2H), 5.15 (s, 1H), 4.84-4.73 (m, 1H), 4.55-4.42 (m, 1H), 4.04 (s, 0.2H), 3.76 (s, 3H), 3.75-3.72 (m, 0.4H), 3.45-3.33 (m, 2H), 3.16 (s, 1H), 2.97-2.91 (m, 1H), 2.72-2.51 (m, 3H), 2.43-2.31 (m, 1H), 2.04-1.78 (m, 4H), 1.72-1.62 (m, 5H), 1.59-1.33 (m, 4H), 1.25-1.20 (m, 6H), 0.90-0.85 (m, 3H).

**LCMS:** 529.3 ([M+H]^+^).

### (S)-N-((R)-1-(4-((4,6-dichloro-1,3,5-triazin-2-yl)amino)piperidin-1-yl)-4-(4-methoxyphenyl)-1-oxobutan-2-yl)-1-(3,3-dimethyl-2-oxopentanoyl)piperidine-2-carboxamide (42)

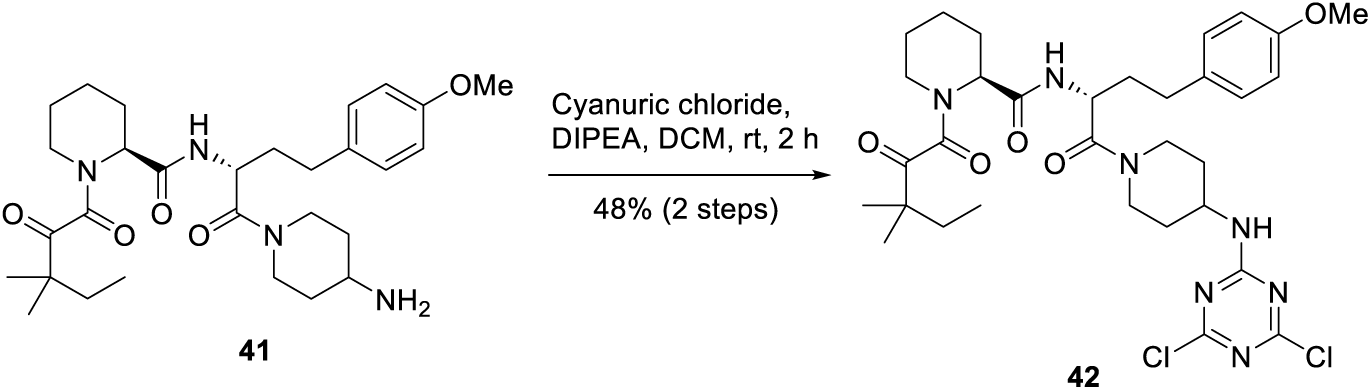

To a solution of (S)-N-((R)-1-(4-aminopiperidin-1-yl)-4-(4-methoxyphenyl)-1-oxobutan-2-yl)-1-(3,3-dimethyl-2-oxopentanoyl)piperidine-2-carboxamide (**41**) (1.0 g, 1.9 mmol, 1.0 eq) in DCM (10 mL) was added DIPEA (1.2 g, 9.5 mmol, 5.0 eq) and cyanuric chloride (420.5 mg, 2.2 mmol, 1.2 eq). The mixture was stirred at room temperature for 2 h and then concentrated to give a residue. The residue was purified by silica gel chromatography (Petroleum ether: EtOAc = 1: 1) to give (S)-N-((R)-1-(4-((4,6-dichloro-1,3,5-triazin-2-yl)amino)piperidin-1-yl)-4-(4-methoxyphenyl)-1-oxobutan-2-yl)-1-(3,3-dimethyl-2-oxopentanoyl)piperidine-2-carboxamide (**42**) (620.0 mg, 48% yield (2 steps)) as a yellow solid.

**LCMS:** 676.2 ([M+H]^+^).

### (S)-N-((R)-1-(4-((4-chloro-6-((1-(methylsulfonyl)azetidin-3-yl)amino)-1,3,5-triazin-2-yl)amino)piperidin-1-yl)-4-(4-methoxyphenyl)-1-oxobutan-2-yl)-1-(3,3-dimethyl-2-oxopentanoyl)piperidine-2-carboxamide (43)

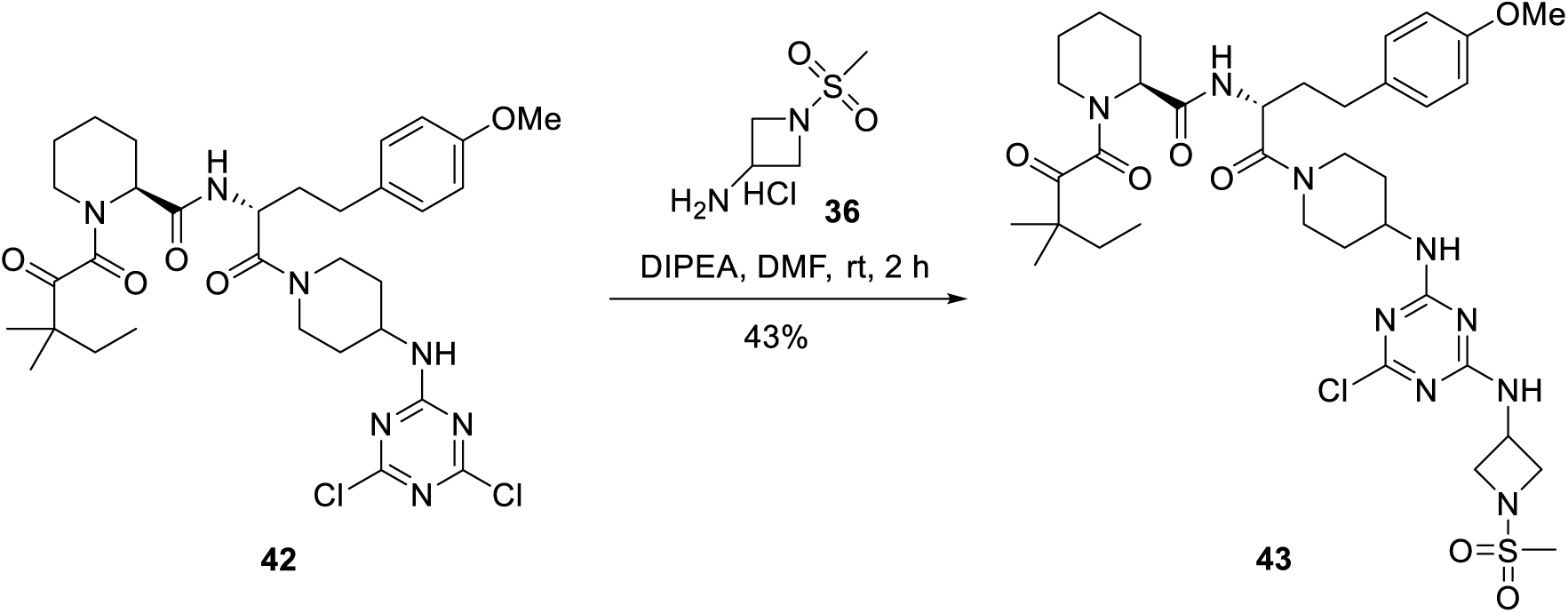

To a solution of (S)-N-((R)-1-(4-((4,6-dichloro-1,3,5-triazin-2-yl)amino)piperidin-1-yl)-4-(4-methoxyphenyl)-1-oxobutan-2-yl)-1-(3,3-dimethyl-2-oxopentanoyl)piperidine-2-carboxamide (**42**) (300 mg, 0.44 mmol, 1.0 eq) in DMF (3 mL) was added DIPEA (284 mg, 2.20 mmol, 5.0 eq) and 1-(methylsulfonyl)azetidin-3-amine hydrochloride (**36**) (98.6 mg, 0.53 mmol, 1.2 eq). The mixture was stirred at room temperature for 2 h, then water (10 mL) was added and the mixture extracted with EtOAc (10 mL x 3). The combined organic phase was washed with brine (10 mL x 3), dried over Na2SO4, filtered and concentrated to afford (S)-N-((R)-1-(4-((4-chloro-6-((1-(methylsulfonyl)azetidin-3-yl)amino)-1,3,5-triazin-2-yl)amino)piperidin-1-yl)-4-(4-methoxyphenyl)-1-oxobutan-2-yl)-1-(3,3-dimethyl-2-oxopentanoyl)piperidine-2-carboxamide (**43**) (150 mg, 43% yield) as a yellow solid.

**LCMS:** 790.2 ([M+H]^+^).

### 4-(4-((1-((R)-2-((S)-1-(3,3-dimethyl-2-oxopentanoyl)piperidine-2-carboxamido)-4-(4-methoxyphenyl)butanoyl)piperidin-4-yl)amino)-6-((1-(methylsulfonyl)azetidin-3-yl)amino)-1,3,5-triazin-2-yl)-N-ethylpiperazine-1-carboxamide (2)

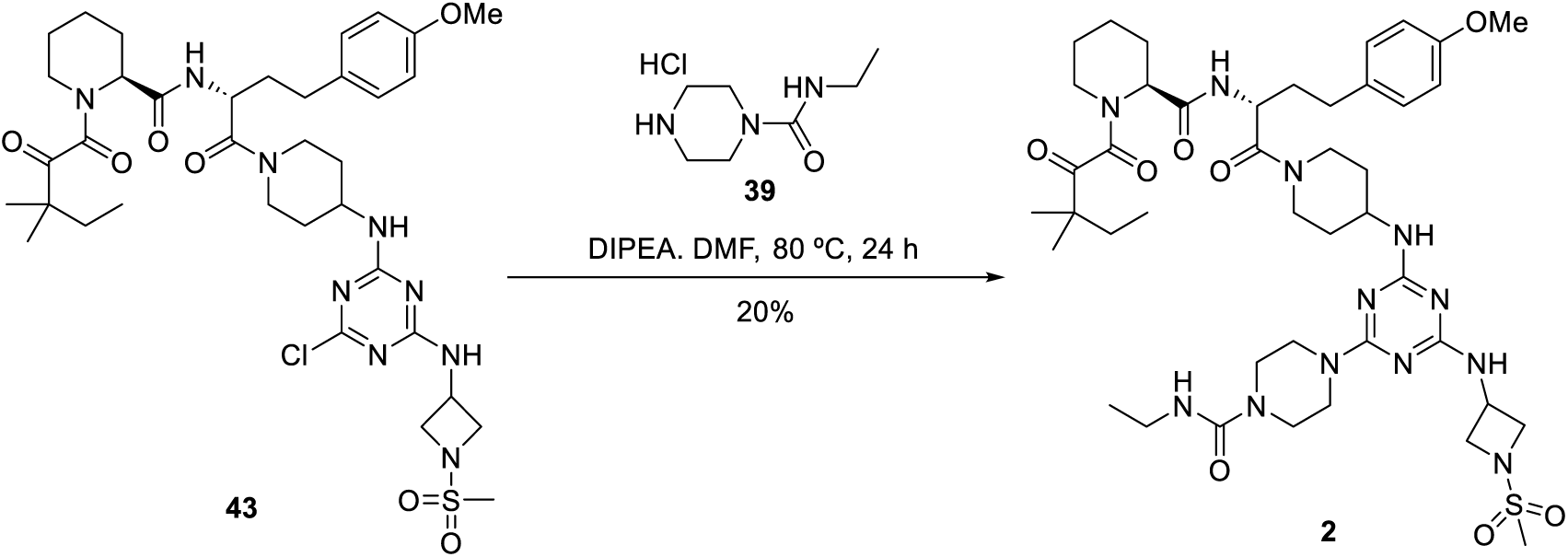

To a solution of (S)-N-((R)-1-(4-((4-chloro-6-((1-(methylsulfonyl)azetidin-3-yl)amino)-1,3,5-triazin-2-yl)amino)piperidin-1-yl)-4-(4-methoxyphenyl)-1-oxobutan-2-yl)-1-(3,3-dimethyl-2-oxopentanoyl)piperidine-2-carboxamide (**43**) (150 mg, 0.19 mmol, 1.0 eq) in DMF (2 mL) was added DIPEA (123 mg, 0.95 mmol, 5.0 eq) and N-ethylpiperazine-1-carboxamide hydrochloride (**39**) (55 mg, 0.29 mmol, 1.5 eq) at room temperature. The mixture was heated to 80 °C and stirred for 24 h. The mixture was diluted with water (10 mL) and extracted with EtOAc (10 mL x 3). The combined organic phase was washed with brine (10 mL x 3), dried over Na2SO4 and concentrated to give a residue. The residue was purified by Prep-TLC (DCM: MeOH = 10: 1) to give 4-(4-((1-((R)-2-((S)-1-(3,3-dimethyl-2-oxopentanoyl)piperidine-2-carboxamido)-4-(4-methoxyphenyl)butanoyl)piperidin-4-yl)amino)-6-((1-(methylsulfonyl)azetidin-3-yl)amino)-1,3,5-triazin-2-yl)-N-ethylpiperazine-1-carboxamide (**2**) (35 mg, 20% yield) as a yellow solid.

**^1^H NMR (400 MHz, DMSO-*d6*):** *δ* ppm 8.14-8.87 (m, 1H), 7.99-7.92 (m, 1H), 7.48-7.22 (m, 0.4H), 7.07 (d,*J* = 8.4 Hz, 2H), 6.84 (d, *J* = 8.5 Hz, 2H), 6.49 (s, 0.6H), 4.99-4.98 (m, 0.6H), 4.58 (s, 1H), 4.29-4.25 (m, 0.4H), 4.25-4.14 (m, 2H), 4.05-4.01 (m, 2H), 3.81 (t, *J* = 7.0 Hz, 2H), 3.71 (s, 3H), 3.60-3.52 (m, 5H), 3.28-3.19 (m, 6H), 3.07-3.04 (m, 2H), 3.01 (s, 3H), 2.18-2.15 (m, 2H), 1.86-1.76 (m, 4H), 1.64-1.53 (m, 8H), 1.36-1.35 (m, 4H), 1.11-1.08 (m, 6H), 1.01 (t, *J* = 7.1 Hz, 3H), 0.85-0.83 (m, 1H), 0.80-0.76 (m, 3H).

**LCMS:** 911.4 ([M+H]^+^).

### Synthetic scheme for compound 1

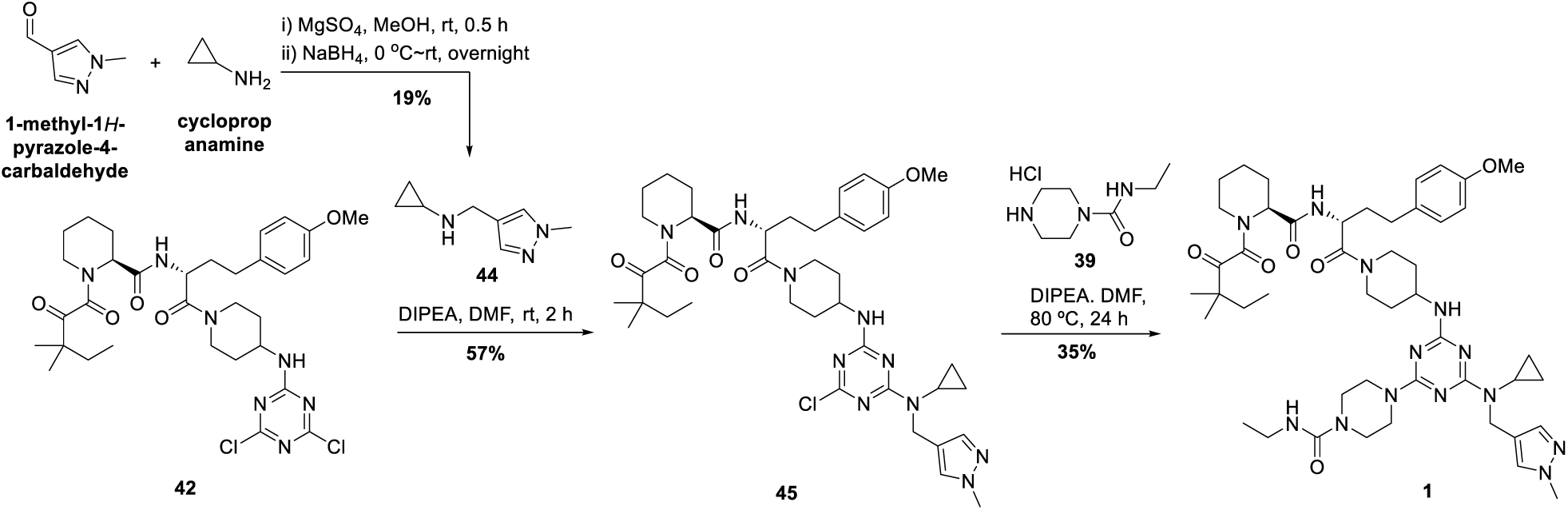

### N-((1-methyl-1H-pyrazol-4-yl)methyl)cyclopropanamine (44)

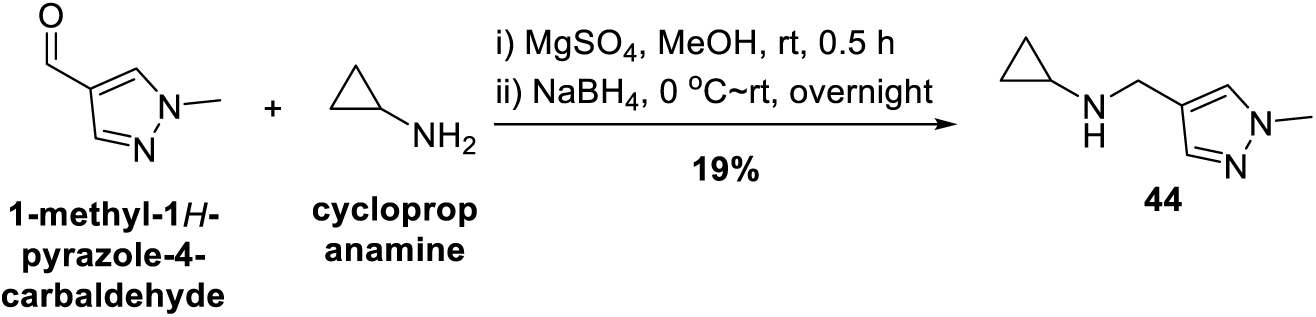

To a solution of 1-methyl-1H-pyrazole-4-carbaldehyde (1.1 g, 10.0 mmol, 1.0 eq) and cyclopropanamine (571 mg, 10.0 mmol, 1.0 eq) in MeOH (10 mL) was added MgSO4 (1.8 g, 15.0 mmol, 1.5 eq) at room temperature. The mixture was stirred at room temperature for 0.5 h, then NaBH4 (757 mg, 20.0 mmol, 2.0 eq) was added at 0 °C and the mixture was stirred at room temperature overnight. Water (10 mL) was added to the mixture and extracted with EtOAc (10 mL x 3). The combined organic phase was washed with brine (10 mL x 3), dried over Na2SO4, filtered and concentrated. The residue was purified by silica gel chromatography (Petroleum ether: EtOAc = 5: 1) to afford N-((1-methyl-1H-pyrazol-4-yl)methyl)cyclopropanamine (**44**) (280 mg, 19%) as a yellow oil.

### (S)-N-((R)-1-(4-((4-chloro-6-(cyclopropyl((1-methyl-1H-pyrazol-4-yl)methyl)amino)-1,3,5-triazin-2-yl)amino)piperidin-1-yl)-4-(4-methoxyphenyl)-1-oxobutan-2-yl)-1-(3,3-dimethyl-2-oxopentanoyl)piperidine-2-carboxamide (45)

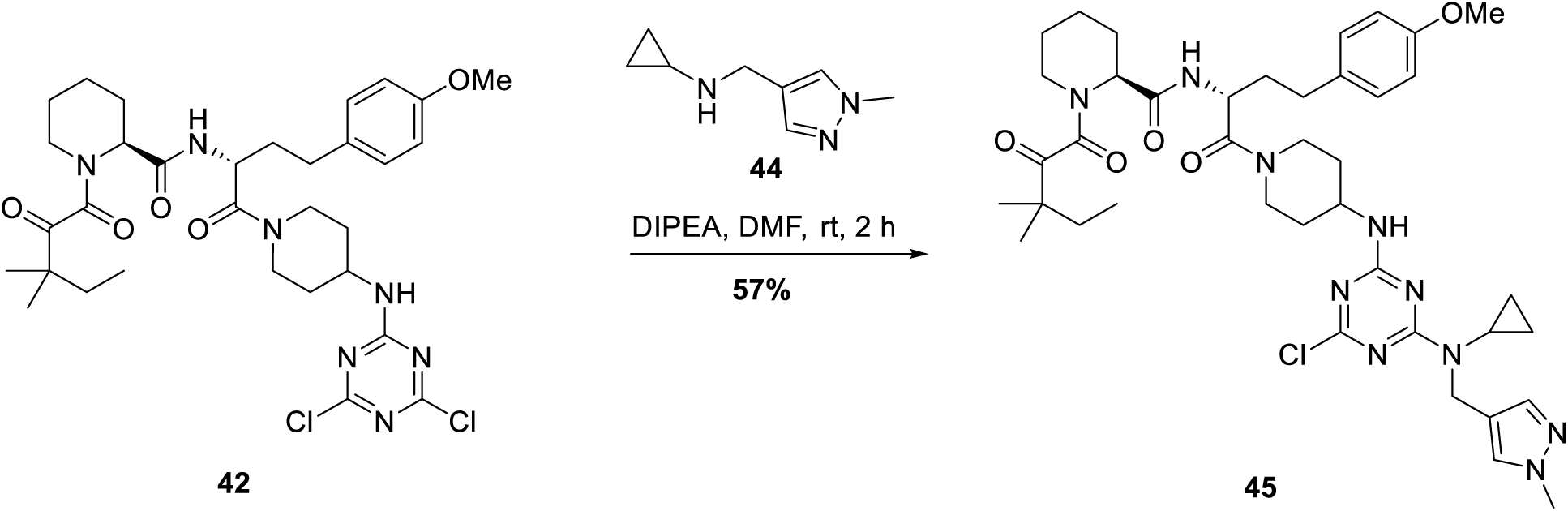

To a solution of (S)-N-((R)-1-(4-((4,6-dichloro-1,3,5-triazin-2-yl)amino)piperidin-1-yl)-4-(4-methoxyphenyl)-1-oxobutan-2-yl)-1-(3,3-dimethyl-2-oxopentanoyl)piperidine-2-carboxamide (**42**) (150 mg, 0.22 mmol, 1.0 eq) in DMF (2 mL) was added DIPEA (142 mg, 1.10 mmol, 5.0 eq) and N-((1-methyl-1H-pyrazol-4-yl)methyl)cyclopropanamine (**44**) (39 mg, 0.26 mmol, 1.2 eq). The mixture was stirred at room temperature for 2 h. After completed, H2O (10 mL) was added to the mixture and extracted with EtOAc (10 mL x 3). The organic phase was washed with brine (10 mL x 3), dried over Na2SO4, filtered and concentrated to give compound **17** (100 mg, 57%) as a yellow solid.

**LCMS:** 791.3 ([M+H]^+^).

### 4-(4-(cyclopropyl((1-methyl-1H-pyrazol-4-yl)methyl)amino)-6-((1-((R)-2-((S)-1-(3,3-dimethyl-2-oxopentanoyl)piperidine-2-carboxamido)-4-(4-methoxyphenyl)butanoyl)piperidin-4-yl)amino)-1,3,5-triazin-2-yl)-N-ethylpiperazine-1-carboxamide (1)

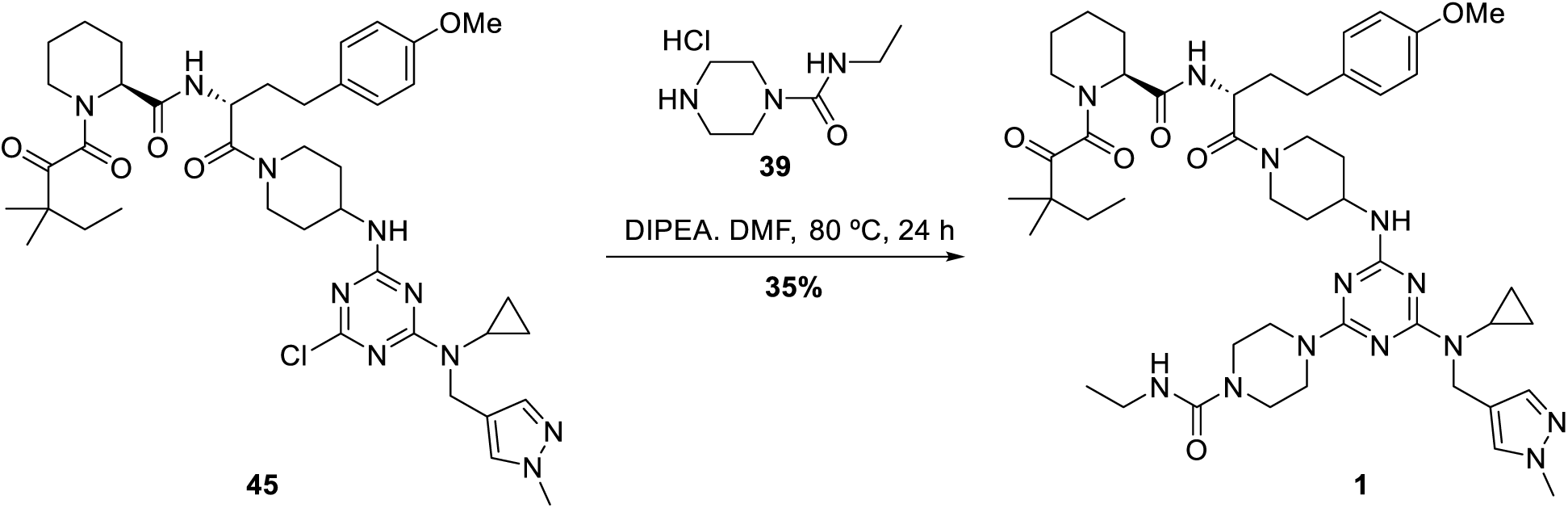

To a solution of (S)-N-((R)-1-(4-((4-chloro-6-(cyclopropyl((1-methyl-1H-pyrazol-4-yl)methyl)amino)-1,3,5-triazin-2-yl)amino)piperidin-1-yl)-4-(4-methoxyphenyl)-1-oxobutan-2-yl)-1-(3,3-dimethyl-2-oxopentanoyl)piperidine-2-carboxamide (**45**) (100 mg, 0.13 mmol, 1.0 eq) in DMF (2 mL) were added DIPEA (84 mg, 0.65 mmol, 5.0 eq) and N-ethylpiperazine-1-carboxamide hydrochloride (**39**) (38 mg, 0.20 mmol, 1.5 eq) at room temperature. The mixture was stirred at 80 °C for 24 h, then diluted with water (10 mL) and the mixture extracted with EtOAc (10 mL x 5). The combined organic phase was washed with brine (10 mL x 3), dried over Na2SO4 and concentrated to give a residue. The residue was purified by Prep-TLC (DCM: MeOH = 10: 1) to afford 4-(4-(cyclopropyl((1-methyl-1H-pyrazol-4-yl)methyl)amino)-6-((1-((R)-2-((S)-1-(3,3-dimethyl-2-oxopentanoyl)piperidine-2-carboxamido)-4-(4-methoxyphenyl)butanoyl)piperidin-4-yl)amino)-1,3,5-triazin-2-yl)-N-ethylpiperazine-1-carboxamide (**1**) (40 mg, 35%) as a white solid.

**^1^H NMR (400 MHz, DMSO-*d6*):** *δ* ppm 8.13-8.07 (m, 1H), 7.97-7.92 (m, 1H), 7.52 (s, 1H), 7.29 (s, 1H), 7.06 (d, *J* = 8.4 Hz, 2H), 6.84 (d, *J* = 8.4 Hz, 2H), 6.80-6.72 (m, 0.5H), 6.48-6.47 (m, 0.5H), 5.00-4.98 (m, 1H), 4.49 (s, 2H), 4.28-4.12 (m, 3H), 4.02-4.00 (m, 1H), 3.76 (s, 3H), 3.70 (s, 3H), 3.64 (s, 3H), 3.55-3.48 (m, 1H), 3.22-3.18 (m, 2H), 3.08-3.02 (m, 2H), 2.44-2.38 (m, 3H), 2.31-.15 (m, 2H), 1.87-1.76 (m, 4H), 1.63-1.50 (m, 8H), 1.36-1.35 (m, 5H), 1.11-1.07 (m, 6H), 1.00 (t, *J* = 7.1 Hz, 3H), 0.93-0.85 (m, 1H), 0.80-0.75 (m, 5H), 0.62 (s, 2H).

**LCMS:** 912.4 ([M+H]^+^).

## ACKNOWLEDGEMENTS

We thank Michael Connolly, Steve M. Canham, Patricia Horton, Shuang Liu, Michael Salcius and Antonin Tutter for their technical assistance around compound synthesis, protein production and cellular assays.

---

Table 1. Crystallographic data and refinement statistics.

**Figure S1.**
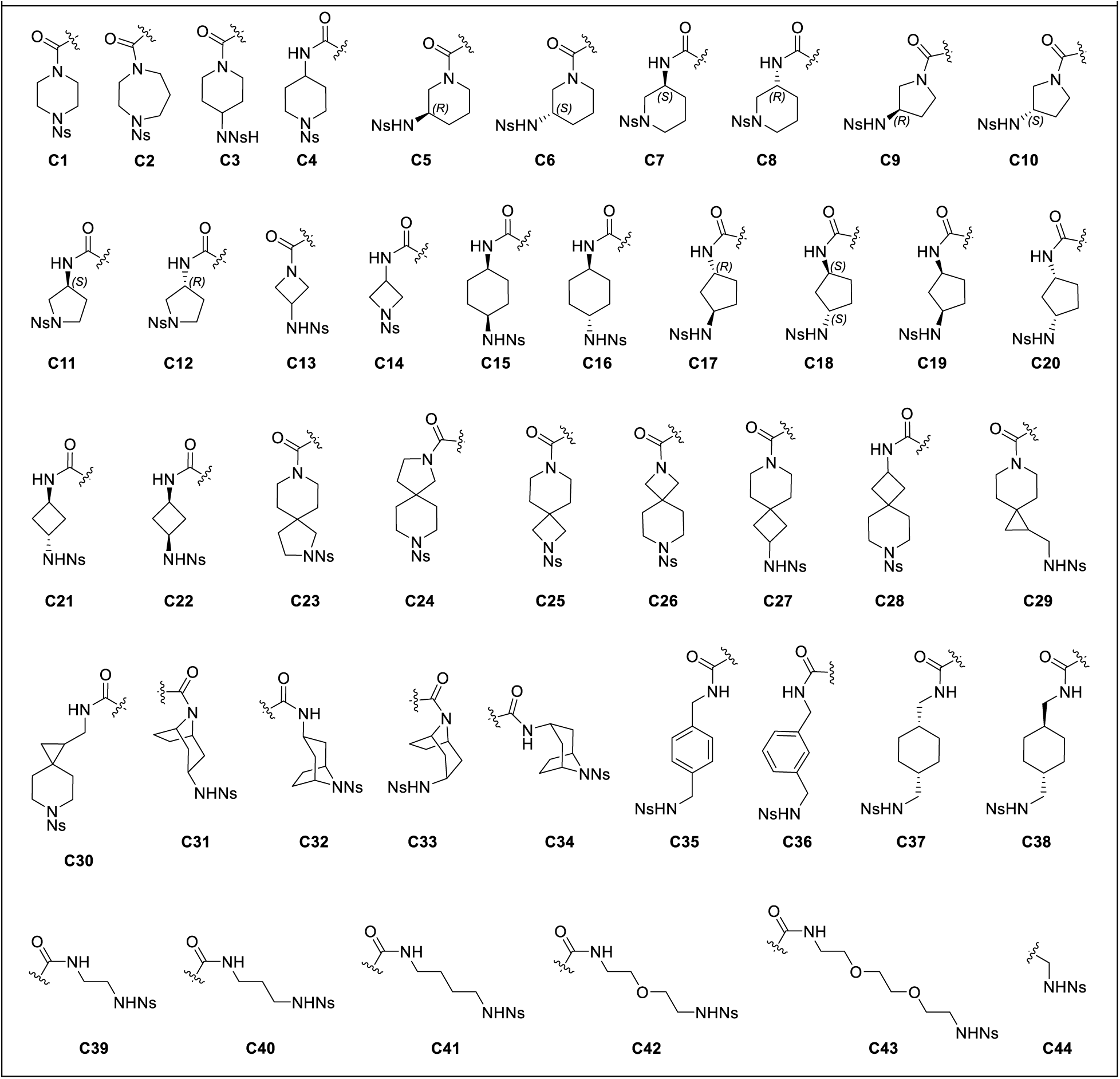
44 Nosyl protected linkers used in the construction of the FKBP-CIP-DEL.

**Figure S2.**
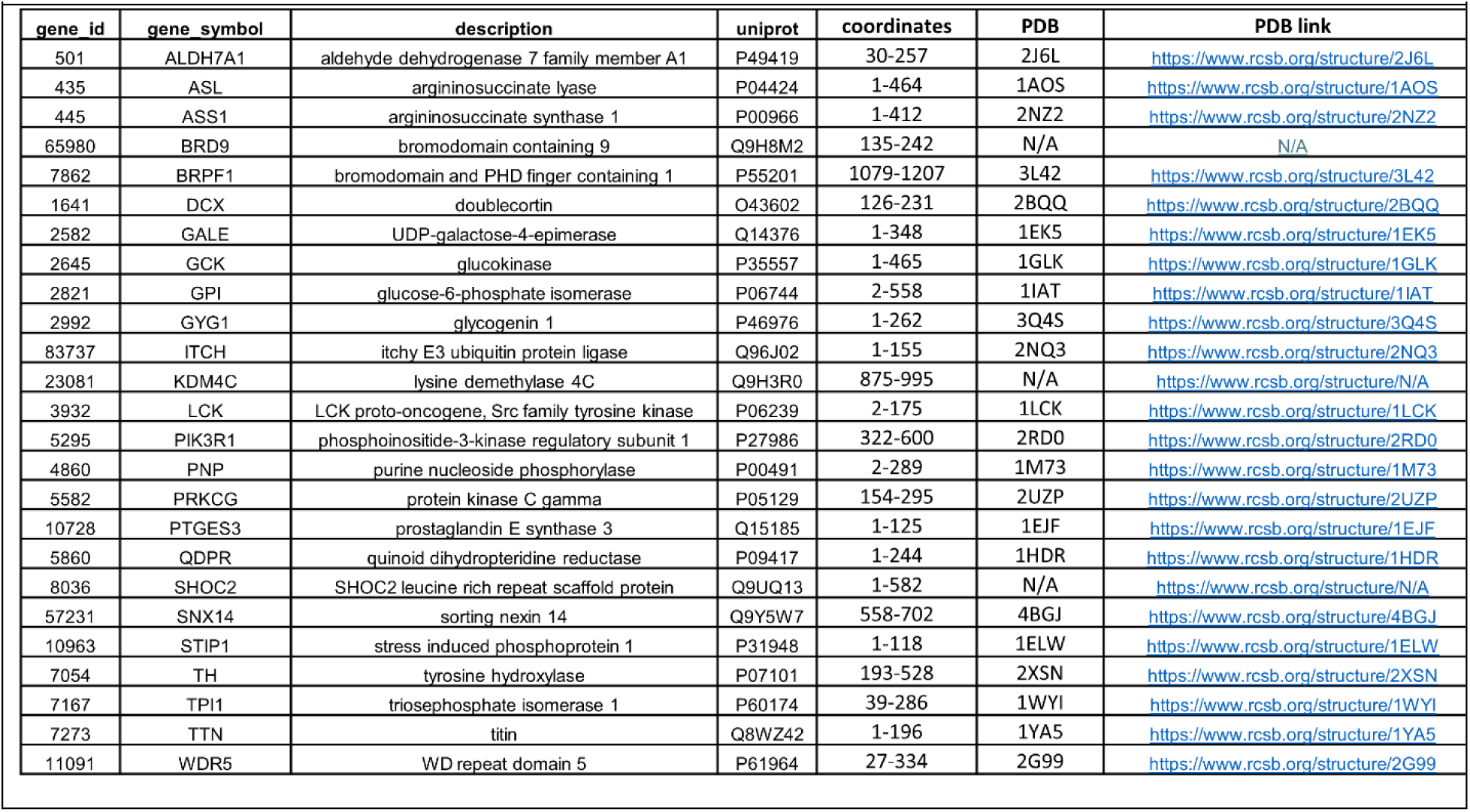
A. Twenty-five proteins were expressed, purified, and screened with the FKBP12-CIP-DEL. For each protein, the amino acid coordinates, and the corresponding PDB reference code along with additional information are listed for each gene symbol.

**Figure S3.**
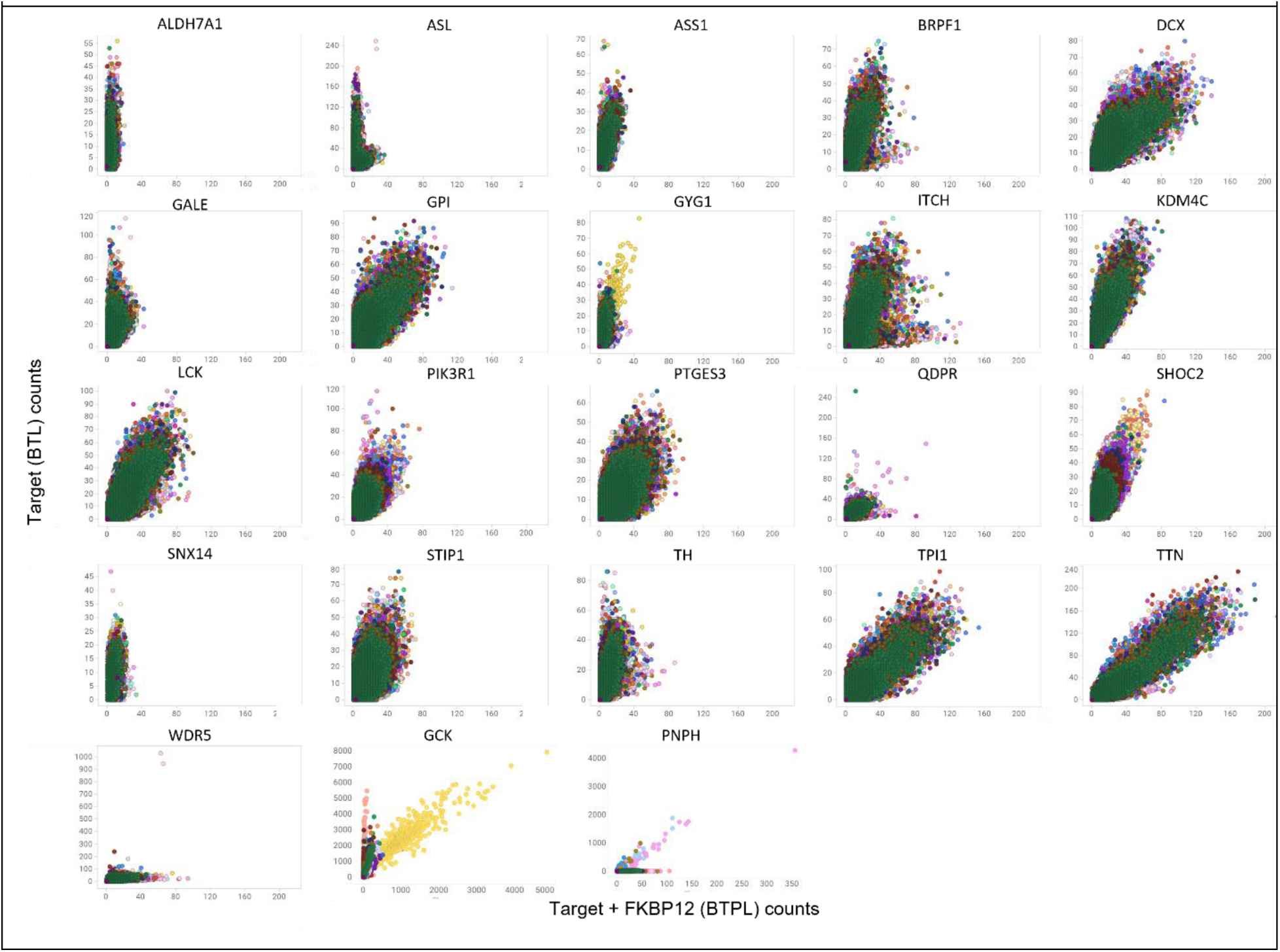
Plots of FKBP-CIP-DEL screening data for 25 targets. Y-axis shows counts against immobilized target alone, BTL, and X-axis shows counts against immobilized target in the presence of 200 µM FKBP12, BTLP.

**Figure S4.**
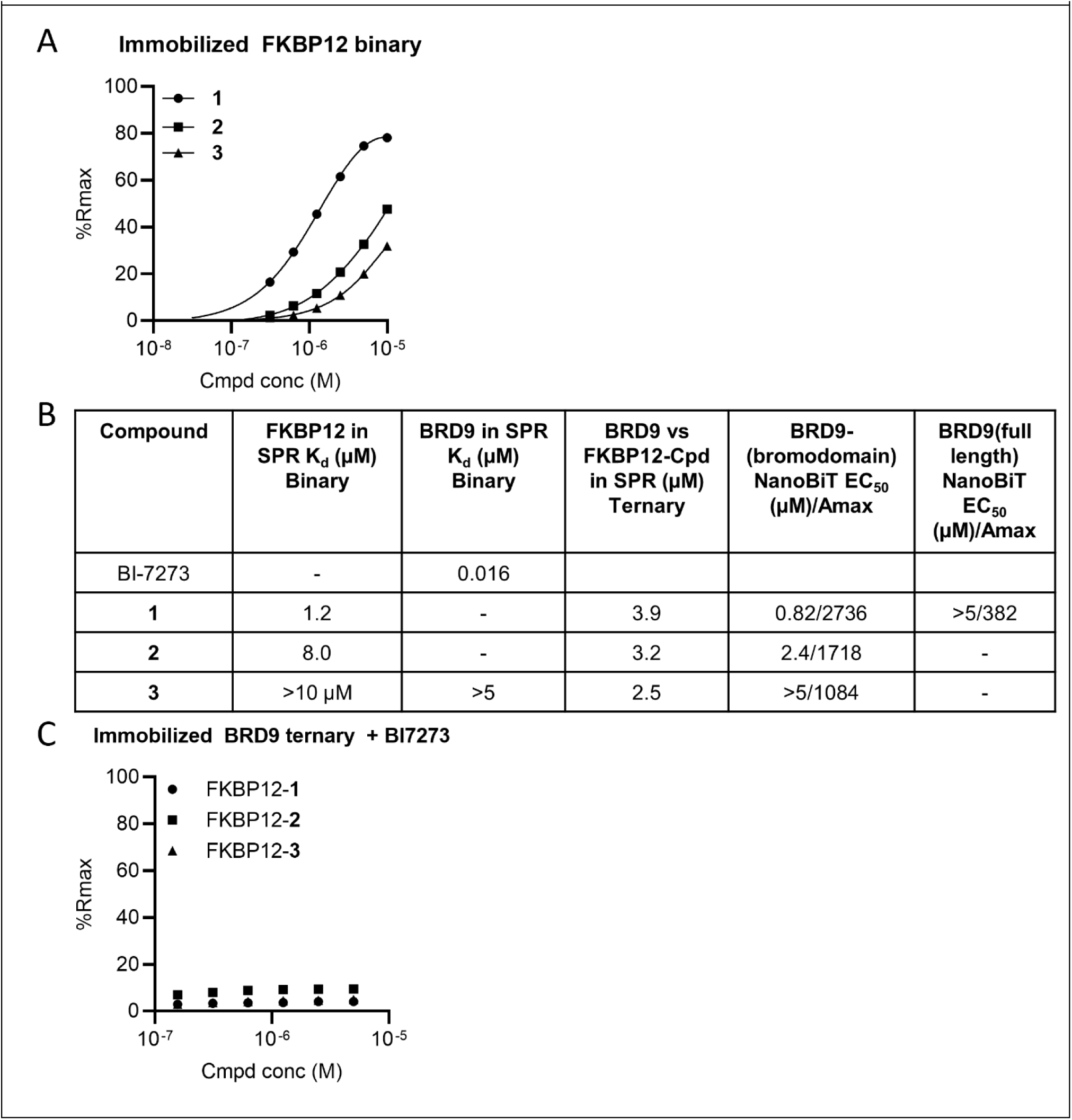
A. SPR graphs for immobilized FKBP12 profiled with titrations of Compounds 1-3. B. The SPR data, binary (compounds only with targets only) and ternary complexes for compounds 1-3. Affinities (KD) are reported in micromolar units; NanoBiT data (EC50s/Amax) are reported for BRD9 (bromodomain) and BRD9(full-length) C. SPR graphs for immobilized BRD9 profiled with titrations of Compounds 1-3 in the presence of BI-7273 and 10 µM.

**Figure S5.**
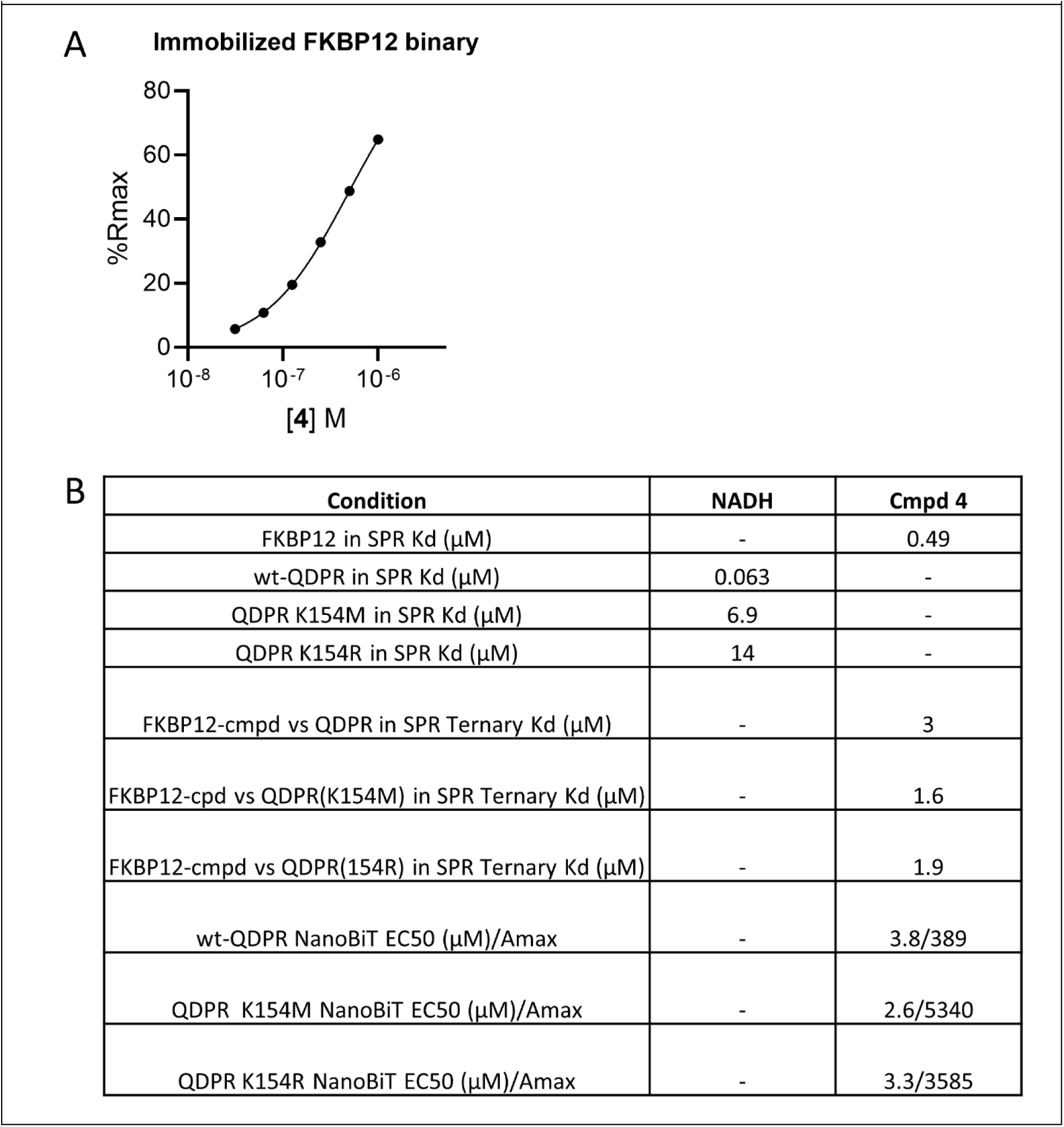
A. SPR graphs for immobilized FKBP12 profiled with titration of compound 4. B. The SPR data, binary (compounds only with targets only) and ternary complexes for Compound 4. Affinities (KD) are reported in micromolar units; NanoBiT data (EC50s/Amax) are reported for QDPR (wildtype and mutants) in micromolar/normalized recruitment.

**Figure S6.**
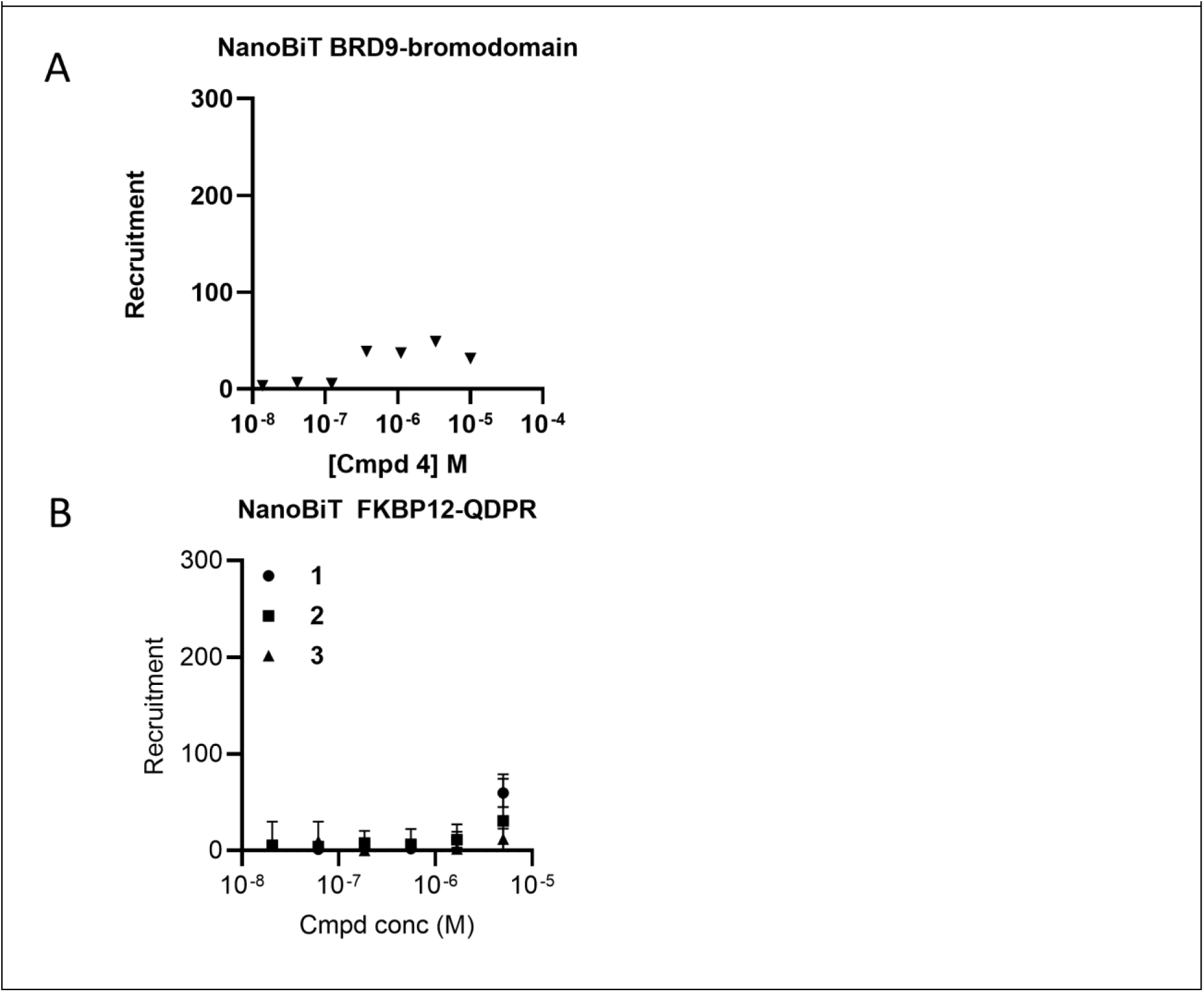
A. NanoBiT data/graphs for the BRD9 bromodomain with Compound 4; B. NanoBiT data/graphs for QDPR with Compound 1.

**Figure S7.**
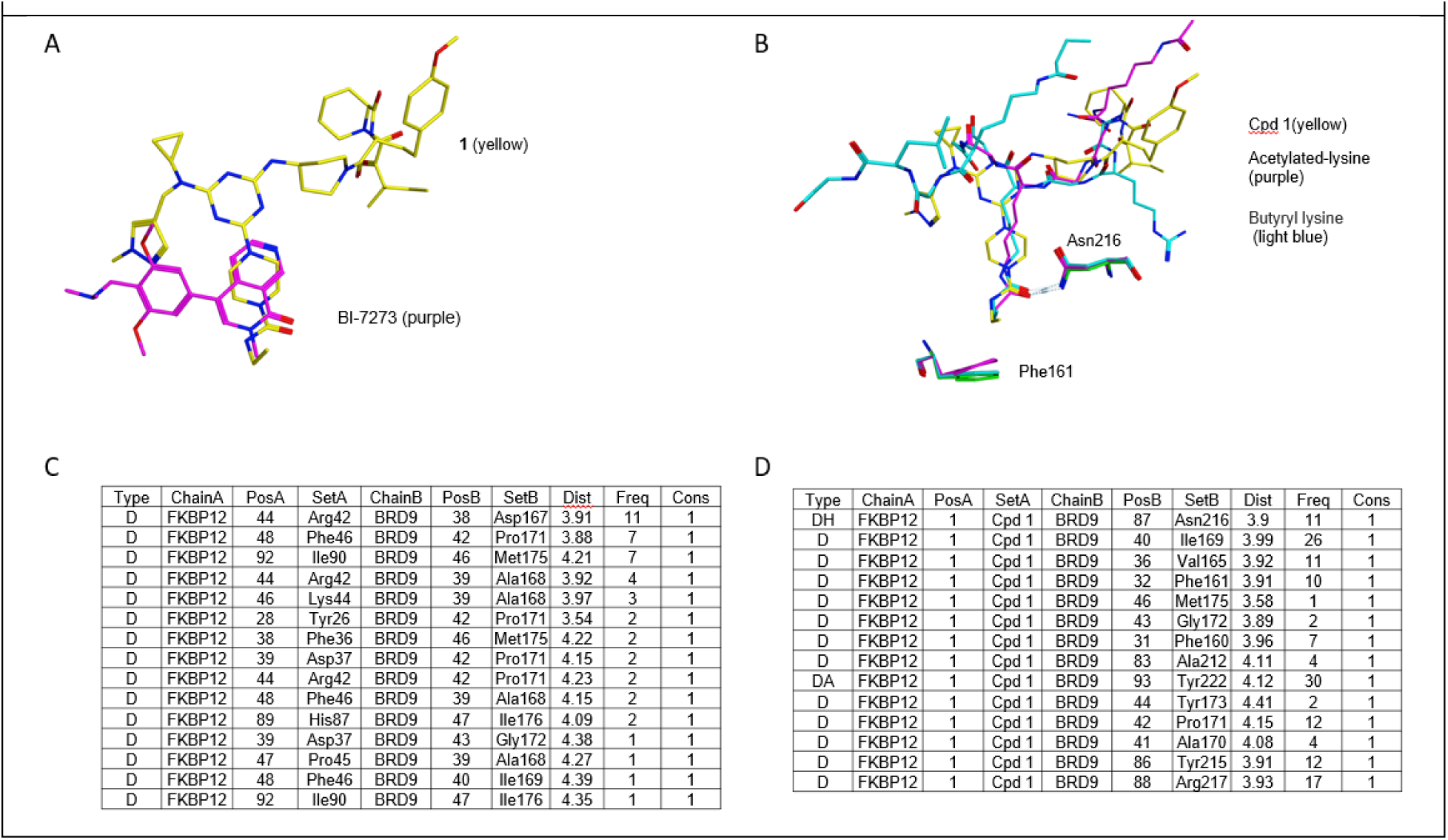
A. An overlay of Compound 1 (yellow) from FKBP12-cpd1-BRD9 ternary complex with BRD9-BI-7273 (purple) binary structure (PDB ID code 5eu1). Only the compounds are shown to illustrate that Compound 1 and BI-7273 occupy similar space in the acetyl-lysine binding pocket; B. An overlay of Compound 1 (yellow) with the x-ray structures for acetyl-lysine (purple, PDB ID code 4yyi) and butyryl-lysine (light blue, PDB ID code 4yy6), with Asn216 H-bonding to each molecule. Phe161 is also shown in green (9du1), purple (4yyi) and light blue (4yy6). C. List of all protein-protein interactions, within 4.5 Å, for FKBP12-Compound 1-BRD9 ternary complex, export from MOE. Distances are aggregated per residue. D. List of Compound 1-BRD9 interactions, within 4.5 Å, for FKBP12-Compound 1-BRD9 ternary complex, exported from MOE. Distances are aggregated per residue.

**Figure S8.**
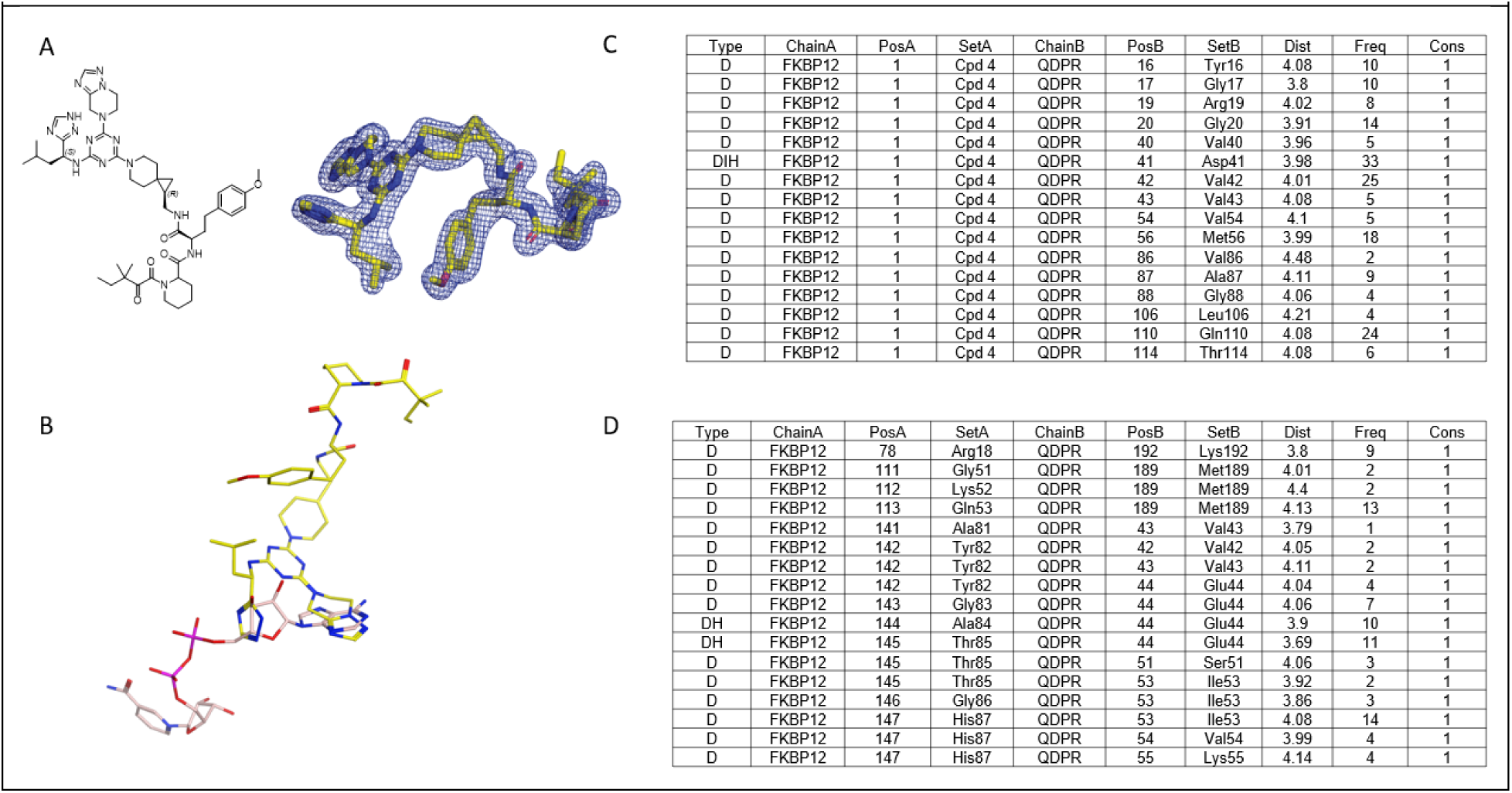
A. Compound 4 with assigned stereochemistry based on the high-resolution ternary structure data. 2fo-fc electron density map within 1.9 Å of the ligand contoured at 1σ level. B. An overlay of Compound 4 (yellow) from the FKBP12-cpd4-QDPR ternary complex with QDPR-NADH (salmon) binary structure (PDB ID code 4hdr). Only the compounds are shown to illustrate that Compound 4 and NADH occupy an overlapping space in the NADH-binding pocket; C. List of Compound 4-QDPR interactions, within 4.5 Å, for the FKBP12-Compound 4-QDPR ternary complex, exported from MOE. Distances are aggregated per residue. D. List of all protein-protein interactions, within 4.5 Å, for the FKBP12-cpd4-QDPR ternary complex, exported from MOE. Distances are aggregated per residue.

## REFERENCES

(1) Schreiber, S. L. The Rise of Molecular Glues. Cell 2021, 184 (1), 3–9. 10.1016/j.cell.2020.12.020.

(2) Bonazzi, S.; d’Hennezel, E.; Beckwith, R. E. J.; Xu, L.; Fazal, A.; Magracheva, A.; Ramesh, R, Cernijenko, A.; Antonakos, B.; Bhang, H. C.; Caro, R. G.; Cobb, J. S.; Ornelas, E.; Ma, X.;Wartchow, C. A.; Clifton, M. C.; Forseth, R. R.; Fortnam, B. H.; Lu, H.; Csibi, A.; Tullai, J.;Carbonneau, S.; Thomsen, N. M.; Larrow, J.; Chie-Leon, B.; Hainzl, D.; Gu, Y.; Lu, D.; Meyer, M. J.; Alexander, D.; Kinyamu-Akunda, J.; Sabatos-Peyton, C. A.; Dales, N. A.; Zecri, F. J.; Jain, R. K.; Shulok, J.; Wang, Y. K.; Briner, K.; Porter, J. A.; Tallarico, J. A.; Engelman, J. A.; Dranoff, G.; Bradner, J. E.; Visser, M.; Solomon, J. M. Discovery and Characterization of a Selective IKZF2 Glue Degrader for Cancer Immunotherapy. Cell Chem Biol 2023, 30 (3), 235–247 e12. 10.1016/j.chembiol.2023.02.005.

(3) Henning, N. J.; Boike, L.; Spradlin, J. N.; Ward, C. C.; Liu, G.; Zhang, E.; Belcher, B. P.; Brittain, S. M.; Hesse, M. J.; Dovala, D.; McGregor, L. M.; Valdez Misiolek, R.; Plasschaert, L. W.; Rowlands, D. J.; Wang, F.; Frank, A. O.; Fuller, D.; Estes, A. R.; Randal, K. L.; Panidapu, A.; McKenna, J. M.; Tallarico, J. A.; Schirle, M.; Nomura, D. K. Deubiquitinase-Targeting Chimeras for Targeted Protein Stabilization. Nat Chem Biol 2022, 18 (4), 412–421. 10.1038/s41589-022-00971-2.

(4) Siriwardena, S. U.; Munkanatta Godage, D. N. P.; Shoba, V. M.; Lai, S.; Shi, M.; Wu, P.; Chaudhary, S. K.; Schreiber, S. L.; Choudhary, A. Phosphorylation-Inducing Chimeric Small Molecules. J Am Chem Soc 2020, 142 (33), 14052–14057. 10.1021/jacs.0c05537.

(5) Chen, P.-H.; Hu, Z.; An, E.; Okeke, I.; Zheng, S.; Luo, X.; Gong, A.; Jaime-Figueroa, S.; Crews, C. M. Modulation of Phosphoprotein Activity by Phosphorylation Targeting Chimeras (PhosTACs). ACS Chem Biol 2021, 16 (12), 2808–2815. 10.1021/acschembio.1c00693.

(6) Gourisankar, S.; Krokhotin, A.; Ji, W.; Liu, X.; Chang, C.-Y.; Kim, S. H.; Li, Z.; Wenderski, W.;Simanauskaite, J. M.; Yang, H.; Vogel, H.; Zhang, T.; Green, M. R.; Gray, N. S.; Crabtree, G. R. Rewiring Cancer Drivers to Activate Apoptosis. Nature 2023, 620 (7973), 417–425. 10.1038/s41586-023-06348-2.

(7) Schreiber, S. L. Molecular Glues and Bifunctional Compounds: Therapeutic Modalities Based on Induced Proximity. Cell Chem Biol 2024, 31 (6), 1050–1063. 10.1016/j.chembiol.2024.05.004.

(8) Bussiere, D. E.; Xie, L.; Srinivas, H.; Shu, W.; Burke, A.; Be, C.; Zhao, J.; Godbole, A.; King, D.;Karki, R. G.; Hornak, V.; Xu, F.; Cobb, J.; Carte, N.; Frank, A. O.; Frommlet, A.; Graff, P.; Knapp, M.; Fazal, A.; Okram, B.; Jiang, S.; Michellys, P. Y.; Beckwith, R.; Voshol, H.; Wiesmann, C.; Solomon, J. M.; Paulk, J. Structural Basis of Indisulam-Mediated RBM39 Recruitment to DCAF15 E3 Ligase Complex. Nat Chem Biol 2020, 16 (1), 15–23. 10.1038/s41589-019-0411-6.

(9) Lu, G.; Middleton, R. E.; Sun, H.; Naniong, M.; Ott, C. J.; Mitsiades, C. S.; Wong, K. K.; Bradner, J. E.; Kaelin Jr., W. G. The Myeloma Drug Lenalidomide Promotes the Cereblon-Dependent Destruction of Ikaros Proteins. Science (1979) 2014, 343 (6168), 305–309. 10.1126/science.1244917.

(10) Chamberlain, P. P.; Lopez-Girona, A.; Miller, K.; Carmel, G.; Pagarigan, B.; Chie-Leon, B.; Rychak, E.; Corral, L. G.; Ren, Y. J.; Wang, M.; Riley, M.; Delker, S. L.; Ito, T.; Ando, H.; Mori, T.; Hirano, Y.; Handa, H.; Hakoshima, T.; Daniel, T. O.; Cathers, B. E. Structure of the Human Cereblon-DDB1-Lenalidomide Complex Reveals Basis for Responsiveness to Thalidomide Analogs. Nat Struct Mol Biol 2014, 21 (9), 803–809. 10.1038/nsmb.2874.

(11) Matyskiela, M. E.; Lu, G.; Ito, T.; Pagarigan, B.; Lu, C. C.; Miller, K.; Fang, W.; Wang, N. Y.;Nguyen, D.; Houston, J.; Carmel, G.; Tran, T.; Riley, M.; Nosaka, L.; Lander, G. C.; Gaidarova, S.; Xu, S.; Ruchelman, A. L.; Handa, H.; Carmichael, J.; Daniel, T. O.; Cathers, B. E.; Lopez-Girona, A.; Chamberlain, P. P. A Novel Cereblon Modulator Recruits GSPT1 to the CRL4(CRBN) Ubiquitin Ligase. Nature 2016, 535 (7611), 252–257. 10.1038/nature18611.

(12) Shigdel, U. K.; Lee, S. J.; Sowa, M. E.; Bowman, B. R.; Robison, K.; Zhou, M.; Pua, K. H.; Stiles, D. T.; Blodgett, J. A. V; Udwary, D. W.; Rajczewski, A. T.; Mann, A. S.; Mostafavi, S.; Hardy, T.;Arya, S.; Weng, Z.; Stewart, M.; Kenyon, K.; Morgenstern, J. P.; Pan, E.; Gray, D. C.; Pollock, R. M.; Fry, A. M.; Klausner, R. D.; Townson, S. A.; Verdine, G. L. Genomic Discovery of an Evolutionarily Programmed Modality for Small-Molecule Targeting of an Intractable Protein Surface. Proc Natl Acad Sci U S A 2020, 117 (29), 17195–17203. 10.1073/pnas.2006560117.

(13) Petzold, G.; Fischer, E. S.; Thoma, N. H. Structural Basis of Lenalidomide-Induced CK1alpha Degradation by the CRL4(CRBN) Ubiquitin Ligase. Nature 2016, 532 (7597), 127–130. 10.1038/nature16979.

(14) Liu, S.; Tong, B.; Mason, J. W.; Ostrem, J. M.; Tutter, A.; Hua, B. K.; Tang, S. A.; Bonazzi, S.; Briner, K.; Berst, F.; Zécri, F. J.; Schreiber, S. L. Rational Screening for Cooperativity in Small-Molecule Inducers of Protein–Protein Associations. J Am Chem Soc 2023, 145 (42), 23281–23291. 10.1021/jacs.3c08307.

(15) Jeremy W. Mason Matthias V. Westphal Antonin Tutter Gregory Michaud Wei Shu Xiaolei Ma Connor W. Coley Paul A. Clemons Simone Bonazzi Frédéric Berst Frédéric J. Zécri Karin Briner View ORCID ProfileStuart L. Schreiber, L. H. DNA-Encoded Library (DEL)-Enabled Discovery of Proximity-Inducing Small Molecules. bioRxiv 2022.

(16) Clackson, T.; Yang, W.; Rozamus, L. W.; Hatada, M.; Amara, J. F.; Rollins, C. T.; Stevenson, L. F.; Magari, S. R.; Wood, S. A.; Courage, N. L.; Lu, X.; Cerasoli, F.; Gilman, M.; Holt, D. A. Redesigning an FKBP–Ligand Interface to Generate Chemical Dimerizers with Novel Specificity. Proceedings of the National Academy of Sciences 1998, 95 (18), 10437–10442. 10.1073/pnas.95.18.10437.

(17) Filippakopoulos, P.; Picaud, S.; Mangos, M.; Keates, T.; Lambert, J.-P.; Barsyte-Lovejoy, D.; Felletar, I.; Volkmer, R.; Müller, S.; Pawson, T.; Gingras, A.-C.; Arrowsmith, C. H.; Knapp, S. Histone Recognition and Large-Scale Structural Analysis of the Human Bromodomain Family. Cell. 2012, 149 (1), 214–231. 10.1016/j.cell.2012.02.013.

(18) Flynn, E. M.; Huang, O. W.; Poy, F.; Oppikofer, M.; Bellon, S. F.; Tang, Y.; Cochran, A. G. A Subset of Human Bromodomains Recognizes Butyryllysine and Crotonyllysine Histone Peptide Modifications. Structure 2015, 23 (10), 1801–1814. 10.1016/j.str.2015.08.004.

(19) Martin, L. J.; Koegl, M.; Bader, G.; Cockcroft, X.-L.; Fedorov, O.; Fiegen, D.; Gerstberger, T.; Hofmann, M. H.; Hohmann, A. F.; Kessler, D.; Knapp, S.; Knesl, P.; Kornigg, S.; Müller, S.; Nar, H.; Rogers, C.; Rumpel, K.; Schaaf, O.; Steurer, S.; Tallant, C.; Vakoc, C. R.; Zeeb, M.; Zoephel, A.; Pearson, M.; Boehmelt, G.; McConnell, D. Structure-Based Design of an in Vivo Active Selective BRD9 Inhibitor. J Med Chem 2016, 59 (10), 4462–4475. 10.1021/acs.jmedchem.5b01865.

(20) Lockyer, J.; Cook, R. G.; Milstien, S.; Kaufman, S.; Woo, S. L.; Ledley, F. D. Structure and Expression of Human Dihydropteridine Reductase. Proceedings of the National Academy of Sciences 1987, 84 (10), 3329–3333. 10.1073/pnas.84.10.3329.

(21) Kiefer, P. M.; Varughese, K. I.; Su, Y.; Xuong, N.-H.; Chang, C.-F.; Gupta, P.; Bray, T.; Whiteley, J. M. Altered Structural and Mechanistic Properties of Mutant Dihydropteridine Reductases. Journal of Biological Chemistry 1996, 271 (7), 3437–3444. 10.1074/jbc.271.7.3437.

(22) Futer, O.; DeCenzo, M. T.; Aldape, R. A.; Livingston, D. J. FK506 Binding Protein Mutational Analysis. Journal of Biological Chemistry 1995, 270 (32), 18935–18940. 10.1074/jbc.270.32.18935.

(23) Choi, J.; Chen, J.; Schreiber, S. L.; Clardy, J. Structure of the FKBP12-Rapamycin Complex Interacting with Binding Domain of Human FRAP. Science (1979) 1996, 273 (5272), 239–242. 10.1126/science.273.5272.239.

(24) Ichikawa, S.; Flaxman, H. A.; Xu, W.; Vallavoju, N.; Lloyd, H. C.; Wang, B.; Shen, D.; Pratt, M. R.; Woo, C. M. The E3 Ligase Adapter Cereblon Targets the C-Terminal Cyclic Imide Degron. Nature 2022, 610 (7933), 775–782. 10.1038/s41586-022-05333-5.

(25) Varughese, K. I.; Skinner, M. M.; Whiteley, J. M.; Matthews, D. A.; Xuong, N. H. Crystal Structure of Rat Liver Dihydropteridine Reductase. Proceedings of the National Academy of Sciences 1992, 89 (13), 6080–6084. 10.1073/pnas.89.13.6080.

(26) Su, Y.; Varughese, K. I.; Xuong, N. H.; Bray, T. L.; Roche, D. J.; Whiteley, J. M. The Crystallographic Structure of a Human Dihydropteridine Reductase NADH Binary Complex Expressed in Escherichia Coli by a CDNA Constructed from Its Rat Homologue. J Biol Chem 1993, 268 (36), 26836–26841. 10.2210/pdb1hdr/pdb.

(27) Mo, X.; Niu, Q.; Ivanov, A. A.; Tsang, Y. H.; Tang, C.; Shu, C.; Li, Q.; Qian, K.; Wahafu, A.; Doyle, S. P.; Cicka, D.; Yang, X.; Fan, D.; Reyna, M. A.; Cooper, L. A. D.; Moreno, C. S.; Zhou, W.;Owonikoko, T. K.; Lonial, S.; Khuri, F. R.; Du, Y.; Ramalingam, S. S.; Mills, G. B.; Fu, H. Systematic Discovery of Mutation-Directed Neo-Protein-Protein Interactions in Cancer. Cell 2022, 185 (11), 1974–1985.e12. 10.1016/j.cell.2022.04.014.

(28) Kabsch, W. XDS. Acta Crystallogr D Biol Crystallogr 2010, 66 (2), 125–132. 10.1107/S0907444909047337.

(29) McCoy, A. J.; Grosse-Kunstleve, R. W.; Adams, P. D.; Winn, M. D.; Storoni, L. C.; Read, R. J. Phaser Crystallographic Software. J Appl Crystallogr 2007, 40 (Pt 4), 658–674. 10.1107/S0021889807021206.

(30) Martinelli, P.; Schaaf, O.; Mantoulidis, A.; Martin, L. J.; Fuchs, J. E.; Bader, G.; Gollner, A.; Wolkerstorfer, B.; Rogers, C.; Balıkçı, E.; Lipp, J. J.; Mischerikow, N.; Doebel, S.; Gerstberger, T.; Sommergruber, W.; Huber, K. V. M.; Böttcher, J. Discovery of a Chemical Probe to Study Implications of BPTF Bromodomain Inhibition in Cellular and *in Vivo* Experiments. ChemMedChem 2023, 18 (6). 10.1002/cmdc.202200686.

(31) Wassarman, D. R.; Bankapalli, K.; Pallanck, L. J.; Shokat, K. M. Tissue-Restricted Inhibition of MTOR Using Chemical Genetics. Proceedings of the National Academy of Sciences 2022, 119 (38). 10.1073/pnas.2204083119.

(32) Bricogne G.; Blanc, E.; Brandl, M.; Flensburg, C.; Keller, P.; Paciorek, W.; Roversi, P.; Sharff, A.; Smart, O. S.; Vonrhein C.; Womack T.O. BUSTER. Global Phasing Ltd.: Cambridge, United Kingdom 2017.

(33) Emsley, P.; Lohkamp, B.; Scott, W. G.; Cowtan, K. Features and Development of Coot. Acta Crystallogr D Biol Crystallogr 2010, 66 (Pt 4), 486–501. 10.1107/S0907444910007493.

(34) Adams, P. D.; Afonine, P. V; Bunkoczi, G.; Chen, V. B.; Davis, I. W.; Echols, N.; Headd, J. J.; Hung, L. W.; Kapral, G. J.; Grosse-Kunstleve, R. W.; McCoy, A. J.; Moriarty, N. W.; Oeffner, R.; Read, R. J.; Richardson, D. C.; Richardson, J. S.; Terwilliger, T. C.; Zwart, P. H. PHENIX: A Comprehensive Python-Based System for Macromolecular Structure Solution. Acta Crystallogr D Biol Crystallogr 2010, 66 (Pt 2), 213–221. 10.1107/S0907444909052925.

(35) Laskowski, R. A.; MacArthur, M. W.; Moss, D. S.; Thornton, J. M. PROCHECK: A Program to Check the Stereochemical Quality of Protein Structures. J Appl Crystallogr 1993, 26 (2), 283–291. 10.1107/S0021889892009944.

(36) Vaguine, A. A.; Richelle, J.; Wodak, S. J. *SFCHECK* : A Unified Set of Procedures for Evaluating the Quality of Macromolecular Structure-Factor Data and Their Agreement with the Atomic Model. Acta Crystallogr D Biol Crystallogr 1999, 55 (1), 191–205. 10.1107/S0907444998006684.

